# Discovery, Structure, and Function of Filamentous 3-Methylcrotonyl-CoA Carboxylase

**DOI:** 10.1101/2022.08.19.504621

**Authors:** Jason J. Hu, Jane K.J. Lee, Yun-Tao Liu, Clinton Yu, Lan Huang, Inna Aphasizheva, Ruslan Aphasizhev, Z. Hong Zhou

## Abstract

3-methylcrotonyl-CoA carboxylase (MCC) is a biotin-dependent enzyme necessary for leucine catabolism in most organisms. While the crystal structure of recombinant bacterial MCC has been characterized, the structure and potential polymerization of native MCC remain elusive. Here, we discovered that native MCC from *Leishmania tarentolae* (LtMCC) forms filaments and determined its structure at near-atomic resolution using cryoEM. α_6_β_6_ LtMCC dodecamers assemble in a twisted-stacks architecture, manifesting as supramolecular rods extending up to approximately 400 nanometers. LtMCCs in filaments bind biotin but are not covalently biotinylated and lack coenzyme A. Filaments elongate by stacking α_6_β_6_ LtMCCs onto the exterior α-trimer of the terminal α_6_β_6_ dodecamer. This stacking immobilizes the biotin carboxylase domains, sequestering the enzyme in an inactive state within the mitochondrial matrix. Our results support a new model for LtMCC catalysis, termed the dual-swinging-domains model, and cast new light on the functional significance of polymerization in the carboxylase superfamily and beyond.

## INTRODUCTION

3-methylcrotonyl-CoA carboxylase (MCC) is a biotin-dependent enzyme necessary for the catabolism of leucine (Anderson et al., 1998; Knappe et al., 1961; Nikolau et al., 2003), an essential branched-chain amino acid involved in regulating cellular metabolism (Gondáš et al., 2022; Yang et al., 2010), protein synthesis (Gondáš et al., 2022; Yang et al., 2010), and anabolic signaling (Gran and Cameron-Smith, 2011; Paulussen et al., 2021). MCC belongs to a superfamily of biotin-dependent carboxylases with different substrate preferences, such as acetyl-CoA carboxylase (ACC), geranyl-CoA carboxylase (GCC), propionyl-CoA carboxylase (PCC), and pyruvate carboxylase (PC). In eukaryotes, MCC resides in the mitochondrial matrix (Tong, 2013). Documented in many species (Diez et al., 1994; Gallardo et al., 2001; Höschle et al., 2005), MCC shares high cross-species sequence homology (Figure S1). In humans, upregulated MCC expression often correlates with various cancers (Chen et al., 2021; Dai et al., 2020; Gondáš et al., 2022; He et al., 2020; Liu et al., 2019). On the other hand, MCC deficiency, one of the most common metabolic disorders in newborns (Lee and Hong, 2014), may cause vomiting, seizures, and other neurological abnormalities (Grünert et al., 2012; Kim et al., 2017).

As part of the mitochondrial leucine degradation pathway, MCC catalyzes the conversion of 3-methylcrotonyl-CoA to 3-methylglutaconyl-CoA (Gallardo et al., 2001). Upon covalent biotinylation of the apoenzyme, MCC accelerates two successive reactions: biotin carboxylation and carboxyl group transfer (Lynen et al., 1961; Moss and Lane, 1971). First, biotin is carboxylated at an α-subunit active site with bicarbonate as the carbon dioxide donor upon concomitant ATP hydrolysis (Lynen et al., 1961; Moss and Lane, 1971; Tong, 2013). Next, the carboxylated biotin is translocated to the corresponding β-subunit active site, where the carboxyl group is transferred from biotin to 3-methylcrotonyl-CoA (Lynen et al., 1961; Moss and Lane, 1971; Tong, 2013).

The atomic structure of bacterial MCC has been solved by X-ray crystallography using recombinant *Pseudomonas aeruginosa* MCC (PaMCC^rec^) expressed in *Escherichia coli* with His-tags to the β-subunits (Huang et al., 2012). PaMCC^rec^ subunits oligomerize into a dodecameric complex with a core of six β-subunits sandwiched by two α-trimers in an α_6_β_6_ architecture (Huang et al., 2012). Whether MCCs may exist in other forms is unclear. Albeit, their supramolecular assembly was conjectured based on rod-shaped aggregations of *Achromobacter* IVS MCC observed by negative-stain electron microscopy (Apitz-Castro et al., 1970). Recent advances in cryogenic electron microscopy (cryoEM) have uncovered unexpected modes of enzyme polymerization and elucidated the regulatory roles of such architectural forms (Lynch et al., 2017; Mattei et al., 2020; Maud et al., 2022; Park and Horton, 2019; Wang et al., 2019; Webb et al., 2017). For example, near-atomic resolution cryoEM structures illuminated the regulatory functions of several filamentous forms of eukaryotic ACC (Hunkeler et al., 2018). In the absence of high-resolution structures of the native MCC enzyme, it remains unsettled whether native MCCs can similarly form supramolecular assemblies.

Here, we used a bottom-up structural proteomics cryoEM approach (Ho et al., 2020; Pfab et al., 2021) to identify and determine the near-atomic resolution structure of native *Leishmania tarentolae* MCC (LtMCC) enriched from mitochondrial fraction by streptavidin affinity pulldown. We discovered that LtMCC filaments consist of stacks of LtMCC α_6_β_6_ dodecamers. The biotin carboxylase (BC) domains of adjacent α_6_β_6_ dodecamers bind with each other, inhibiting their mobility. In contrast, at the filament termini, the BC domain is flexible, while the linker between the biotin carboxyl carrier protein (BCCP) and the BC-CT mediating (BT) domain is rigid. These observations support a dual-swinging-domains model for LtMCC catalysis and suggest a regulatory role of LtMCC filamentation in the mitochondrial matrix.

## RESULTS

### Discovery and identification of LtMCC filaments from mitochondrial lysate

While examining complexes enriched by sedimentation of mitochondrial lysate from parasitic protist *L. tarentolae* in 10%-30% glycerol gradient (Figure 1A) and affinity pulldown with streptavidin-coated magnetic beads, we serendipitously discovered filamentous structures (Figure 1B). We identified the constituents of these filamentous structures by following the workflow detailed in Figures 1B–1C, which is based on the cryoID bottom-up structural proteomics approach (Ho et al., 2020). Using single-particle reconstruction, we obtained two differently centered cryoEM maps of the filament middle segments, one at 3.4 Å and another at 3.9 Å resolution (Figure 1B). Additionally, we processed the termini particles and found that the two termini have the same structure, which was resolved at 7.3 Å resolution (Figure 1B).

**Figure 1.**
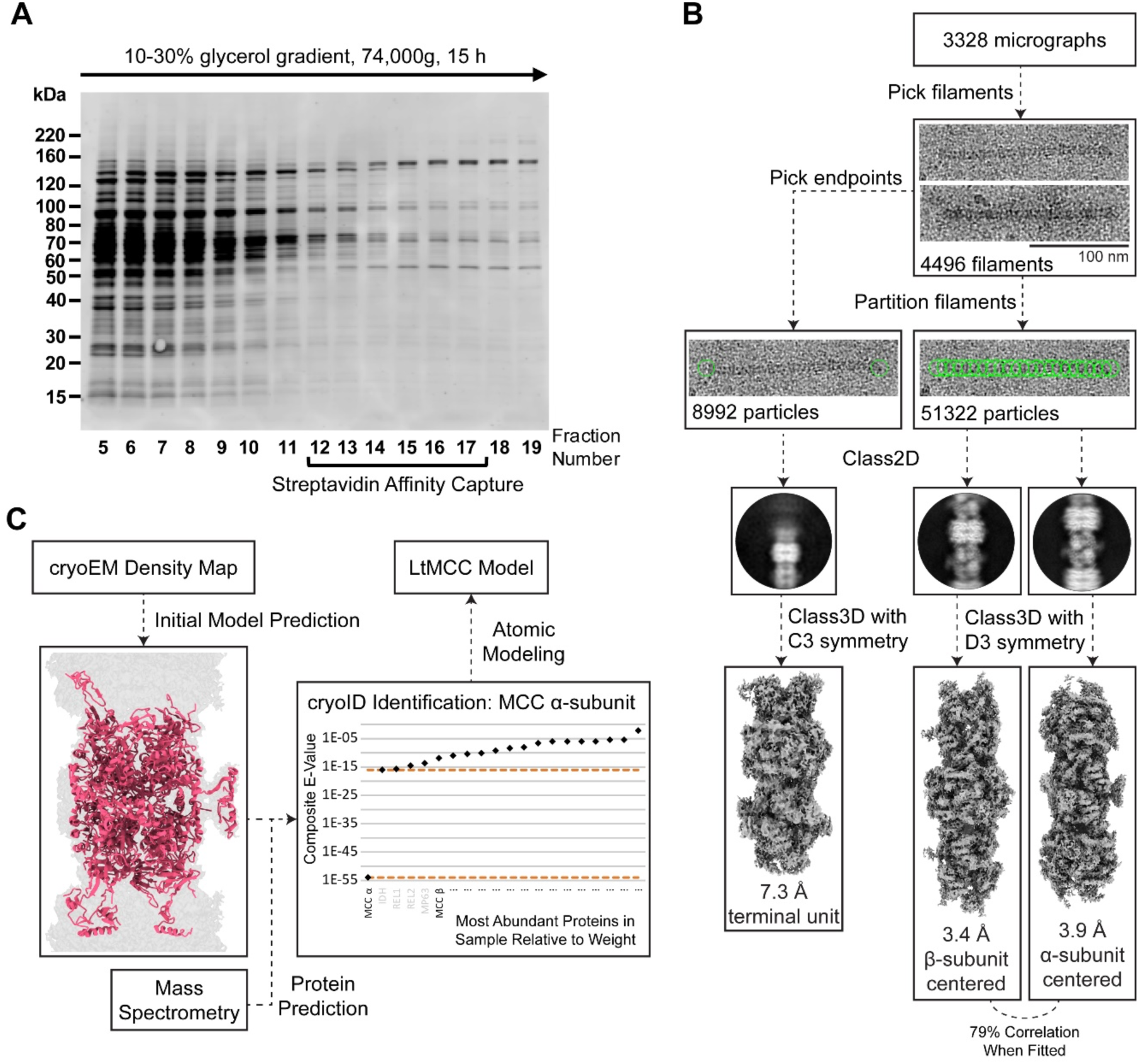
Discovery of filamentous 3-methylcrotonyl-CoA carboxylase from mitochondria of *L. tarentolae*. (A) Sodium dodecyl sulfate–polyacrylamide gel electrophoresis (SDS-PAGE) of the glycerol gradient fractions obtained from mitochondrial lysate. The gel was stained by Sypro Ruby. (B) Workflow for cryoEM imaging and reconstruction. Filaments observed in cryoEM micrographs were visually identified, partitioned into segments of equal length, classified in 2D and 3D. Refinement of 3D reconstructions yielded an α-centered and β-centered cryoEM map. The filament ends were picked and reconstructed similarly, converging into a single map for both ends of the filament (i.e., the termini map). (C) Workflow for protein identification and atomic modeling. From an input cryoEM map, DeepTracer (Pfab et al., 2021) generated an initial model with predicted amino acid sequence, which was input into cryoID (Ho et al., 2020) together with mass spectrometry data. CryoID predicted that the filaments contained LtMCC. Finally, de novo atomic modeling of LtMCC was performed in Coot (Emsley et al., 2010) using an AlphaFold predicted model as a reference (Jumper et al., 2021; Mirdita et al., 2022).

**Table 1.**
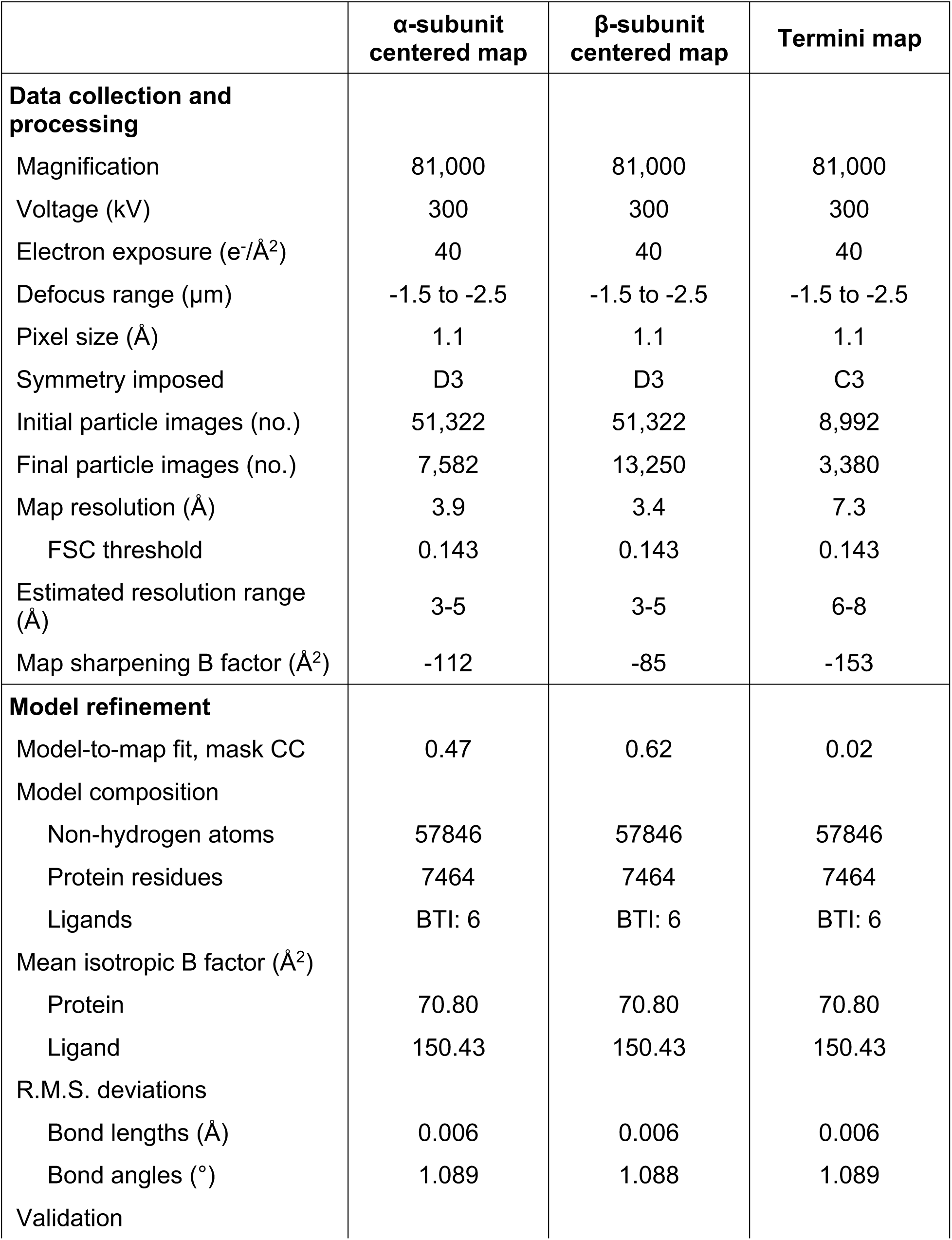
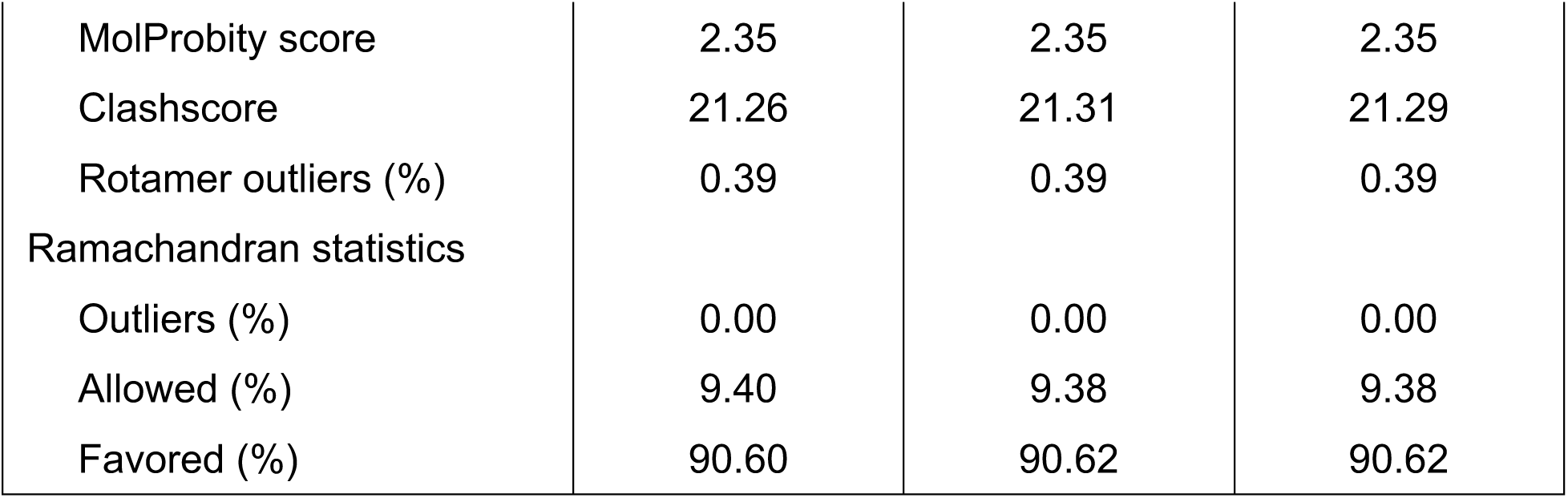
CryoEM map and model statistics.

To establish the protein identities of the cryoEM maps, we first obtained a partial model prediction from DeepTracer (Pfab et al., 2021) using the 3.9 Å map. The sequence of the output model from DeepTracer (Pfab et al., 2021) was then input into cryoID (Figure 1C). Searching for proteins detected by mass spectrometry (Table S1), cryoID identified LtMCC α-subunits as a filament component. Upon visual inspection, it was clear that the filaments also contain LtMCC β-subunits. Indeed, LtMCC α- and β- subunits produced the most peptide spectral counts of all proteins detected by mass spectrometry (Table S1). Furthermore, we ruled out GCCs as the filament components because GCCs are not found in eukaryotes (Waldrop et al., 2012). We built the atomic model for the filament *de novo*, using an AlphaFold (Jumper et al., 2021; Mirdita et al., 2022) predicted model for the α-subunit and the β-subunit as references. We found that both the 3.4 Å and 3.9 Å maps contain the same subunits but center on the β- and α- subunits, respectively. After translating one map for half of an α_6_β_6_ stack along the filament and a slight rotation around the filament axis, the two maps match well with one another (cross-correlation coefficient of 0.74). The slightly better resolution of the map centered on the β-subunits is probably because the architecture of the β-subunits is more stable.

### Structure of native eukaryotic LtMCC filaments

LtMCC filaments are predominantly straight (Figure 2A) although some are curved (Figure 2B), with a maximum observed curvature reaching 3.1 μm^-1^. With a mean length of 993 Å, the length distribution resembles a Poisson distribution (Figure 2C), similar to that of ACC filaments isolated from animal tissues and observed by negative stain electron microscopy (Kleinschmidt et al., 1969). Most filaments consist of four to six α_6_β_6_ dodecamers, or stacks (Figure 2C). With an architecture similar to that of PaMCC^rec^, each α_6_β_6_ stack in the LtMCC filament is characterized by an α_6_β_6_ cylindrical dodecamer exhibiting D3 symmetry (Figure 2D). Measuring to a height of 216 Å and width of 148 Å, the dodecamer is composed of a core of six β-subunits (β-core) sandwiched by two layers of trimeric α-subunits (Figure 2D, Video S1). Modeling the atomic structures of LtMCC α_6_β_6_ stacks into our cryoEM reconstruction reveals that the filament has a twisted-stacks architecture; that is, neighboring α_6_β_6_ stacks are related by a rotation of 23° and a translation of 216 Å (Figure 2D). In the filament, the active sites are exposed in both α- and β-subunits (indicated by stars in the middle panel of Figure 2D).

**Figure 2.**
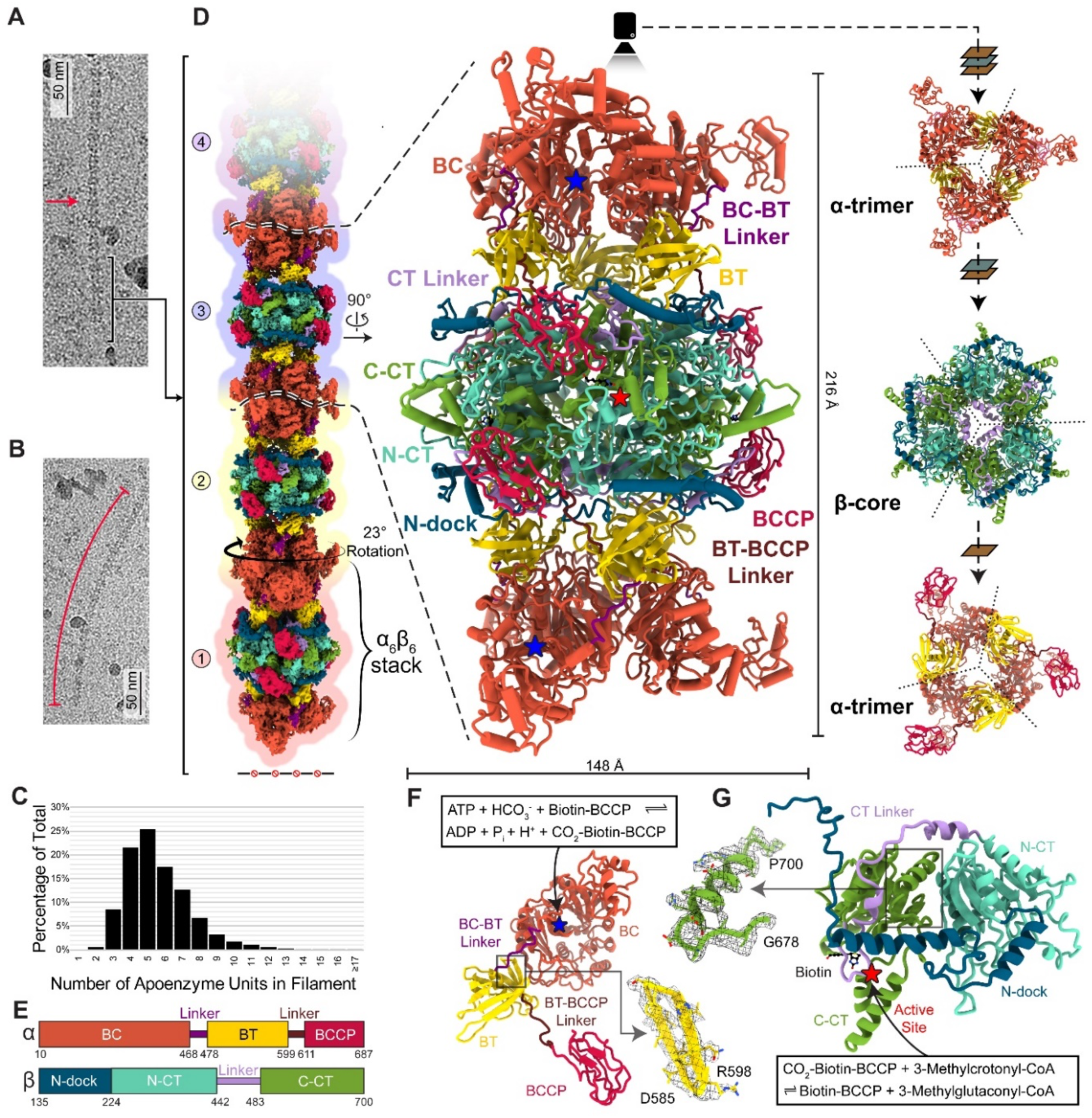
Structure of the filamentous LtMCC. (A-C) CryoEM images of a straight (A) and curved (B) LtMCC filament, and distribution of filament lengths (C). (D-E) CryoEM map and atomic models of the LtMCC filament. The composite cryoEM map of the filament (left column in D) is composed of three copies of the α-centered map, three copies of the β-centered map, and one copy of the termini map, with overlapping subunits. The atomic models (middle and right columns in D) of an α_6_β_6_ dodecamer and its trimeric stacks are shown in various views as ribbons, colored by domains according to the scheme in (E). The view directions are indicated by arrows, planes, and the camera. The dashed lines in the right column demarcate the boundaries between neighboring subunits (β-core is two-layered). (F-G) Atomic model of an α-subunit (F) and a β-subunit (G) of LtMCC shown as ribbons colored by domains with active sites labeled. Insets: example cryoEM densities superimposed with the atomic model.

Each α-subunit contains the BC, BT, and BCCP domains, and two linkers: the BC-BT linker and the BT-BCCP linker, the latter of which was unmodeled in PaMCC^rec^ (Figures 2E). A prominent feature of the BC domain is the presence of two large central β-sheets surrounded by α-helices. The active site of each α-subunit is located inside the central pocket of the BC domain (Figure 2F) and is responsible for carboxylating BCCP-bound biotin. In contrast to the BT domain of PaMCC^rec^, which contains a central α-helix accompanied by a seven-stranded antiparallel β-barrel (Huang et al., 2012), the LtMCC BT domain features eight β-strands (Figure S2). When the dodecameric PaMCC^rec^ and LtMCC are superimposed globally, the BC, BT, and BCCP domains in LtMCC are rotated with respect to those in PaMCC^rec^ (Figure S3A–B). However, when individual domains are superimposed independently, they fit well with each other (Figure S3C–E), suggesting that the above observed structural differences between the LtMCC filament and PaMCC^rec^ are likely due to filamentation.

Like in PaMCC^rec^, each LtMCC β-subunit possesses an N-carboxyltransferase (N-CT) domain and a C-carboxyltransferase (C-CT) domain. We have designated two new domains in LtMCC: the N-dock domain and CT-linker (Figure 2E), located before and after the N-CT domain, respectively. The N-dock domain was introduced to facilitate description of interactions between α- and β-subunits in the filament (see below). Designating the helix-loop-helix-loop fragment between the N-CT and C-CT domains as a separate domain (CT-linker) renders the N-CT and C-CT domains to have near-identical backbone folds (Figure S4), despite sharing only 19% sequence identity. Each of the six β-core active sites is wedged at the N-CT and C-CT interface between a top and a bottom β-subunit (Figures 2G, 3A) and is responsible for carboxylating 3-methylcrotonyl-CoA. Similar to that in PaMCC^rec^, each LtMCC β-core active site is roughly 80 Å away from its corresponding α-subunit active site.

**Figure 3.**
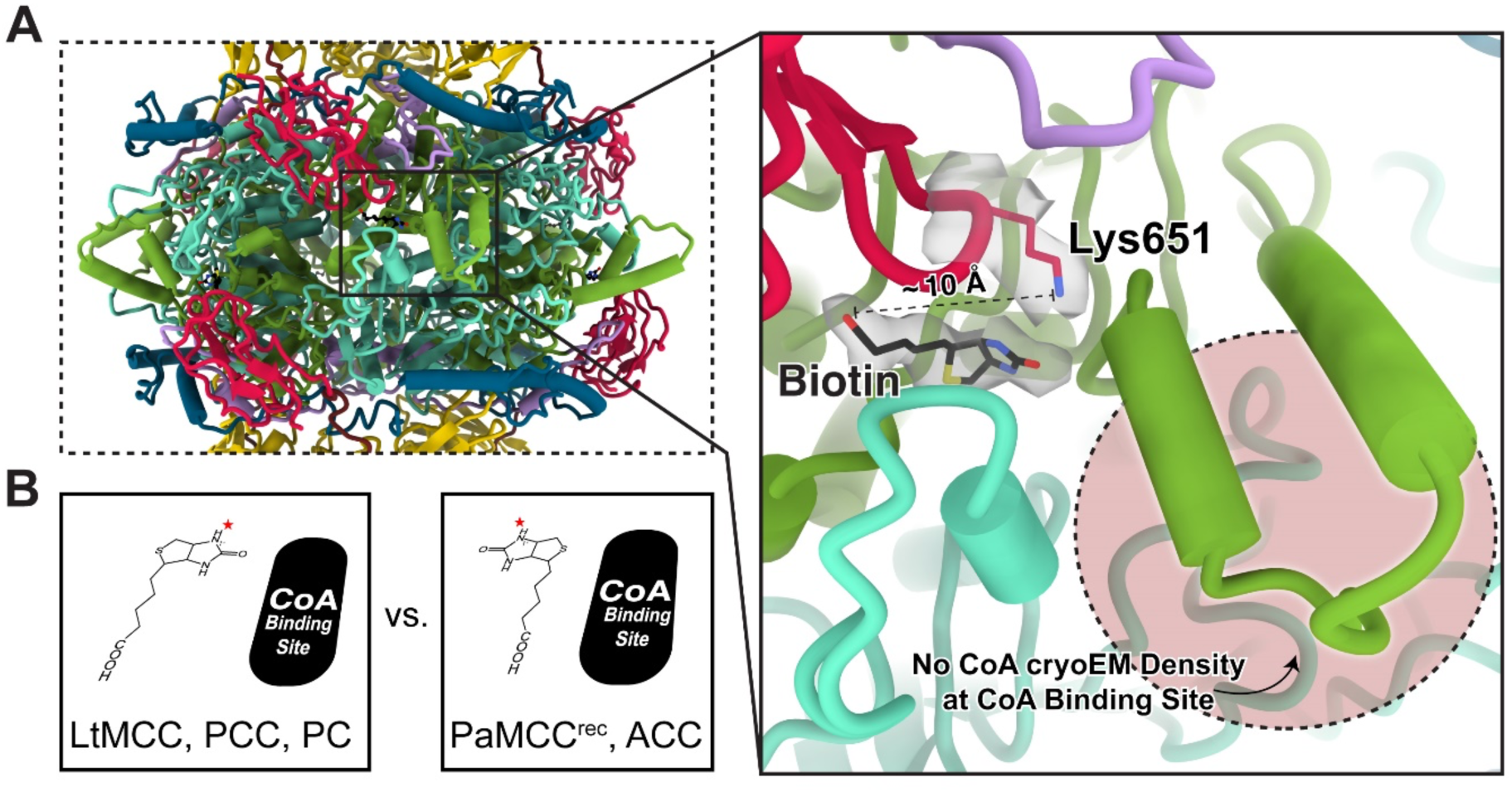
Presence of non-covalently bound biotin and lack of CoA in the LtMCC filament. (A) Close-up view of the middle column of Figure 2D, showing the biotin-containing site surrounded by the CT linker (purple), the BCCP (red), N-CT (turquoise), and C-CT (green) domains. The conserved Lys651 (red) and the non-covalently bound biotin (black) are shown as sticks, superimposed with their cryoEM densities (semi- transparent gray). The distance between the carboxyl group of biotin and the ε-amino group of Lys651 is about 10 Å. Notably, the CoA binding site (pink circle region) lacks cryoEM density, indicating absence of CoA. (B) Comparison of biotin and CoA orientations documented in various carboxylases (Fry et al., 1985; Huang et al., 2010, 2012; Scheffen et al., 2021; Wei and Tong, 2015). The red star indicates the location of the N1′ atom of biotin.

### LtMCC filaments bind biotin but lack coenzyme A

PaMCC^rec^ was crystalized as both a holoenzyme (with bound CoA and covalently-linked biotin) and an apoenzyme (without CoA and biotin). By contrast, α_6_β_6_ stacks in the LtMCC filament contain non-covalently-bound biotin but lack CoA (Figure 3A). For LtMCC to perform its two-step catalytic process, covalent biotinylation must occur first. Although biotin is observed at the expected β-core active sites, it is untethered to Lys651 in the highly conserved Ala-Met-Lys-Met sequence in the BCCP domain.

Indeed, the carboxyl group of biotin is about 10 Å apart from the ε-amino group of Lys651 (Figure 3A), which is too far to form a prosthetic group (Moss and Lane, 1971). The existence of non-biotinylated BCCP domains in LtMCC filaments suggests that the filamentous LtMCC is inactive. Similarly, inactive forms of PCC also bind biotin without forming a covalent adduct (Arabolaza et al., 2010; Diacovich et al., 2004).

The bicyclic ring system in biotin consists of an ureido ring and a thiophane ring (Hu and Cronan, 2020). Carboxylation requires the transfer of the carboxyl group attached to the N1′ atom of the biotin uredo ring to acyl-CoA (Knowles, 1989).

Therefore, the orientation of the N1′ atom of biotin is important for carboxylation. In LtMCCs, the N1′ atom in the ureido ring of biotin is proximal to the CoA binding site, but not in PaMCC^rec^ (Figure 3A). This is because the biotin bicyclic ring in LtMCC is approximately 180° rotated relative to that of PaMCC^rec^ (Figure 3B). The spatial arrangement of LtMCC biotin is consistent with PCC biotin (Huang et al., 2010; Scheffen et al., 2021) and PC biotin (Fry et al., 1985) (Figure 3B). In contrast, the arrangement of PaMCC^rec^ biotin is consistent with ACC biotin (Huang et al., 2012; Wei and Tong, 2015) (Figure 3B). Therefore, biotin orientation is variable at the β-core active site for different carboxylases and carboxylases in different catalytic states.

### α-subunits associate more with β-subunits than with intra-trimer α-subunits

Many interactions are observed between an α-subunit and the β-core, with 1,515 Å^2^ of buried surface area as calculated by ChimeraX (Goddard et al., 2018) and a binding affinity of -11.7 kcal mol^-1^ as calculated by PRODIGY (Vangone and Bonvin, 2015; Xue et al., 2016). Interactions between each α-subunit and the β-core are mainly mediated by a hook between the central α-helix and β1 strand of one BT domain that curls into a groove of a β-subunit (yellow segment from Asp494–Thr506 in Figure 4A). This groove is formed by the N-CT, C-CT, and N-dock domains (Figure S5). Due to its shorter sequence, the LtMCC hook contains only one hairpin in contrast to two hairpins in the PaMCC^rec^ hook (Figure S2). In both structures, the hook is crucial for interactions with the β-subunit to reinforce inter-subunit associations (Huang et al., 2012). The additional structures resolved in our cryoEM map also unveil previously unrecognized associations among domains of three neighboring subunits: the BT-BCCP linker of an α-subunit, N- dock domain of a β-subunit, and N-CT domain of another β-subunit (Figure 4B). Here, the BT-BCCP linker “docks” onto the crescent N-dock domain (Figure 4B).

**Figure 4.**
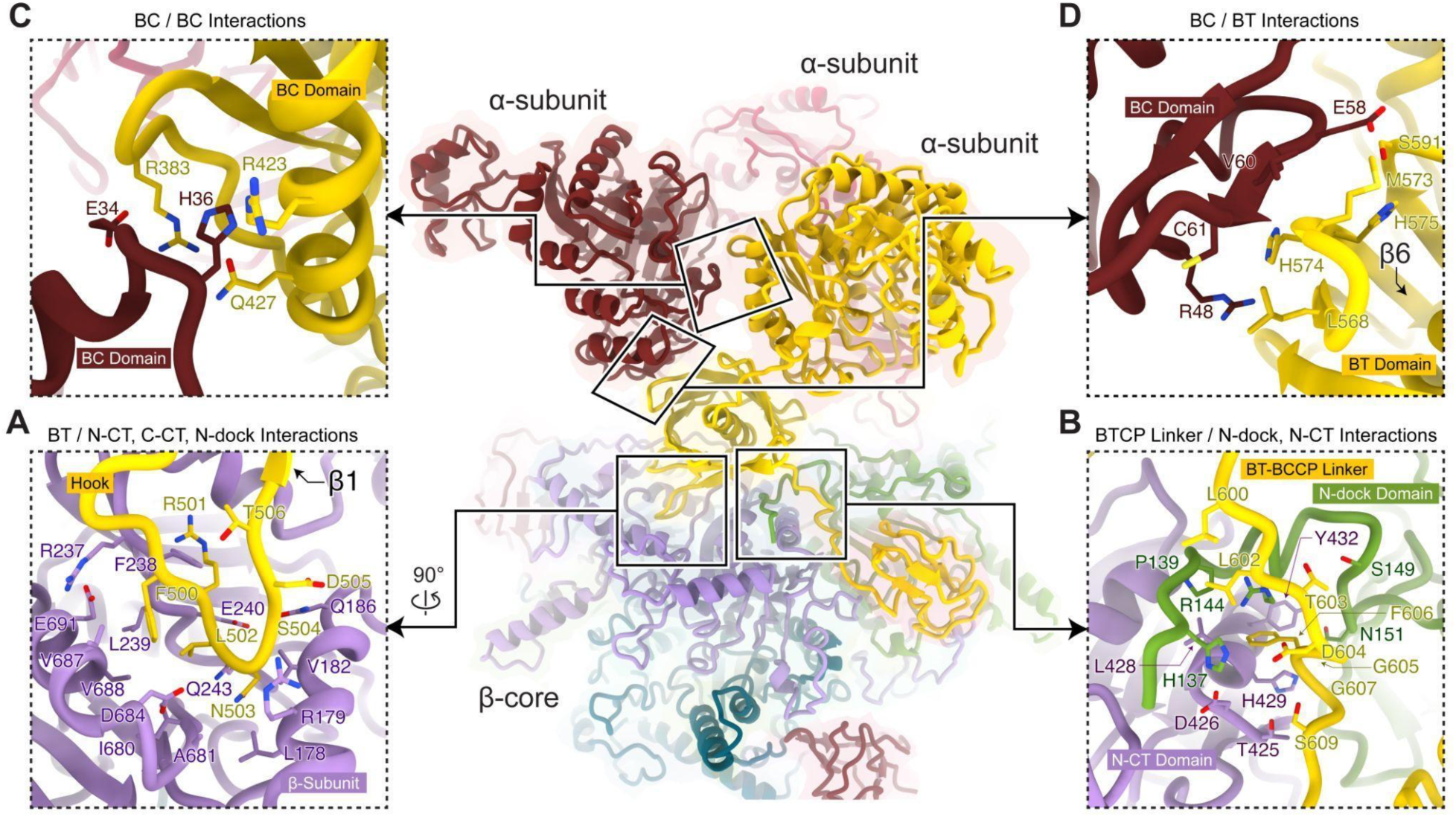
Subunit interactions in the filamentous LtMCC. Interacting regions are boxed and examined in close-up views (A-D) with their interfacial residues labeled. Interactions between α- and β-subunits (A-B) are more extensive than interactions among α-subunits (C-D).

Fewer interactions are observed between intra-trimer α-subunits, contributing just 579 Å^2^ of buried surface area and a binding affinity of -6.1 kcal mol^-1^. Within an α-trimer, each α-subunit interacts with its neighboring α-subunits through the BC-BC and BC-BT interfaces. At the BC-BC interface, a loop from one BC domain (Glu34–His36) interacts with a loop (Phe379–Pro385) and a helix (Arg423–Gln427) from a neighboring BC domain (Figure 4C). At the BC-BT interface, strand β6 (Gly572–Ala581) from the β- barrel in the BT domain interacts with a short β-strand (Ala59–Cys61) in the BC domain through β-sheet augmentation (Harrison, 1996) (Figure 4D).

### Structure of filament termini reveals mechanism of LtMCC polymerization

To investigate the mechanism of LtMCC polymerization, we reconstructed the filament termini at 7.3 Å resolution (Figure 5A). We found that both ends of the filament terminate at α-trimers, with partially resolved densities attributed to the BC domains (Figure 5A). When displayed at a high threshold, cryoEM densities for the BC, BCCP domain, and the BC-BT linker disappear, while density for the BT domain and BT-BCCP linker of the same α-subunit remain visible (Figure 5A). Therefore, the weaker densities of the BC and BCCP domains are due to their flexibility, rather than low occupancy of the α-subunit at the termini, which would make all domains of the α-subunit disappear.

**Figure 5.**
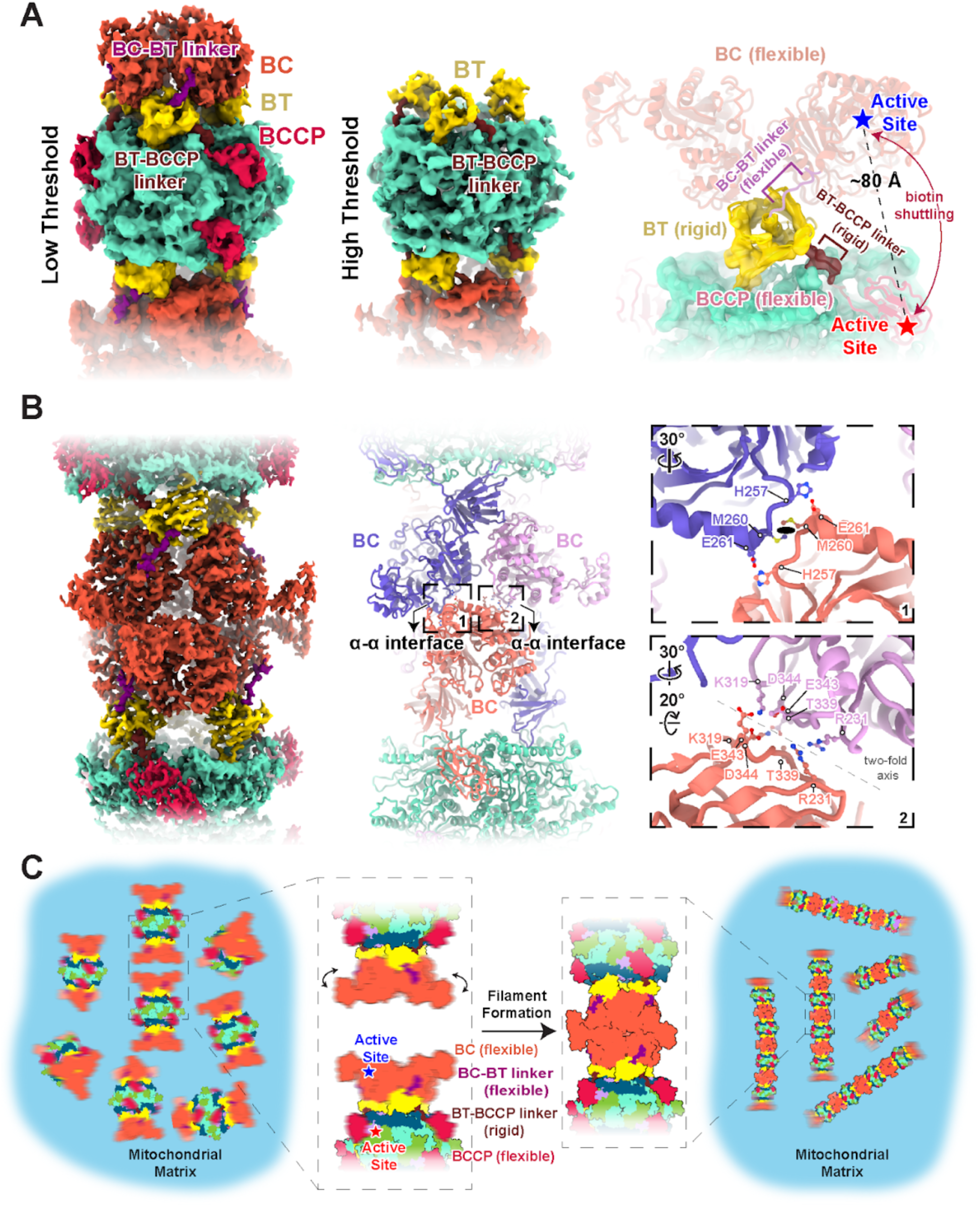
Filament formation stabilizes flexible BCCP and BT domains in LtMCC. (A) Structure of the filament termini. The left and middle panels show the 7.3 Å cryoEM map of the filament termini at a low and high density threshold, respectively. The α- subunit is colored by domain according to the scheme in Figure 2E and the entire β- core is turquoise. Note, while visible at low threshold, the BCCP and BC domain densities disappear at high threshold. The right panel shows the atomic model of the filament terminus with the rigid and flexible domains in harder and softer shades, respectively. (B) Structure of the α-α interface in the LtMCC filament. The cryoEM map (left panel) is colored as in (A) but the atomic models (middle panel) of the interfacing α-subunits are colored differently from each other to delineate individual α-subunits. Close-up views (right panel) of the two boxed areas in the middle panel show interfacing residues at the α-α interface. The ellipse in the top right panel denotes the local two-fold symmetry axis. (C) Proposed model of filament formation and stabilization. Blurry domains indicate their flexibility.

The flexibility of the BC and BCCP domains is further supported by limited intra-trimer α- subunit associations (Figures 4C–D). In contrast, the BT-BCCP linker is mostly rigid as it interacts extensively with the β-subunits (Figure 4B).

Our observation that LtMCC filaments terminate at α-subunits suggests that the building block for the filament is an α_6_β_6_ stack. Filaments plausibly elongate one stack at a time, attaching through BC domains to form an α-α interface (Video S2), which is defined as the associations between two α-trimers from adjacent stacks (Figure 5B).

The α-α interface has a total buried surface area of 1,689 Å^2^ and a binding affinity of -11.6 kcal mol^-1^. At the α-α interface, the BC domain of an α-subunit from the lower α- trimer interacts with the BC domains of two α-subunits from the upper α-trimer (Figure 5B). Interactions contributing to the α-α interface occur primarily between loops, along a local twofold axis (right panel of Figure 5B): the first site of interaction is between two loops (Leu338–Asp344), with its neighboring residues Lys319 and Arg231 also participating in α-α interactions; the second site of interaction occurs between two loops connected to two α-helices (His257–Glu261). These interactions at the α-α interface stabilize the BC domains between neighboring stacks in the LtMCC filament (Figure 5B).

## DISCUSSION

This study reveals an important, yet previously unrecognized role of the BC domain in MCC catalysis. During catalysis, the BCCP domain must shuttle biotin between the CT active site (β-subunit) and the BC active site (α-subunit), located approximately 80 Å apart (Figure 5A). However, we observed that the BT-BCCP linker is fixed (Figure 5A). Therefore, the maximum distance that the BCCP domain can translocate by itself is about 60 Å, which is insufficient for the BCCP domain to reach the BC active site (right panel of Figure 5A). As such, for catalysis to occur, both the BCCP and active-site-containing BC domain must move so that BCCP-bound biotin can reach the BC active site. Indeed, at the filament termini, the BC domain is flexible, thus permitting BC domain movement. We term this the dual-swinging-domains model to highlight the difference from the original swinging-domain model (Tong, 2013), in which only the BCCP domain swings. Beyond MCC, both the BCCP and BC domains have been documented to move in pyruvate carboxylase (Maurice et al., 2007; Xiang and Tong, 2008). These observations suggest that carboxylase catalysis in general may involve a dual-swinging domains mechanism.

In LtMCC filaments, the α-α interface fixes the BC domain, thus preventing its swinging. According to our dual-swinging-domains model, immobilization of the BC domain inhibits catalysis. Therefore, filamentation likely sequesters LtMCCs in a quiescent state in the mitochondrial matrix (Figure 5C), which may provide a capacity to reactivate the enzyme in response to environmental signals. Polymerization appears to be a readily deployable and economic mechanism for regulating the activity of abundant mitochondrial matrix enzymes in response to rapid changes in cellular metabolism requirements. Such capacity is particularly relevant for parasitic protists cycling between the amino acid-rich gut environment of insect vectors and the high glucose concentration in the bloodstream of their hosts.

Our inquiries into MCC polymerization further reflect the diversity of polymerization in the carboxylase superfamily. It is known that both isoforms of ACC, ACC1 and ACC2, form filaments (Beaty and Lane, 1983; Hunkeler et al., 2018; Kim et al., 2010; Kleinschmidt et al., 1969; Meredith and Lane, 1978; Park et al., 2013; Wei and Tong, 2018). Recently, high-resolution structures of filamentous ACC1 have been reported (Hunkeler et al., 2018). In contrast to LtMCC, ACC1 assembles into two filamentous forms: an active single-stranded and inactive double-stranded helical structure. Eukaryotic MCC and ACC filamentation exhibit different architectures and regulatory functions, supporting the distinct evolutionary lineages of these enzymes (Huang et al., 2012). Nonetheless, filament formation may be a general way to regulate enzymatic activity in the carboxylase superfamily.

Evolutionarily, employing the same gene costs less than “inventing” a new gene for regulation. Polymerization is a remarkable example of such a cost-effective strategy. In the case of LtMCC, the stacking of active α_6_β_6_ dodecamers can inhibit biotin shuttling (Figure 5C), thus sequestering enzymes by polymerization. In other enzymes, polymerization can alter specificity (Kim et al., 2005), increase activity (Polley et al., 2019), and even introduce non-enzymatic functions such as conducting electrons between bacterial cells (Wang et al., 2019) and regulating cell curvature (Ingerson-Mahar et al., 2010). Just like how complex life forms arose by the aggregation and polymerization of unicellular forms, the diverse functions arising from enzyme polymerization can be considered as a rudimentary form of evolution.

## STAR★METHODS

### KEY RESOURCES TABLE

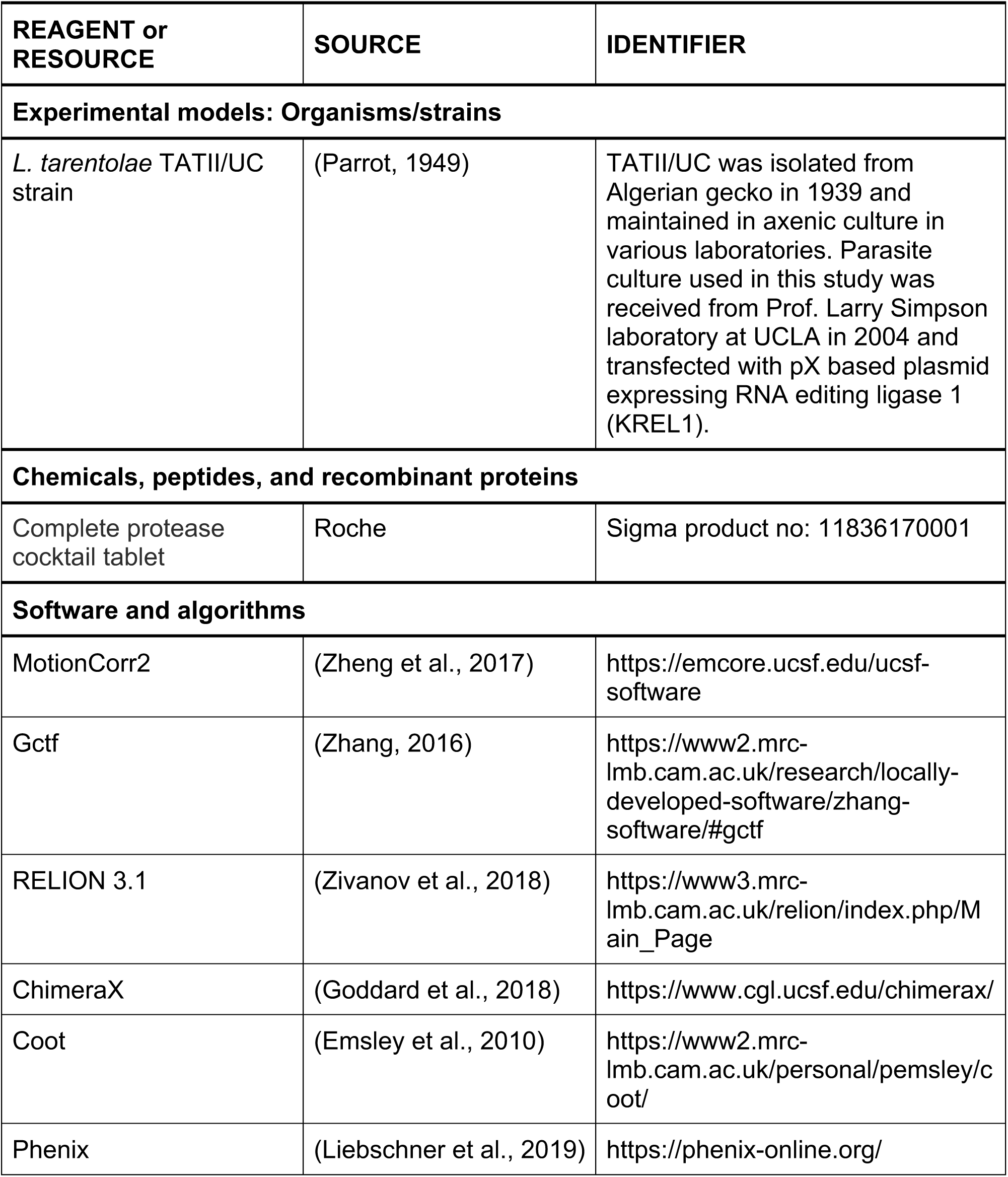
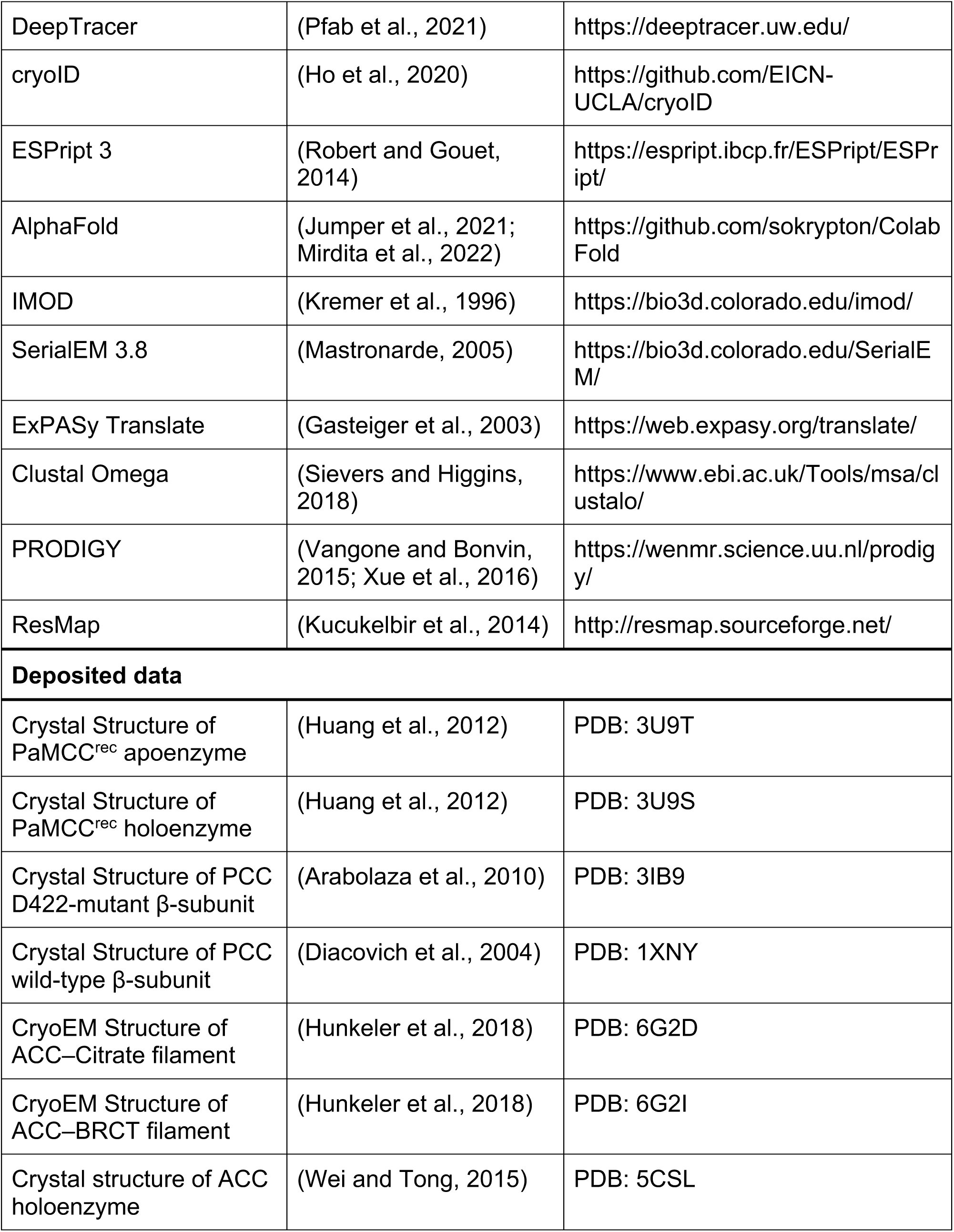
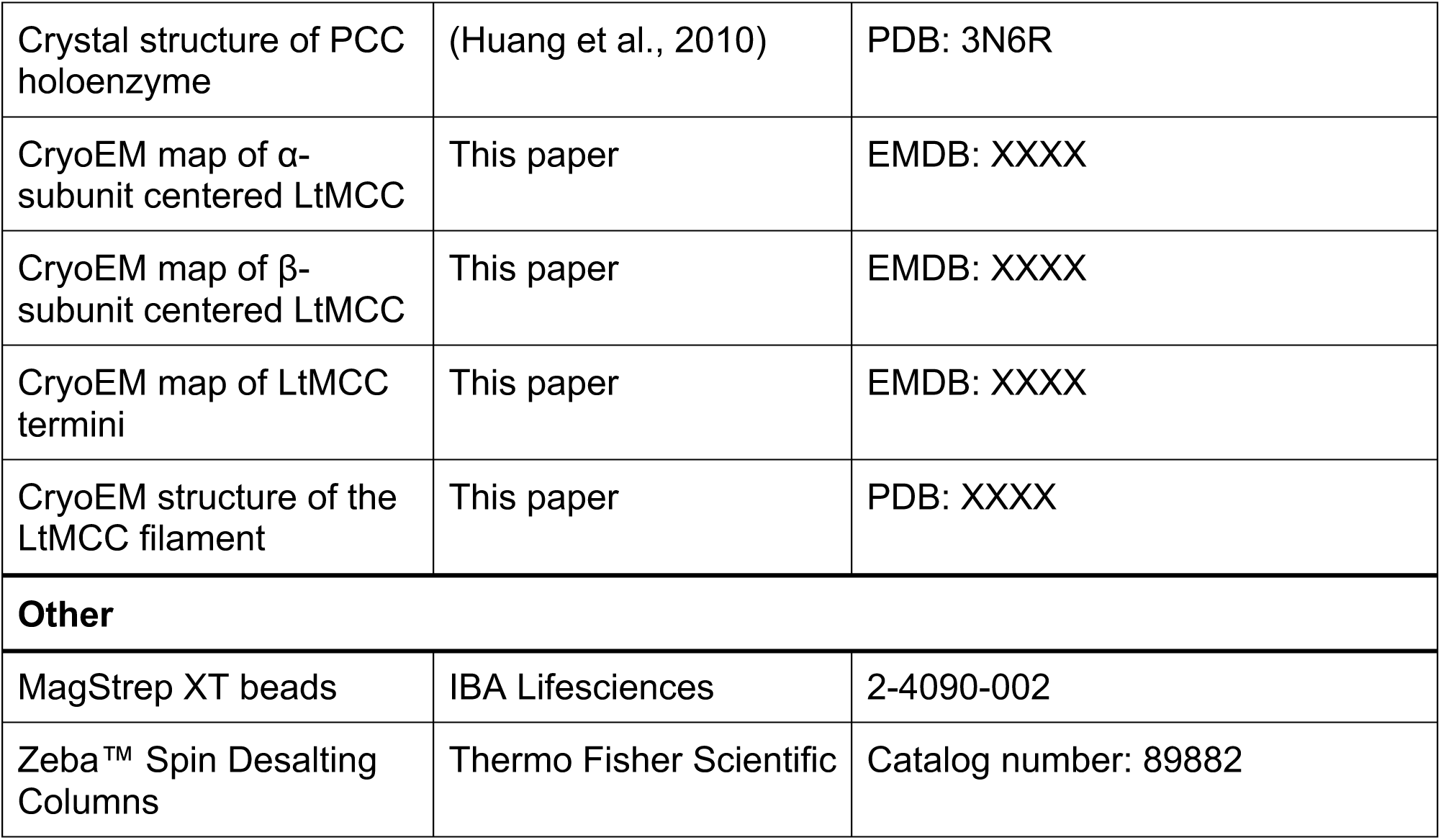

### RESOURCE AVAILABILITY

#### Lead contact

Further information and requests for resources and reagents should be directed to and will be fulfilled by the lead contact, Z. Hong Zhou.

### EXPERIMENTAL MODEL AND SUBJECT DETAILS

*L. tarentolae* cells were grown at 27°C in brain heart infusion media supplemented with 5 mg/L of hemin and harvested during late-exponential growth phase at ∼2x10^8^ cell/ml.

### METHOD DETAILS

#### Preparation of mitochondrial lysate

Mitochondrial fraction was enriched by hypotonic cell lysis and sequential separation of membrane-containing fraction on RenoCal76 density gradients (Pelletier et al., 2007). Mitochondrial pellets (0.5-0.8 g wet weight) were lysed in 1 ml of pH 7.3, 50 mM HEPES, 150 mM KCl, 2 mM EDTA, 1% NP40, and 50 μl of 20x Complete protease cocktail by sonication at 24W for 15 seconds and centrifuged at 30,000 RPM in an SW55 rotor for 15 minutes. The supernatant was recovered and separated on a continuous 10-30% gradient glycerol in pH 7.3, with 20 mM HEPES, 100 mM KCl, and 1 mM EDTA and prepared in SW28/32 Setton clear tubes for 15 hours at 72,000 g.

Glycerol gradient fractions of 1.5 ml were collected from the top and those corresponding to 20S-40S region were combined.

#### Purification of LtMCC by streptavidin affinity pulldown

Glycerol gradient fractions were supplemented octylglucoside to 2 mM and incubated on Strep-Tactin®XT magnetic beads in a Binding Buffer (50 mM Tris-HCl, pH 8.0, 100 mM KCl, 1 mM EDTA, 2 mM OG) for 1 hour at 4°C on a nutating mixer. Beads were washed with 5 ml of Binding Buffer twice. For elution, the beads were incubated in 0.2 ml of Elution Buffer (20 mM Tris-HCl, pH 8.0, 100 mM KCl, 1 mM EDTA, 2 mM OG, 100 mM biotin) at 4°C for 10 minutes. The 130 μl of purified material was exchanged into Sample Buffer (20 mM Tris-HCl, pH 7.5, 60 mM KCl, 5 mM MgCl_2_, 1 mM DTT, 5 mM OG) using Zeba™ Spin Desalting Columns, 7K MWCO (0.5 ml). The sample was then centrifuged at 21,000g for 10 minutes and the supernatant was stored on ice before grid preparation.

#### Protein identification by LC-MS/MS

Affinity-purified complexes were sequentially digested with LysC peptidase and trypsin. LC-MS/MS was carried out by nanoflow reversed phase liquid chromatography (RPLC) using an UltiMate 3000 RSLC (Thermo Fisher Scientific) coupled on-line to an Orbitrap Fusion Lumos mass spectrometer (Thermo Fisher Scientific). A cycle of full FT scan mass spectrum (m/z 375-1500, resolution of 60,000 at m/z 400) was followed by MS/MS spectra acquired in the linear ion trap for 3 seconds at top speed with normalized collision energy (HCD, 30%). Following data extraction to MGF format using MSConvert from ProteoWizard (Chambers et al., 2012), resultant peak lists for each LC-MS/MS experiment were submitted to Protein Prospector (UCSF) for database searching (Fang et al., 2012). Each project was searched against a normal form concatenated with the random form of the *L. tarentolae* Parrot Tar II from the TriTrypDB database (Amos et al., 2022). The mass accuracy for parent ions and fragment ions were set as ± 10 ppm and 0.6 Da, respectively. Trypsin was set as the enzyme, with a maximum of two missed cleavages allowed. Cysteine carbamidomethylation was set as a fixed modification, and protein N-terminal acetylation, methionine oxidation, and N-terminal conversion of glutamine to pyroglutamic acid were selected as variable modifications.

#### CryoEM sample preparation

Lacey carbon cryoEM grids with a 2 nm continuous carbon film (Ted Pella) were first glow discharged for 45 seconds with a target current of 15 mA, using PELCO easiGlowTM. Then a 2.5 μL sample was applied to the grids. After waiting for 5 seconds, the grids were blotted for 4 seconds with blot force 0, at 100% humidity and 4°C temperature; after blotting, the grids were plunge-frozen into liquid ethane using an FEI Mark IV Vitrobot (Thermo Fisher Scientific). The grids were stored in a liquid nitrogen dewar before imaging.

#### CryoEM image acquisition

The cryoEM grids were loaded and imaged using a Titan Krios (Thermo Fisher Scientific) operated at 300 kV, with a Gatan K3 camera and a Gatan Imaging Filter Quantum LS. Movies were recorded with SerialEM (Mastronarde, 2005) by electron counting in super-resolution mode at a pixel size of 0.55 Å/pixel. The exposure time was 2 seconds and fractionated to 40 frames. The defocus range was between -1.5 to -2.5 μm. The total dosage was approximately 40 electrons/Å2. A total of 3,328 movies were collected.

#### CryoEM image processing

The movies were processed with MotionCor2 (Zheng et al., 2017), leading to dose-weighted and drift-corrected electron micrographs with a calibrated pixel size of 1.1 Å. We discarded the first frame due to severe drift of this frame. The defocus of the micrographs was determined by Gctf (Zhang, 2016). Using IMOD (Kremer et al., 1996), the micrographs were binned by a factor of 10 to make a stack for particle picking. After directly observing filaments on the micrographs, we manually picked the start and end of each filament using the 3dmod command in IMOD (Kremer et al., 1996). A total of 4,496 filaments were picked. To derive the constituents of the filaments, we coded a Python script to subdivide the filaments into equally spaced segments of 87 Å. The coordinates were then converted through a Bash script to RELION format. A total of 51,322 particles were extracted in total through RELION 3.1, with a box size of 384.

After extraction, we performed reference-free 2D class averaging on the particles with RELION 3.1 (Zivanov et al., 2018). We chose the best classes and proceeded to 3D classification. We made an initial 3D model from the best 3D class averages through RELION 3.1. The 3D classification was done with D3 symmetry and yielded α- and β- centered 3D class averages. After removing duplicates, the α- and β-centered classes contained 7,582 and 13,250 particles, respectively. We did 3D refinement for the α- and β-centered map independently with D3 symmetry using RELION 3D refinement. We then performed CTF refinement for both maps independently using RELION 3.1. We used CTF refinement (Zivanov et al., 2018) to improve the accuracy of the defocus for each particle and to correct beam tilt for the dataset. After performing CTF refinement, we performed 3D refinement for both maps again, independently. At the end of all processing, based on RELION’s gold-standard Fourier shell correlation (FSC) at the 0.143 criterion, we obtained α- and β-centered maps at 3.9 and 3.4 Å resolution, respectively (Figure S6). The resolution towards the center of the map is better than the edge of the map (Figure S6).

For the filament termini map, we used RELION 3.1 to extract the previously picked filament start-and-end points at a box size of 384 pixels. We calculated the in-plane rotation angle using the direction of vectors from the start to end of the filament. We used a Python code to include the in-plane rotation angle information in the RELION star file for the particles. We performed 2D classification for these particles without local search around the precalculated in-plane rotation angle. To generate an initial 3D model, we used the IMOD clip function (Kremer et al., 1996) to change the center of the β-centered map. The best particles were selected for 3D reconstruction with C3 symmetry. We performed 3D classification and removed bad particles. During 3D classification, we set the “limit_tilt” parameter to 85 to keep side-view particles. We proceeded to 3D refinement with 3,380 particles. We obtained a final reconstruction with C3 symmetry at 7.3 Å resolution based on the gold-standard Fourier shell correlation at the 0.143 criterion.

#### Sequence determination

We re-sequenced the α-subunit gene with the same strain of *L. tarentolae* as used in cryoEM sample preparation. The amino acid sequence for the LtMCC β-subunit was obtained by analyzing the genome sequencing data of *L. tarentolae* Parrot Tar II (Goto et al., 2020), since the putative, shotgun LtMCC β-subunit sequence on the NCBI protein database was notably shorter than MCCs from other organisms, deviating from the high sequence homogeneity observed across MCC sequences. A frameshift occurs near the middle of the sequence, likely caused by a sequencing error. The frameshift is not present in the previous genome sequencing data of *L. tarentolae* Parrot Tar II (Raymond et al., 2012). Since MCC plays an essential role in the catabolism of leucine across many species, it is unlikely that MCC exhibits a highly different sequence in *L. tarentolae*.

#### Atomic modeling

An initial AlphaFold (Jumper et al., 2021) model prediction was generated from ColabFold (Mirdita et al., 2022) using the LtMCC α-subunit and β-subunit sequences. Atomic model building was conducted de novo in regions of low sequence conservation. First, an individual α-subunit and β-subunit were modeled into the cryoEM map in Coot (Emsley et al., 2010) and then real-space refined in Phenix (Liebschner et al., 2019).

The α-subunit was modeled using the α-centered map, and respectively for the β- subunit. Biotin was modeled de novo into its observed density. Then, the subunits were duplicated according to D3 symmetry, fitted into the cryoEM maps, and refined iteratively using Phenix (Liebschner et al., 2019) and Coot (Emsley et al., 2010).

#### Model analysis

Buried surface area and map cross-correlation coefficient were calculated in ChimeraX (Goddard et al., 2018). Binding affinity calculations were done using the PRODIGY web server (Vangone and Bonvin, 2015; Xue et al., 2016). Sequence alignments and similarity calculations were performed using Clustal Omega (Sievers and Higgins, 2018) from the EMBL-EBI web server and visualized with ESPript 3 (Robert and Gouet, 2014).

## Supporting information

PDB Validation Report

## Data and code availability

CryoEM maps of the α-subunit centered, β-subunit centered, and filament termini map have been deposited in the Electron Microscopy Data Bank under accession numbers EMD-XXXX, EMD-XXXX, and EMD-XXXX, respectively. The coordinates of LtMCC have been deposited in the Protein Data Bank under accession number XXXX. All aforementioned deposited data are publicly available as of the date of publication and accession numbers are also listed in the key resources table. This paper does not report original code. DOIs are listed in the key resources table. This paper analyzes existing, publicly available data. These accession numbers for the datasets are listed in the key resources table. Any additional information required to reanalyze the data reported in this paper is available from the lead contact upon request.

## Acknowledgments

Our research was supported in part by grants from the U.S. National Institutes of Health (R01GM071940 to Z.H.Z., R01AI101057 to R.A., and R01GM074830 to L.H.). J.J.H. acknowledges support from STROBE NSF Science & Technology Center Grant DMR- 1548924, as well as the Stone, Irving & Jean Award and the Rose Gilbert, in Memory of Maggie Gilbert Award granted by the UCLA Honors Programs. We acknowledge the use of resources at the Electron Imaging Center for Nanomachines of UCLA supported by U.S. NIH (S10RR23057 and S10OD018111) and U.S. NSF (DMR-1548924 and DBI-133813). We thank James Zhen for his advice in manuscript writing. We thank Shiqing Liao and Samuel Liu for their help with selecting curved filaments.

## Author contributions

Z.H.Z. and R.A. initiated and supervised the project. I.A. prepared the sample. C.Y. and L.H. conducted mass spectrometry. Y-T.L. carried out cryoEM imaging and data processing with assistance from J.K.J.L. and J.J.H. J.K.J.L. and J.J.H. built the atomic models. J.K.J.L., J.J.H., and Y-T.L. analyzed the data. J.J.H., J.K.J.L., and Y-T.L. made illustrations and wrote the original draft. Z.H.Z., Y-T.L., J.J.H., and J.K.J.L. finalized the manuscript. All authors reviewed and approved the paper.

## Declaration of interests

The authors declare no competing interests.

## SUPPLEMENTAL FIGURES

**Figure S1.**
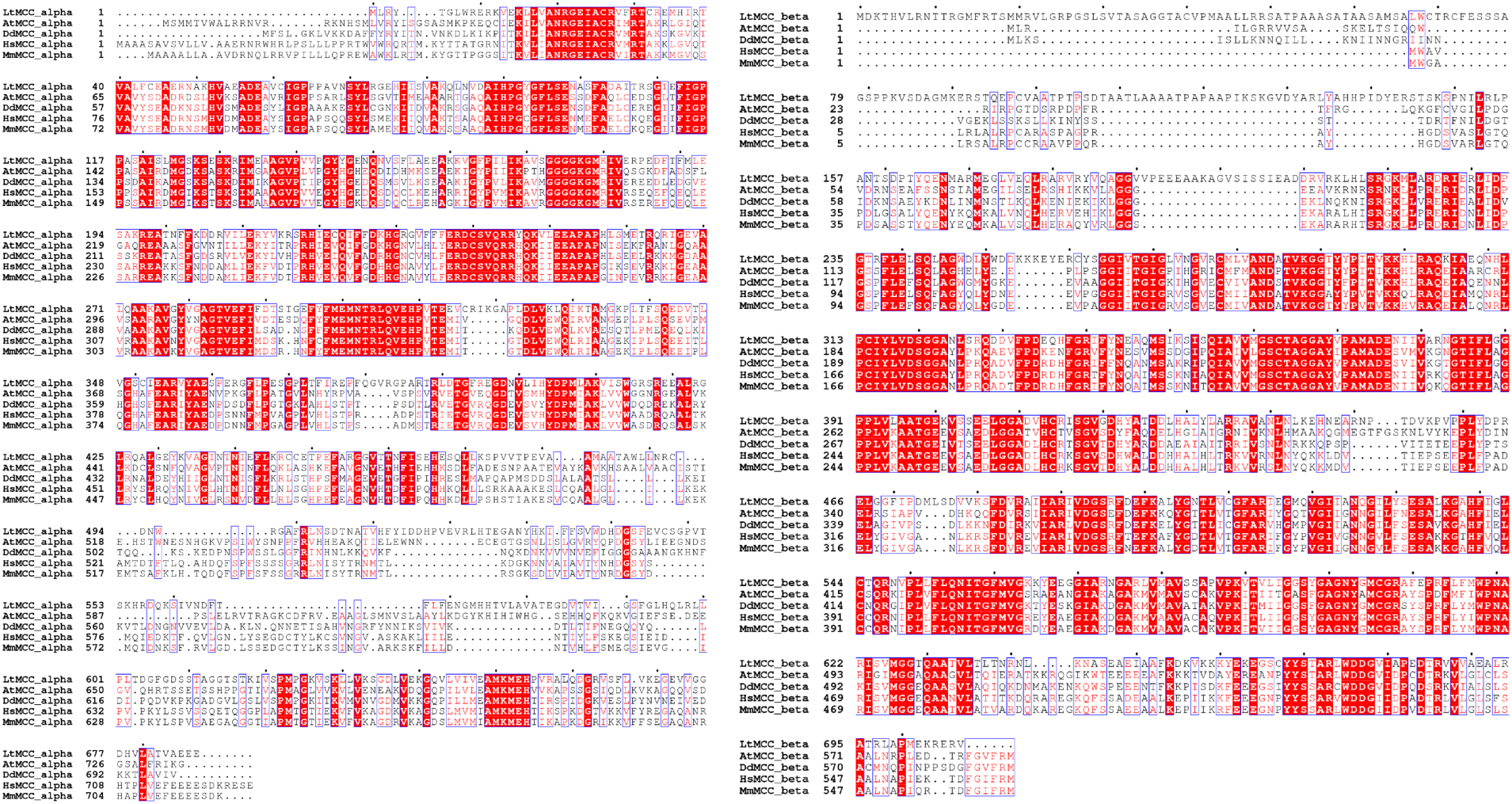
Multiple sequence alignment from *ESPript 3.0* (Robert and Gouet, 2014). Left panel compares LtMCC α-subunit with α-subunits from *Arabidopsis thaliana* MCC (AtMCC), *Dictyostelium discoideum* MCC (DdMCC), *Homo sapien* MCC (HsMCC), and *Mus Musculus* MCC (MmMCC). Right panel compares LtMCC β-subunit with β-subunits from AtMCC, DdMCC, HsMCC, and MmMCC. Boxes with and without solid red background indicate identical residues and similar residues, respectively. Dots on top of the residues demarcate intervals of ten residues.

**Figure S2.**
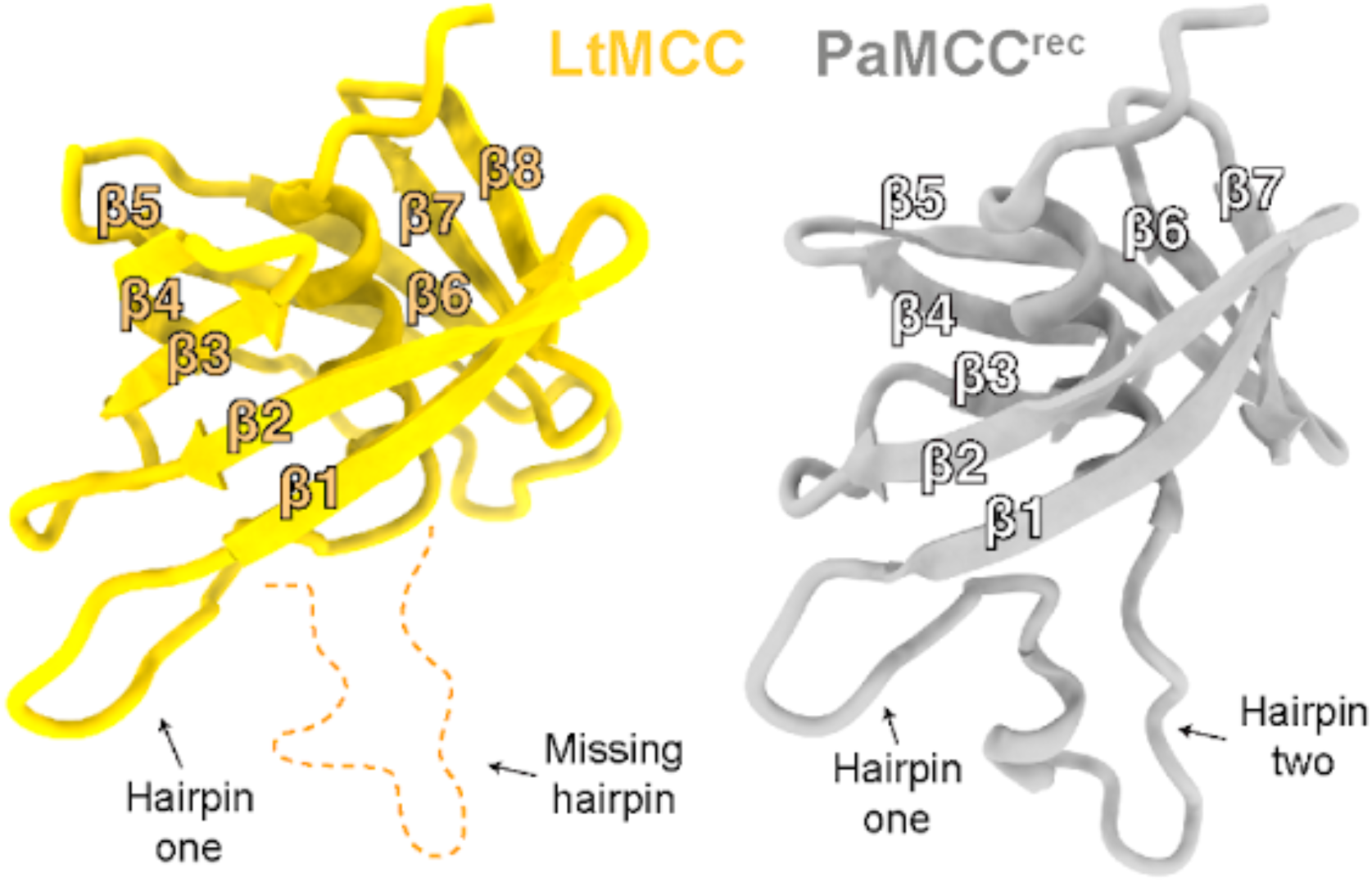
Structural comparison of the BT domain from LtMCC and PaMCC^rec^. The LtMCC BT domain exhibits an 8-stranded β-sheet fold while the PaMCC^rec^ BT domain contains a 7-stranded β-sheet fold. The PaMCC^rec^ BT domain possesses a hook with two hairpins. The LtMCC BT domain possesses a hook with only one hairpin.

**Figure S3.**
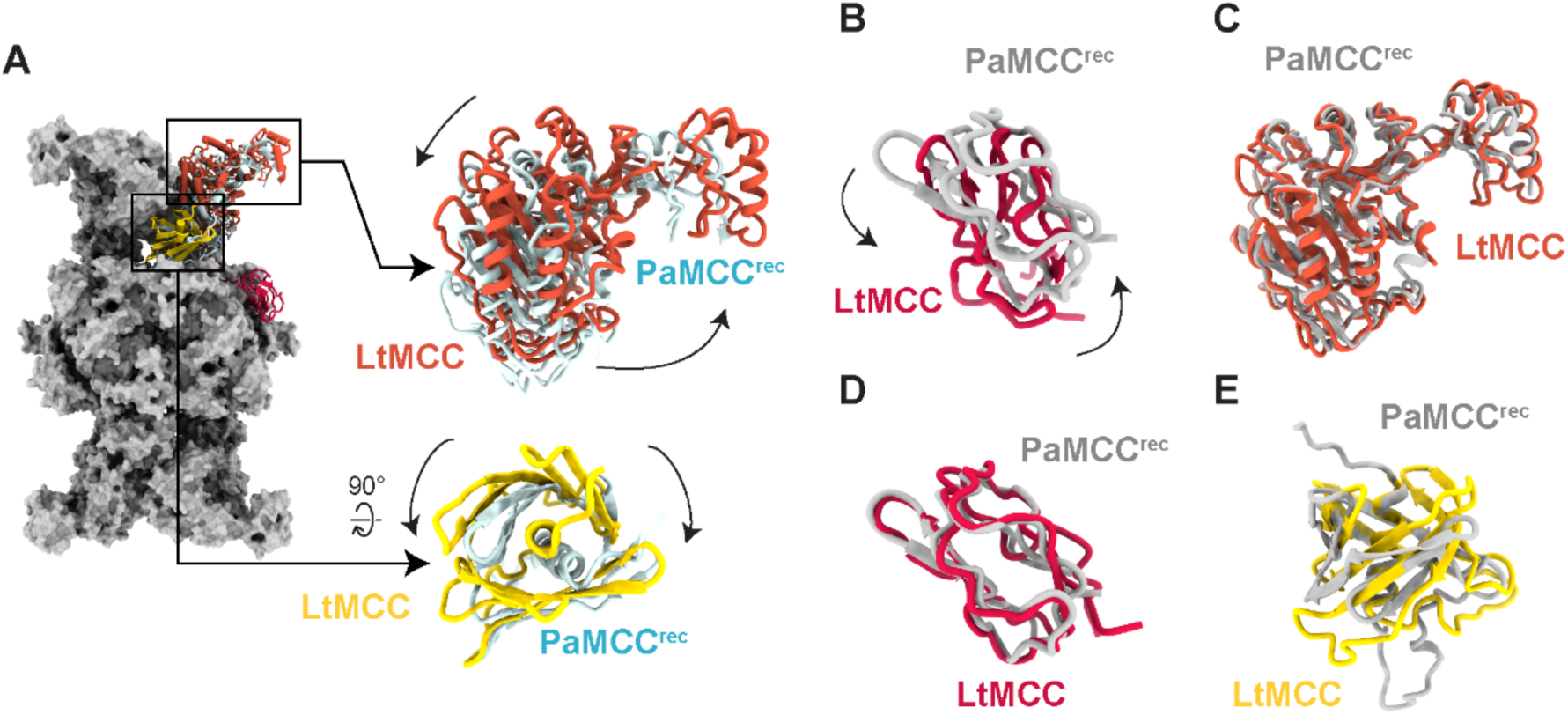
Structural comparison of PaMCC^rec^ and LtMCC. (A-B) Mismatched domain structures when the PaMCC^rec^ and LtMCC α_6_β_6_ dodecamers are superimposed globally. PaMCC^rec^ apoenzyme (A) and holoenzyme (B) are globally superimposed with the LtMCC dodecamer. In (A), the BC and BT domains are rotated in LtMCC compared to the PaMCC^rec^ apoenzyme, as indicated by arrows. The BCCP domain is unmodeled in the PaMCC^rec^ apoenzyme. The entire PaMCC^rec^ apoenzyme is shown as a gray surface. In (B), the BCCP domain in LtMCC is rotated relative to the BCCP domain in the PaMCC^rec^ holoenzyme, as indicated by arrows. (C-E) Matched domain structures when individual domains from PaMCC^rec^ and LtMCC are superimposed locally. BC (C), BCCP (D), and BT (E) domains from the two dodecamers are treated as individual models and superimposed independent from the global dodecameric structure.

**Figure S4.**
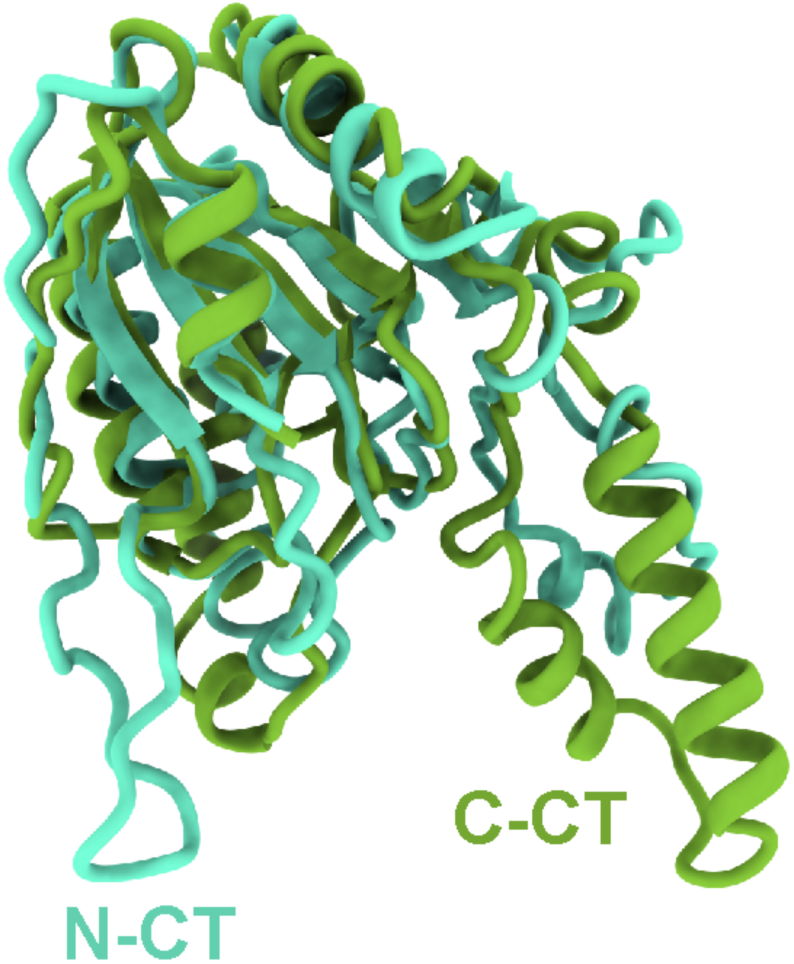
The N-CT and C-CT domain local alignment for LtMCC. The N-CT and C-CT domains are homologous, despite low sequence similarity.

**Figure S5.**
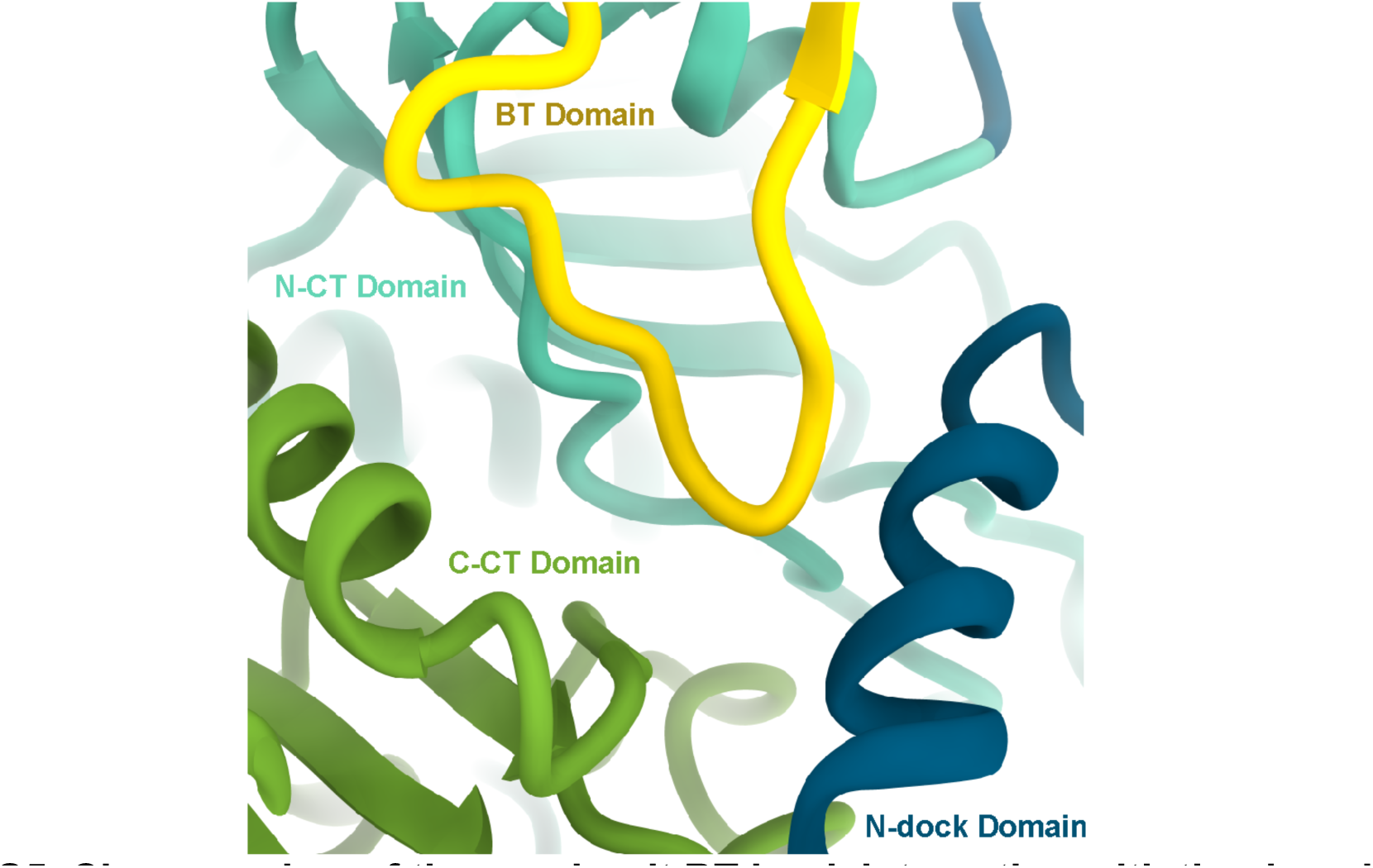
Close-up view of the α-subunit BT hook interacting with the domains that make up the β-subunit groove. The BT hook is colored yellow. The N-CT, C-CT, and N-dock domains from the same β- subunit make up the β-subunit groove. The N-CT domain is colored in light blue, the C- CT domain is colored in green, and the N-dock domain is colored in dark blue.

**Figure S6.**
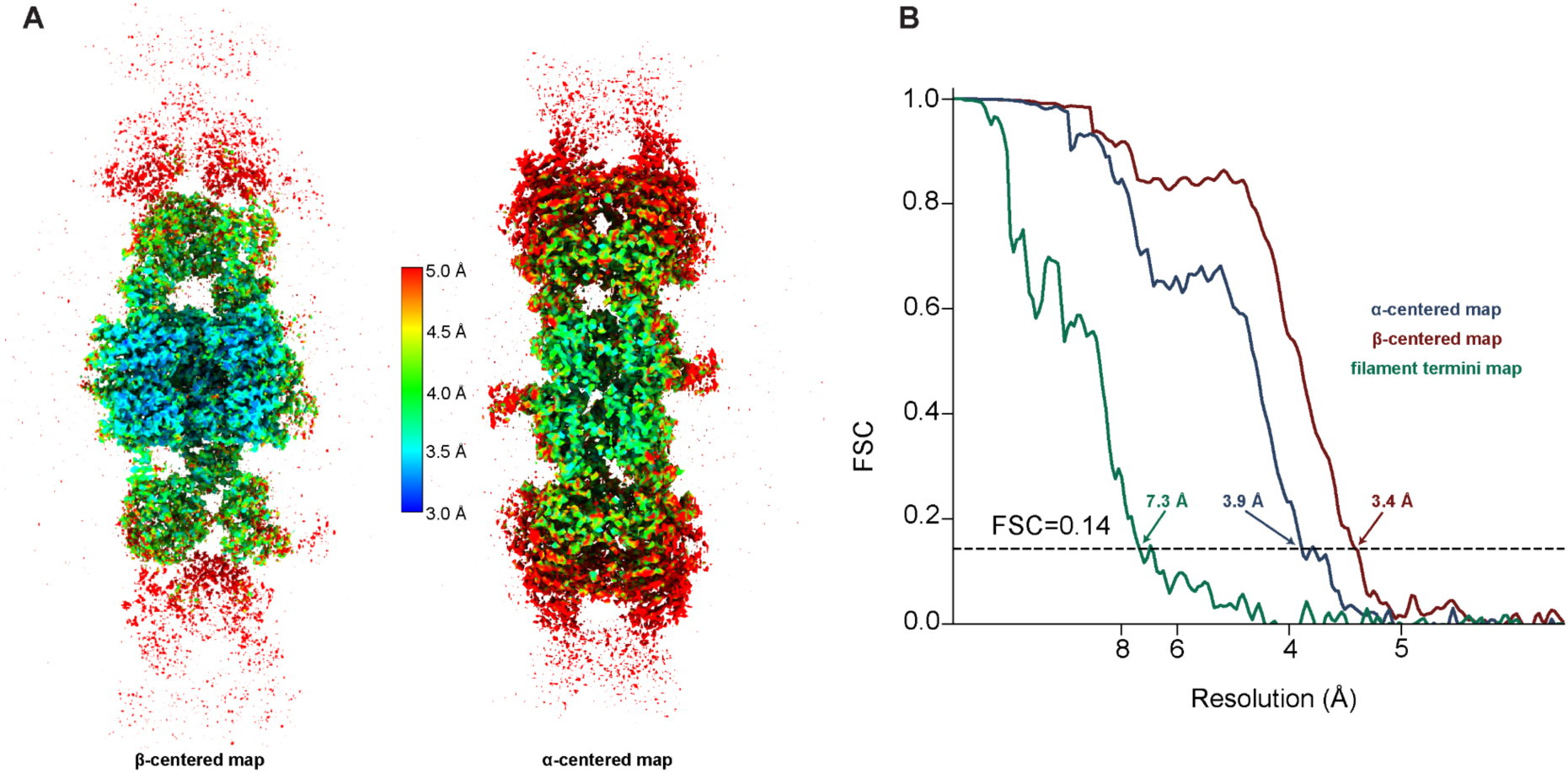
Resolution evaluation of cryoEM reconstruction for LtMCC filaments. (A) Local resolution evaluation by ResMap (Kucukelbir et al., 2014) for the α-centered and β-centered cryoEM maps of LtMCC filaments. The color key between the two maps shows their coloring by resolution. (B) FSC curves of the final α-centered, β-centered, and filament termini cryoEM maps.

## Supplemental Video Legends

Video S1. Ribbon diagram of an α_6_β_6_ dodecamer of the filamentous LtMCC, showing the α-trimer and β-core layers, related to Figure 2D.

Video S2. LtMCC filamentation stabilizes the BC domain, related to Figure 5.

## Supplemental Table Legends

Table S1. Mass spectrometry data with LtMCC subunits colored in red

## Notes

### Competing Interest Statement

The authors have declared no competing interest.

### Summary of Updates

Author name corrected; minor edits to manuscript and supplementary information

