## Supplementary material for "Discovery, Structure, and Function of Filamentous 3-Methylcrotonyl-CoA Carboxylase": PDB Validation Report

### Preliminary Full wwPDB EM Validation Report (i)

Aug 20, 2022 – 04:10 PM EDT

**This wwPDB validation report is NOT for manuscript review**

This is a Preliminary Full wwPDB EM Validation Report.

This report is produced by the standalone wwPDB validation server.
**The structure in question has not been deposited to the wwPDB.**
**This report should not be submitted to journals.**

We welcome your comments at **
A user guide is available at
https://www.wwpdb.org/validation/2017/EMValidationReportHelp
with specific help available everywhere you see the (i) symbol.

The types of validation reports are described at http://www.wwpdb.org/validation/2017/FAQs#types.

Not For Manuscript Review

The following versions of software and data (see references (i)) were used in the production of this report:

| | | |
|---|---|---|
| MolProbity | : | 4.02b-467 |
| Mogul | : | 1.8.5 (274361), CSD as541be (2020) |
| Percentile statistics | : | 20191225.v01 (using entries in the PDB archive December 25th 2019) |
| Ideal geometry (proteins) | : | Engh & Huber (2001) |
| Ideal geometry (DNA, RNA) | : | Parkinson et al. (1996) |
| Validation Pipeline (wwPDB-VP) | : | 2.30 |

### 1 Overall quality at a glance (i)

The following experimental techniques were used to determine the structure:
*ELECTRON MICROSCOPY*

The reported resolution of this entry is unknown.

Percentile scores (ranging between 0-100) for global validation metrics of the entry are shown in the following graphic. The table shows the number of entries on which the scores are based.

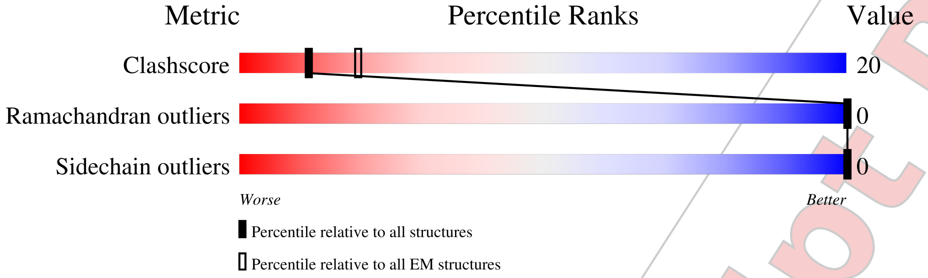


| Metric | Whole archive (#Entries) | EM structures (#Entries) |
|---|---|---|
| Clashscore | 158937 | 4297 |
| Ramachandran outliers | 154571 | 4023 |
| Sidechain outliers | 154315 | 3826 |

The table below summarises the geometric issues observed across the polymeric chains and their fit to the map. The red, orange, yellow and green segments of the bar indicate the fraction of residues that contain outliers for >=3, 2, 1 and 0 types of geometric quality criteria respectively. A grey segment represents the fraction of residues that are not modelled. The numeric value for each fraction is indicated below the corresponding segment, with a dot representing fractions <=5%

| Mol | Chain | Length | Quality of chain |
|---|---|---|---|
| 1 | A | 566 | 64% 36% |
| 1 | B | 566 | 64% 36% |
| 1 | C | 566 | 64% 36% |
| 1 | D | 566 | 65% 35% |
| 1 | E | 566 | 65% 35% |
| 1 | F | 566 | 65% 35% |
| 2 | H | 678 | 68% 32% |
| 2 | I | 678 | 69% 31% |
| 2 | J | 678 | 69% 31% |

*Continued on next page...*

Not For Manuscript Review

*Continued from previous page...*

| Mol | Chain | Length | Quality of chain |
|---|---|---|---|
| 2 | K | 678 | 68% 32% |
| 2 | L | 678 | 68% 32% |
| 2 | M | 678 | 68% 32% |

Not For Manuscript Review

#### 2 Entry composition (i)

There are 3 unique types of molecules in this entry. The entry contains 57846 atoms, of which 0 are hydrogens and 0 are deuteriums.

In the tables below, the AltConf column contains the number of residues with at least one atom in alternate conformation and the Trace column contains the number of residues modelled with at most 2 atoms.

- Molecule 1 is a protein.

| Mol | Chain | Residues | Atoms | | | | | AltConf | Trace |
|---|---|---|---|---|---|---|---|---|---|
| 1 | A | 566 | Total 4350 | C 2740 | N 776 | O 811 | S 23 | 0 | 0 |
| 1 | B | 566 | Total 4350 | C 2740 | N 776 | O 811 | S 23 | 0 | 0 |
| 1 | C | 566 | Total 4350 | C 2740 | N 776 | O 811 | S 23 | 0 | 0 |
| 1 | D | 566 | Total 4350 | C 2740 | N 776 | O 811 | S 23 | 0 | 0 |
| 1 | E | 566 | Total 4350 | C 2740 | N 776 | O 811 | S 23 | 0 | 0 |
| 1 | F | 566 | Total 4350 | C 2740 | N 776 | O 811 | S 23 | 0 | 0 |

- Molecule 2 is a protein.

| Mol | Chain | Residues | Atoms | | | | | AltConf | Trace |
|---|---|---|---|---|---|---|---|---|---|
| 2 | H | 678 | Total 5276 | C 3334 | N 931 | O 984 | S 27 | 0 | 0 |
| 2 | I | 678 | Total 5276 | C 3334 | N 931 | O 984 | S 27 | 0 | 0 |
| 2 | J | 678 | Total 5276 | C 3334 | N 931 | O 984 | S 27 | 0 | 0 |
| 2 | K | 678 | Total 5276 | C 3334 | N 931 | O 984 | S 27 | 0 | 0 |
| 2 | L | 678 | Total 5276 | C 3334 | N 931 | O 984 | S 27 | 0 | 0 |
| 2 | M | 678 | Total 5276 | C 3334 | N 931 | O 984 | S 27 | 0 | 0 |

- Molecule 3 is 5-(HEXAHYDRO-2-OXO-1H-THIENO[3,4-D]IMIDAZOL-6-YL)PENTANAL (three-letter code: BTI) (formula: $C_{10}H_{16}N_2O_2S$).

Not For Manuscript Review

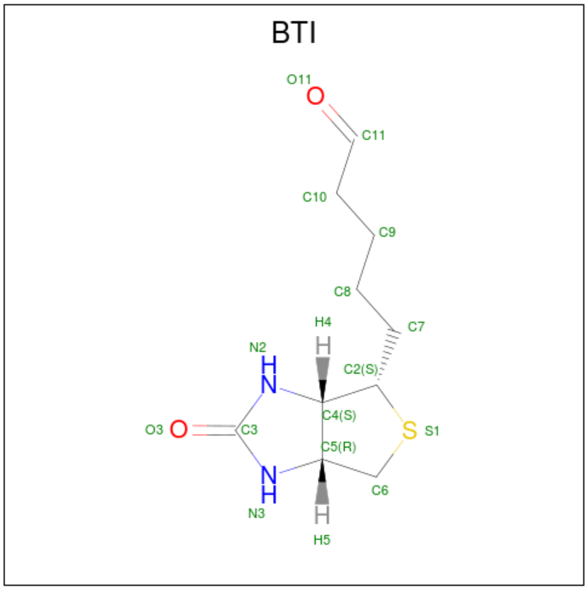


| Mol | Chain | Residues | Atoms | AltConf |
|---|---|---|---|---|
| 3 | G | 1 | Total C N O S<br>90 60 12 12 6 | 0 |
| 3 | G | 1 | Total C N O S<br>90 60 12 12 6 | 0 |
| 3 | G | 1 | Total C N O S<br>90 60 12 12 6 | 0 |
| 3 | G | 1 | Total C N O S<br>90 60 12 12 6 | 0 |
| 3 | G | 1 | Total C N O S<br>90 60 12 12 6 | 0 |
| 3 | G | 1 | Total C N O S<br>90 60 12 12 6 | 0 |

Not For Manuscript Review

### 3 Residue-property plots ⓘ

These plots are drawn for all protein, RNA, DNA and oligosaccharide chains in the entry. The first graphic for a chain summarises the proportions of the various outlier classes displayed in the second graphic. The second graphic shows the sequence view annotated by issues in geometry. Residues are color-coded according to the number of geometric quality criteria for which they contain at least one outlier: green = 0, yellow = 1, orange = 2 and red = 3 or more. Stretches of 2 or more consecutive residues without any outlier are shown as a green connector. Residues present in the sample, but not in the model, are shown in grey.

- Molecule 1:

Chain A:

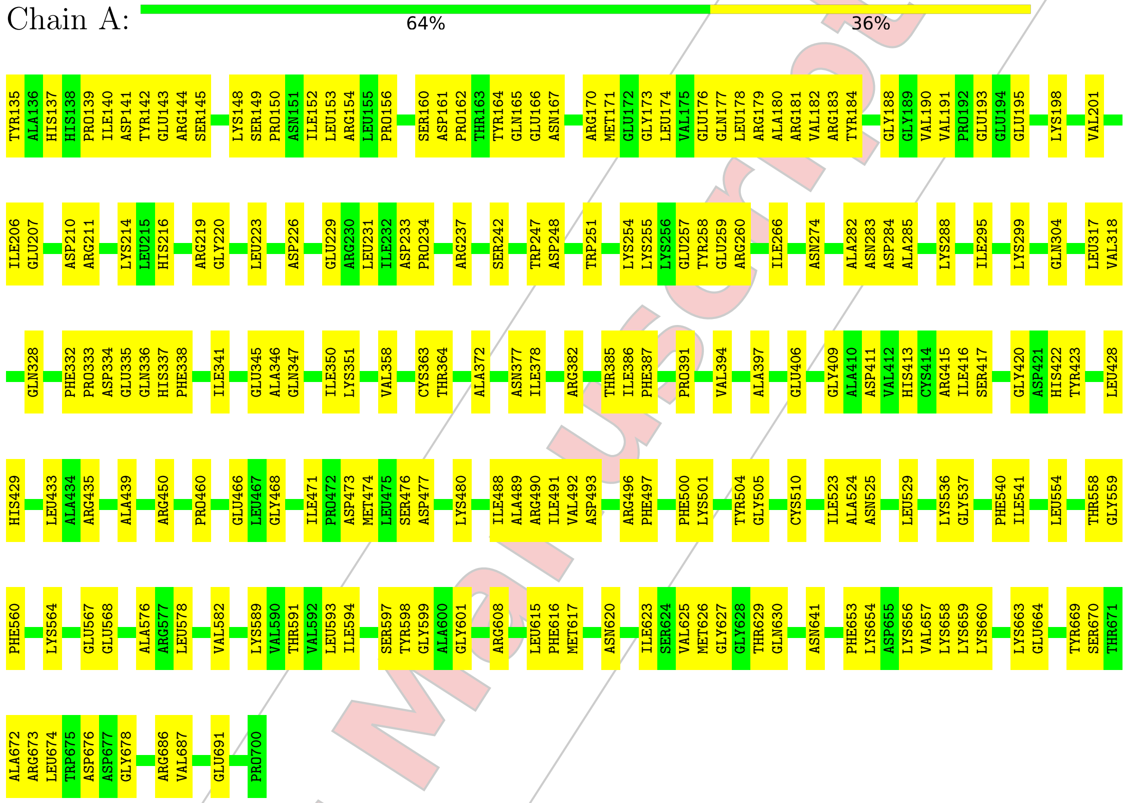


- Molecule 1:

Chain B:

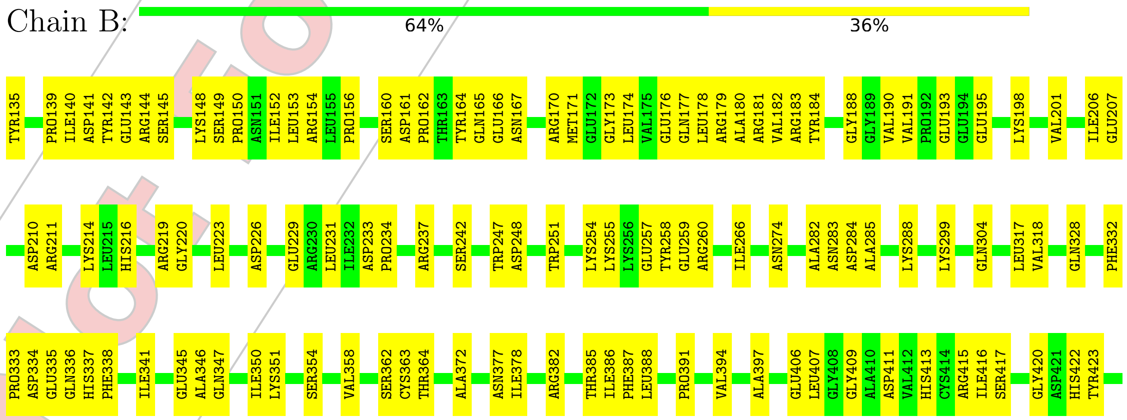


Not For Manuscript Review

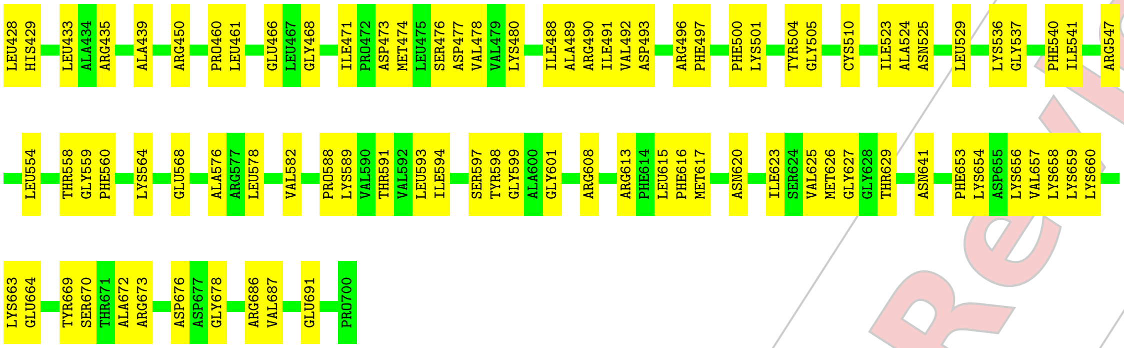


- Molecule 1:

Chain C: 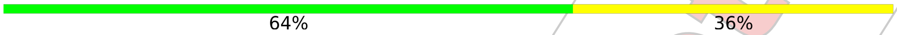


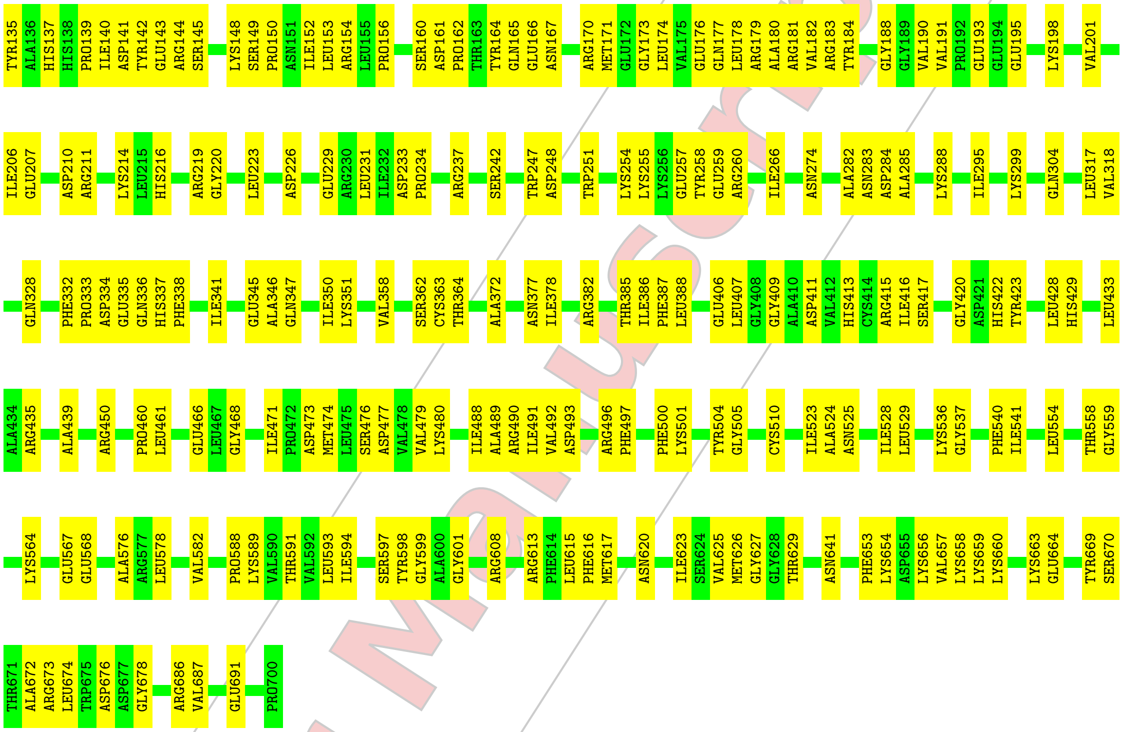


- Molecule 1:

Chain D: 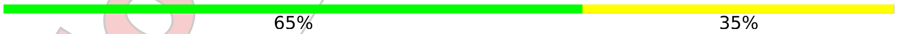


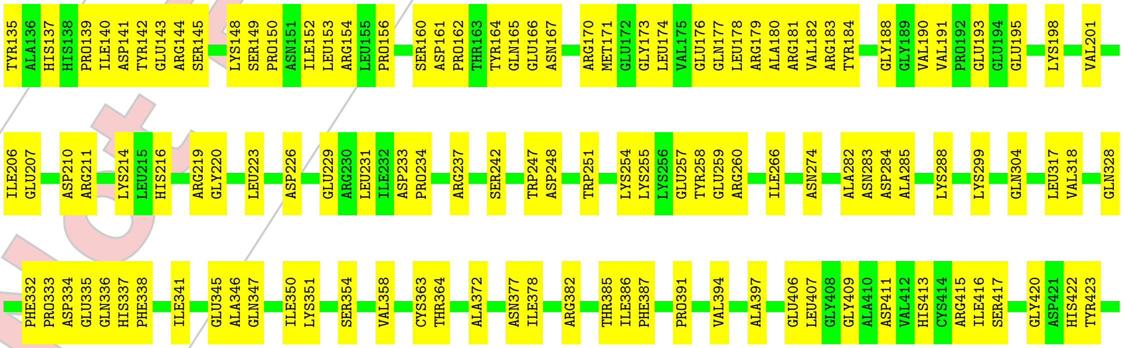




Not For Manuscript Review

LEU428 HIS429 LEU433 ALA434 ARG435 ALA439 ARG450 PRO460 GLU466 LEU467 GLY468 ILE471 PRO472 ASP473 SER476 ASP477 VAL478 VAL479 LYS480 ILE488 ALA489 ARG490 ILE491 VAL492 ASP493 ARG496 PHE497 PHE500 LYS501 TYR504 GLY505 CYS510 ILE523 ALA524 ASN525 LEU529 LYS536 GLY537 PHE540 ILE541 LEU554 THR558

GLY559 PHE560 LYS564 GLU567 GLU568 ALA576 ARG577 LEU578 VAL582 LYS589 VAL590 THR591 VAL592 LEU593 ILE594 SER597 TYR598 GLY599 ALA600 GLY601 ARG608 LEU615 PHE616 MET617 ILE623 SER624 VAL625 MET626 GLY627 GLY628 THR629 ASN641 PHE653 LYS654 ASP655 LYS656 VAL657 LYS658 LYS659 LYS660 LYS663 GLU664 TYR669 SER670 THR671 ALA672 ARG673

ASP676 ASP677 GLY678 ARG686 VAL687 GLU691 PRO700

• Molecule 1:

Chain E: 65% 35%

TYR135 ALA136 HIS137 HIS138 PRO139 ILE140 ASP141 TYR142 GLU143 ARG144 SER145 LYS148 SER149 PRO150 ASN151 ILE152 LEU153 ARG154 LEU155 PRO156 SER160 ASP161 PRO162 THR163 TYR164 GLN165 GLU166 ASN167 ARG170 MET171 GLU172 GLY173 LEU174 VAL175 GLU176 GLN177 LEU178 ARG179 ALA180 ARG181 VAL182 ARG183 TYR184 GLY188 GLY189 VAL190 VAL191 PRO192 GLU193 GLU194 GLU195 LYS198 VAL201

ILE206 GLU207 ASP210 ARG211 LYS214 LEU215 HIS216 ARG219 GLY220 LEU223 ASP226 GLU229 ARG230 LEU231 ILE232 ASP233 PRO234 GLY235 THR236 ARG237 SER242 TRP247 ASP248 TRP251 LYS254 LYS255 LYS256 GLU257 TYR258 GLU259 ARG260 ILE266 ASN274 ALA282 ASN283 ASP284 ALA285 LYS288 LYS299 GLN304 LEU317 VAL318

GLN328 PHE332 PRO333 ASP334 GLU335 GLN336 HIS337 ILE341 GLU345 ALA346 GLN347 ILE350 LYS351 VAL358 CYS363 THR364 ALA372 ASN377 ILE378 ARG382 THR385 ILE386 PHE387 PRO391 VAL394 ALA397 GLU406 LEU407 GLY408 GLY409 ALA410 ASP411 VAL412 HIS413 CYS414 ARG415 ILE416 SER417 GLY420 TYR423 LEU428 HIS429

LEU433 ALA434 ARG435 ALA439 ARG450 PRO460 GLU466 LEU467 GLY468 ILE471 PRO472 ASP473 SER476 ASP477 LYS480 ILE488 ALA489 ARG490 ILE491 VAL492 ASP493 ARG496 PHE497 PHE500 LYS501 TYR504 GLY505 CYS510 ILE523 ALA524 ASN525 LEU529 LYS536 GLY537 PHE540 ILE541 LEU554 THR558 GLY559 PHE560 LYS564

GLU567 GLU568 ALA576 ARG577 LEU578 VAL582 LYS589 VAL590 THR591 VAL592 LEU593 ILE594 SER597 TYR598 GLY599 ALA600 GLY601 ARG608 LEU615 PHE616 MET617 ILE623 SER624 VAL625 MET626 GLY627 GLY628 THR629 GLN630 ASN641 PHE653 LYS654 ASP655 LYS656 VAL657 LYS658 LYS659 LYS660 LYS663 GLU664 TYR669 ARG673 ASP676 ASP677 GLY678

ARG686 VAL687 GLU691 PRO700

• Molecule 1:

Chain F: 65% 35%

TYR135 ALA136 HIS137 HIS138 PRO139 ILE140 ASP141 TYR142 GLU143 ARG144 SER145 LYS148 SER149 PRO150 ASN151 ILE152 LEU153 ARG154 LEU155 PRO156 SER160 ASP161 PRO162 THR163 TYR164 GLN165 GLU166 ASN167 ARG170 MET171 GLU172 GLY173 LEU174 VAL175 GLU176 GLN177 LEU178 ARG179 ALA180 ARG181 VAL182 ARG183 TYR184 GLY188 GLY189 VAL190 VAL191 PRO192 GLU193 GLU194 GLU195 LYS198 VAL201

ILE206 GLU207 ASP210 ARG211 LYS214 LEU215 HIS216 ARG219 GLY220 LEU223 ASP226 GLU229 ARG230 LEU231 ILE232 ASP233 PRO234 GLY235 THR236 ARG237 SER242 GLN243 TRP247 ASP248 TRP251 LYS254 LYS255 LYS256 GLU257 TYR258 GLU259 ARG260 ILE266 ASN274 ALA282 ASN283 ASP284 ALA285 LYS288 LYS299 GLN304 LEU317 VAL318

GLN328 PHE332 PRO333 ASP334 GLU335 GLN336 HIS337 ILE341 GLU345 ALA346 GLN347 ILE350 LYS351 VAL358 CYS363 THR364 ALA372 ASN377 ILE378 ARG382 THR385 ILE386 PHE387 PRO391 VAL394 ALA397 GLU406 LEU407 GLY408 GLY409 ALA410 ASP411 VAL412 HIS413 CYS414 ARG415 ILE416 SER417 GLY420 TYR423 LEU428 HIS429

Not For Manuscript Review

LEU433 ALA434 ARG435 ALA439 ARG450 PRO460 LEU461 GLU466 LEU467 GLY468 ILE471 PRO472 ASP473 SER476 ASP477 VAL478 VAL479 LYS480 ILE488 ALA489 ARG490 ILE491 VAL492 ASP493 ARG496 PHE497 PHE500 LYS501 TYR504 GLY505 CYS510 ILE523 ALA524 ASN525 LEU529 LYS536 GLY537 PHE540 ILE541 ARG547 LEU554

THR558 GLY559 PHE560 LYS564 GLU567 GLU568 ALA576 ARG577 LEU578 VAL582 LYS589 VAL590 THR591 VAL592 LEU593 ILE594 SER597 TYR598 GLY599 ALA600 GLY601 ARG608 LEU615 PHE616 MET617 ILE623 SER624 VAL625 MET626 GLY627 GLY628 THR629 ASN641 PHE653 LYS654 ASP655 LYS656 VAL657 LYS658 LYS659 LYS660 LYS663 GLU664 TYR669 ARG673

ASP676 ASP677 GLY678 ARG686 VAL687 GLU691 PRO700

- Molecule 2:

Chain H: 68% 32%

GLU10 ARG11 LEU17 ASN20 ARG30 THR31 CYS32 ARG33 GLU34 ILE37 CYS44 GLU45 ARG48 GLU58 ILE62 GLY63 PRO64 PRO65 ASN69 SER70 TYR71 GLU75 VAL86 ILE89 TYR93 LEU96 SER97 GLU98 ASN99 ALA100 SER101 PHE102 ILE106 PRO116 PRO117 ALA118 LEU123 MET124 GLY125 SER126 LYS127 SER128 GLU129

SER130 LYS131 ARG132 ILE133 VAL139 PRO140 VAL141 GLY144 LEU155 GLU158 ALA159 LYS160 LYS161 PHE164 PRO165 LYS169 GLY173 VAL182 PHE188 THR189 PHE190 MET191 LEU192 PHE202 ILE209 ARG212 SER217 ARG218 HIS219 ILE220 GLU221 CYS222 GLN223 ILE224 PHE225 PHE226 ASP227 LYS228 VAL233 PHE234 PHE235 PHE236 GLU237 ARG238 ASP239 CYS240

ARG244 ARG245 GLU251 GLU252 ALA253 PRO254 HIS257 LEU258 SER259 MET260 GLU261 THR262 ARG263 GLN264 ARG265 ILE266 TYR279 VAL280 GLY281 ALA282 GLU286 PHE298 THR303 ARG304 HIS309 PRO310 VAL311 THR312 GLU313 GLU314 VAL315 CYS316 ARG317 ILE318 LYS327 LEU328 LYS331 THR332 LYS336 PRO337 LEU338 GLN342 VAL345 THR346 LEU347 SER350

VAL356 LEU372 ILE375 ARG376 GLU377 PRO378 ARG383 GLY384 PRO385 LEU408 SER413 TRP414 ARG418 GLU419 GLU420 LEU425 ARG426 LEU429 TYR432 LYS433 ILE441 LYS445 CYS448 PHE453 ALA454 ARG455 THR460 THR477 PRO478 GLU479 VAL480 MET483 ALA484 ALA485 THR486 ALA487 TRP488 LEU489 LEU490 ASN491 ASP494 ASN495

TRP496 ARG497 ASN503 THR506 PHE512 TYR513 VAL519 HIS524 ASN529 LYS532 ILE533 PHE534 VAL537 PHE544 THR552 SER553 ARG556 ILE561 VAL562 ASN563 ASP564 PHE565 THR566 PHE567 LEU568 PHE569 GLU570 ASN571 GLY572 MET573 HIS574 HIS575 THR576 VAL577 VAL580 ALA581 THR582 ASP585 VAL586 LEU599 LEU600 PRO601 LEU602 THR603 ASP604

GLY605 PHE606 GLY607 ASP608 SER609 ILE619 VAL620 SER621 VAL627 LEU630 LYS633 SER634 GLY635 ASP636 LEU637 VAL638 GLY641 LEU644 VAL645 ILE646 VAL647 MET650 LYS651 MET652 GLU653 HIS654 PRO655 VAL656 ARG657 ARG663 ASP677 LEU680 ALA681 GLU687

- Molecule 2:

Chain I: 69% 31%

GLU10 LEU17 ASN20 ARG30 THR31 CYS32 ARG33 GLU34 ILE37 CYS44 GLU45 ARG48 GLU58 ILE62 GLY63 PRO64 PRO65 ASN69 SER70 TYR71 VAL86 ILE89 TYR93 LEU96 SER97 GLU98 ASN99 ALA100 SER101 PHE102 ILE106 PRO116 PRO117 ALA118 LEU123 SER126 LYS127 SER128 GLU129 SER130 LYS131 ARG132 ILE133

VAL139 PRO140 VAL141 GLY144 LEU155 GLU158 ALA159 LYS160 LYS161 PHE164 PRO165 LYS169 ALA170 GLY173 LYS180 ILE181 VAL182 PHE188 THR189 PHE190 MET191 LEU192 LYS196 PHE202 ILE209 ARG212 SER217 ARG218 HIS219 ILE220 GLU221 CYS222 GLN223 ILE224 PHE225 PHE226 ASP227 LYS228 VAL233 PHE234 PHE235 PHE236 GLU237 ARG238 ASP239

CYS240 ARG244 ARG245 GLU251 GLU252 ALA253 PRO254 HIS257 LEU258 SER259 MET260 GLU261 THR262 ARG263 GLN264 ARG265 ILE266 TYR279 VAL280 GLU286 PHE298 THR303 ARG304 HIS309 PRO310 VAL311 THR312 GLU313 GLU314 VAL315 CYS316 ARG317 ILE318 LYS327 LEU328 LYS331 THR332 LYS336 PRO337 LEU338 GLN342 VAL345 THR346 LEU347 SER350

Not For Manuscript Review

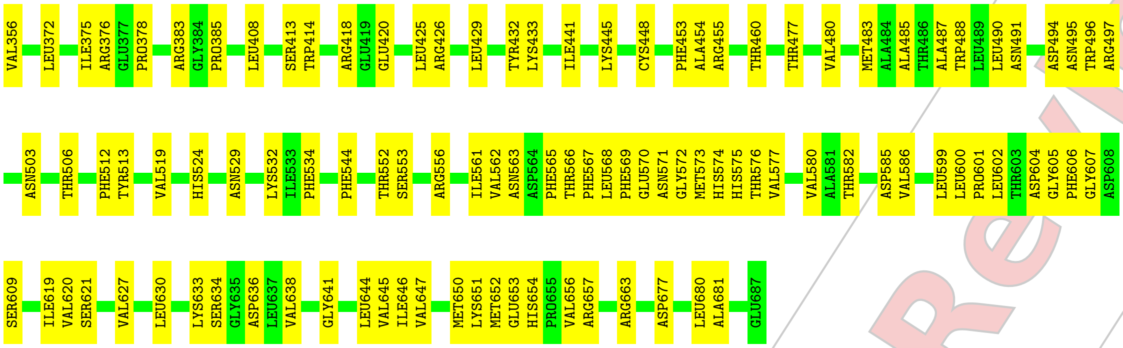


- Molecule 2:

Chain J:

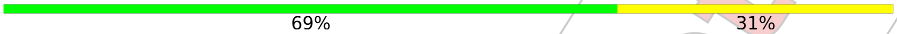


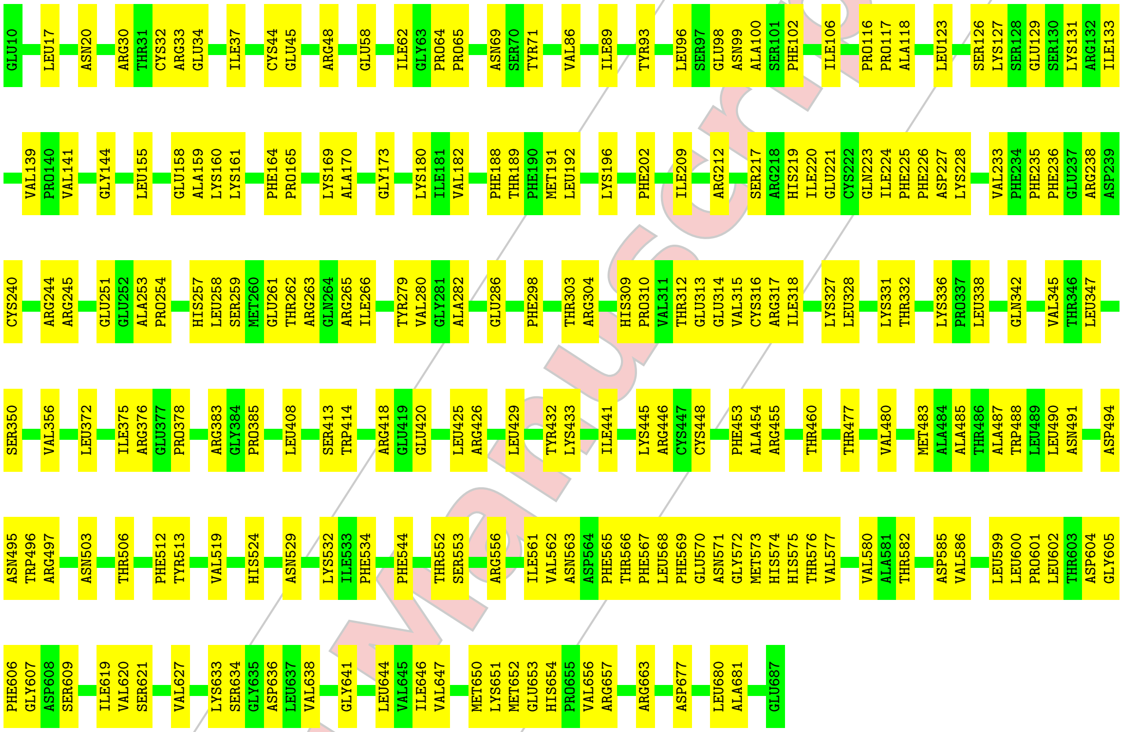


- Molecule 2:

Chain K:

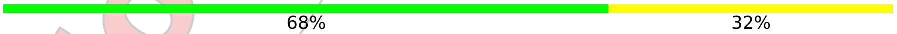


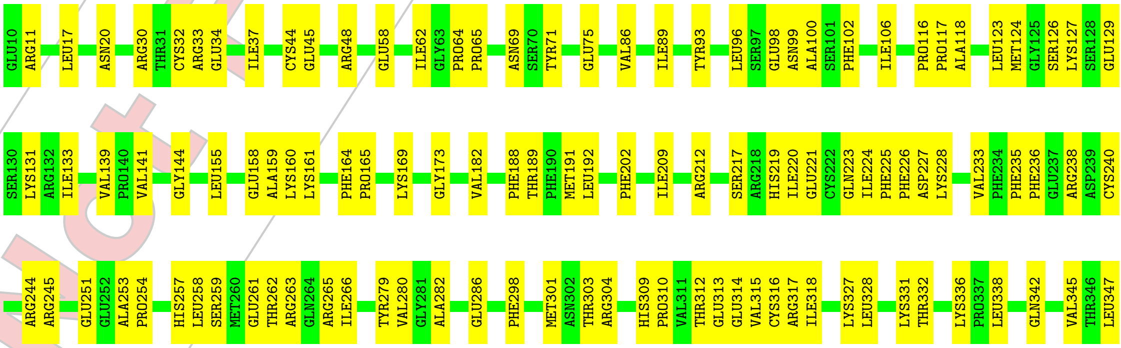


Not For Manuscript Review

SER350 VAL356 LEU372 THR373 PHE374 ILE375 ARG376 GLU377 PRO378 ARG383 GLY384 PRO385 LEU408 SER413 TRP414 ARG418 GLU419 GLU420 LEU425 ARG426 LEU429 TYR432 LYS433 VAL434 ALA435 ILE441 LYS445 ARG446 CYS447 CYS448 PHE453 ALA454 ARG455 THR460 THR477 VAL480 MET483 ALA484 ALA485 THR486 ALA487 TRP488 LEU489

LEU490 ASN491 ASP494 ASN495 TRP496 ARG497 ASN503 THR506 PHE512 TYR513 VAL519 HIS524 ASN529 LYS532 ILE533 PHE534 PHE544 THR552 SER553 ARG556 ILE561 VAL562 ASN563 ASP564 PHE565 THR566 PHE567 LEU568 PHE569 GLU570 ASN571 GLY572 MET573 HIS574 HIS575 THR576 VAL577 VAL580 ALA581 THR582 ASP585 VAL586 LEU599 LEU600 PRO601

ASP604 GLY605 PHE606 GLY607 ASP608 SER609 ILE619 VAL620 SER621 VAL627 LEU630 LYS633 SER634 GLY635 ASP636 LEU637 VAL638 GLY641 LEU644 VAL645 ILE646 VAL647 MET650 LYS651 MET652 GLU653 HIS654 PRO655 VAL656 ARG657 ARG663 ASP677 LEU680 ALA681 GLU687

- Molecule 2:

Chain L: 68% 32%

GLU10 LEU17 ASN20 ARG30 THR31 CYS32 ARG33 GLU34 ILE37 CYS44 GLU45 ARG48 GLU58 ILE62 GLY63 PRO64 PRO65 ASN69 SER70 TYR71 GLU75 VAL86 ILE89 TYR93 LEU96 SER97 GLU98 ASN99 ALA100 SER101 PHE102 ILE106 PRO116 PRO117 ALA118 LEU123 MET124 GLY125 SER126 LYS127 SER128 GLU129 SER130

LYS131 ARG132 ILE133 VAL139 PRO140 VAL141 GLY144 LEU155 GLU158 ALA159 LYS160 LYS161 PHE164 PRO165 LYS169 ALA170 GLY173 VAL182 PHE188 THR189 PHE190 MET191 LEU192 LYS196 PHE202 ILE209 ARG212 SER217 ARG218 HIS219 ILE220 GLU221 CYS222 GLN223 ILE224 PHE225 PHE226 ASP227 LYS228 VAL233 PHE234 PHE235 PHE236 GLU237 ARG238

ASP239 CYS240 ARG244 ARG245 GLU251 GLU252 ALA253 PRO254 HIS257 LEU258 SER259 MET260 GLU261 THR262 ARG263 GLN264 ARG265 ILE266 TYR279 VAL280 GLY281 ALA282 GLU286 MET301 ASN302 THR303 ARG304 HIS309 PRO310 VAL311 THR312 GLU313 GLU314 VAL315 CYS316 ARG317 ILE318 LYS327 LEU328 LYS331 THR332 LYS336 PRO337 LEU338 GLN342 VAL345 THR346 LEU347

SER350 VAL356 LEU372 ILE375 ARG376 GLU377 PRO378 ARG383 GLY384 PRO385 LEU408 SER413 TRP414 ARG418 GLU419 GLU420 LEU425 ARG426 LEU429 TYR432 LYS433 ILE441 LYS445 ARG446 CYS447 CYS448 PHE453 ALA454 ARG455 THR460 THR477 VAL480 MET483 ALA484 ALA485 THR486 ALA487 TRP488 LEU489 LEU490 ASN491

ASP494 ASN495 TRP496 ARG497 ASN503 THR506 PHE512 TYR513 VAL519 HIS524 ASN529 LYS532 ILE533 PHE534 PHE544 THR552 SER553 ARG556 ILE561 VAL562 ASN563 ASP564 PHE565 THR566 PHE567 LEU568 PHE569 GLU570 ASN571 GLY572 MET573 HIS574 HIS575 THR576 VAL577 VAL580 ALA581 THR582 ASP585 VAL586 LEU599 LEU600 PRO601 LEU602 THR603 ASP604

GLY605 PHE606 GLY607 ASP608 SER609 ILE619 VAL620 SER621 LYS626 VAL627 LEU630 LYS633 SER634 GLY635 ASP636 LEU637 VAL638 GLY641 LEU644 VAL645 ILE646 VAL647 MET650 LYS651 MET652 GLU653 HIS654 PRO655 VAL656 ARG657 ARG663 ASP677 LEU680 ALA681 GLU687

- Molecule 2:

Chain M: 68% 32%

GLU10 LEU17 ASN20 ARG30 THR31 CYS32 ARG33 GLU34 ILE37 CYS44 GLU45 ARG48 GLU58 ILE62 GLY63 PRO64 PRO65 ASN69 SER70 TYR71 GLU75 VAL86 ILE89 TYR93 LEU96 SER97 GLU98 ASN99 ALA100 SER101 PHE102 ILE106 PRO116 PRO117 ALA118 LEU123 MET124 GLY125 SER126 LYS127 SER128 GLU129 SER130

LYS131 ARG132 ILE133 VAL139 PRO140 VAL141 GLY144 LEU155 GLU158 ALA159 LYS160 LYS161 PHE164 PRO165 LYS169 ALA170 GLY173 LYS180 ILE181 VAL182 PHE188 THR189 PHE190 MET191 LEU192 LYS196 PHE202 ILE209 ARG212 SER217 ARG218 HIS219 ILE220 GLU221 CYS222 GLN223 ILE224 PHE225 PHE226 ASP227 LYS228 VAL233 PHE234 PHE235 PHE236

GLU237 ARG238 ASP239 CYS240 ARG244 ARG245 GLU251 GLU252 ALA253 PRO254 HIS257 LEU258 SER259 MET260 GLU261 THR262 ARG263 GLN264 ARG265 ILE266 TYR279 VAL280 GLY281 ALA282 GLU286 MET301 ASN302 THR303 ARG304 HIS309 PRO310 VAL311 THR312 GLU313 GLU314 VAL315 CYS316 ARG317 ILE318 LYS327 LEU328 LYS331 THR332 LYS336 PRO337 LEU338 GLN342 VAL345

Not For Manuscript Review

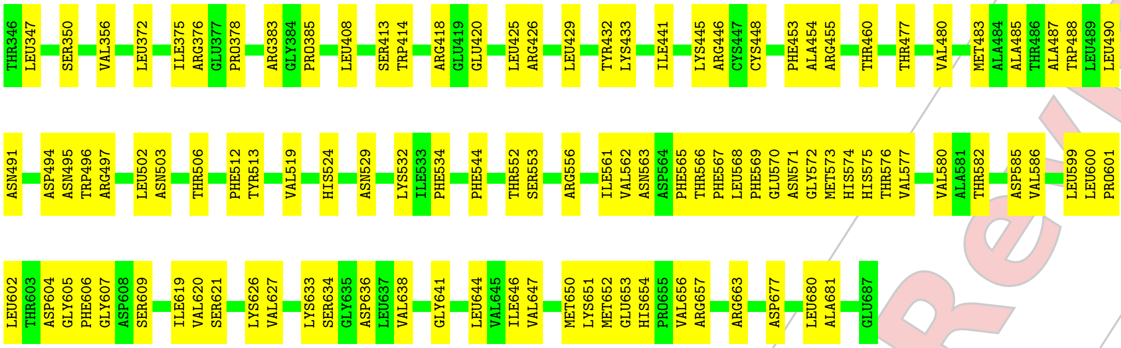


GLOBAL-STATISTICS INFOmissingINFO

Not For Manuscript Review

Not For Manuscript Review

### 4 Model quality (i)

#### 4.1 Standard geometry (i)

Bond lengths and bond angles in the following residue types are not validated in this section: BTI

The Z score for a bond length (or angle) is the number of standard deviations the observed value is removed from the expected value. A bond length (or angle) with $|Z| > 5$ is considered an outlier worth inspection. RMSZ is the root-mean-square of all Z scores of the bond lengths (or angles).

| Mol | Chain | Bond lengths | | Bond angles | |
|---|---|---|---|---|---|
| | | RMSZ | #$\|Z\|$ >5 | RMSZ | #$\|Z\|$ >5 |
| 1 | A | 0.31 | 0/4433 | 0.55 | 0/5998 |
| 1 | B | 0.31 | 0/4433 | 0.55 | 0/5998 |
| 1 | C | 0.31 | 0/4433 | 0.55 | 0/5998 |
| 1 | D | 0.31 | 0/4433 | 0.55 | 0/5998 |
| 1 | E | 0.31 | 0/4433 | 0.55 | 0/5998 |
| 1 | F | 0.31 | 0/4433 | 0.55 | 0/5998 |
| 2 | H | 0.31 | 0/5385 | 0.52 | 0/7274 |
| 2 | I | 0.31 | 0/5385 | 0.52 | 0/7274 |
| 2 | J | 0.31 | 0/5385 | 0.52 | 0/7274 |
| 2 | K | 0.31 | 0/5385 | 0.52 | 0/7274 |
| 2 | L | 0.31 | 0/5385 | 0.52 | 0/7274 |
| 2 | M | 0.31 | 0/5385 | 0.52 | 0/7274 |
| All | All | 0.31 | 0/58908 | 0.54 | 0/79632 |

There are no bond length outliers.

There are no bond angle outliers.

There are no chirality outliers.

There are no planarity outliers.

#### 4.2 Too-close contacts (i)

In the following table, the Non-H and H(model) columns list the number of non-hydrogen atoms and hydrogen atoms in the chain respectively. The H(added) column lists the number of hydrogen atoms added and optimized by MolProbity. The Clashes column lists the number of clashes within the asymmetric unit, whereas Symm-Clashes lists symmetry-related clashes.

| Mol | Chain | Non-H | H(model) | H(added) | Clashes | Symm-Clashes |
|---|---|---|---|---|---|---|
| 1 | A | 4350 | 0 | 4358 | 205 | 0 |

*Continued on next page...*

Continued from previous page...

| Mol | Chain | Non-H | H(model) | H(added) | Clashes | Symm-Clashes |
|---|---|---|---|---|---|---|
| 1 | B | 4350 | 0 | 4358 | 213 | 0 |
| 1 | C | 4350 | 0 | 4358 | 206 | 0 |
| 1 | D | 4350 | 0 | 4358 | 206 | 0 |
| 1 | E | 4350 | 0 | 4358 | 198 | 0 |
| 1 | F | 4350 | 0 | 4358 | 202 | 0 |
| 2 | H | 5276 | 0 | 5237 | 221 | 0 |
| 2 | I | 5276 | 0 | 5237 | 220 | 0 |
| 2 | J | 5276 | 0 | 5237 | 227 | 0 |
| 2 | K | 5276 | 0 | 5237 | 220 | 0 |
| 2 | L | 5276 | 0 | 5237 | 224 | 0 |
| 2 | M | 5276 | 0 | 5237 | 227 | 0 |
| 3 | G | 90 | 0 | 96 | 13 | 0 |
| All | All | 57846 | 0 | 57666 | 2307 | 0 |

The all-atom clashscore is defined as the number of clashes found per 1000 atoms (including hydrogen atoms). The all-atom clashscore for this structure is 20.

All (2307) close contacts within the same asymmetric unit are listed below, sorted by their clash magnitude.

| Atom-1 | Atom-2 | Interatomic distance (Å) | Clash overlap (Å) |
|---|---|---|---|
| 2:I:650:MET:CE | 2:I:651:LYS:CD | 1.78 | 1.61 |
| 2:J:650:MET:CE | 2:J:651:LYS:CD | 1.78 | 1.61 |
| 2:M:650:MET:CE | 2:M:651:LYS:CD | 1.78 | 1.59 |
| 2:L:650:MET:CE | 2:L:651:LYS:CD | 1.78 | 1.59 |
| 2:K:650:MET:CE | 2:K:651:LYS:CD | 1.78 | 1.57 |
| 2:H:650:MET:CE | 2:H:651:LYS:CD | 1.78 | 1.57 |
| 2:J:650:MET:CE | 2:J:651:LYS:HD2 | 1.37 | 1.45 |
| 2:I:650:MET:CE | 2:I:651:LYS:HD2 | 1.37 | 1.45 |
| 2:H:650:MET:HE1 | 2:H:651:LYS:NZ | 1.30 | 1.43 |
| 2:K:650:MET:HE1 | 2:K:651:LYS:NZ | 1.30 | 1.43 |
| 2:L:650:MET:CE | 2:L:651:LYS:HD2 | 1.37 | 1.41 |
| 2:M:650:MET:CE | 2:M:651:LYS:HD2 | 1.37 | 1.41 |
| 2:H:650:MET:CE | 2:H:651:LYS:HD2 | 1.37 | 1.40 |
| 2:K:650:MET:CE | 2:K:651:LYS:HD2 | 1.37 | 1.40 |
| 2:M:650:MET:HE1 | 2:M:651:LYS:NZ | 1.51 | 1.25 |
| 2:L:650:MET:HE1 | 2:L:651:LYS:NZ | 1.51 | 1.25 |
| 2:K:650:MET:HE1 | 2:K:651:LYS:CE | 1.67 | 1.23 |
| 2:H:650:MET:HE1 | 2:H:651:LYS:CE | 1.67 | 1.22 |
| 2:I:650:MET:HE2 | 2:I:651:LYS:NZ | 1.54 | 1.22 |
| 2:J:650:MET:HE2 | 2:J:651:LYS:NZ | 1.56 | 1.20 |
| 2:M:650:MET:HE2 | 2:M:651:LYS:CD | 1.52 | 1.17 |

Continued on next page...

Continued from previous page...

| Atom-1 | Atom-2 | Interatomic distance (Å) | Clash overlap (Å) |
|---|---|---|---|
| 2:L:650:MET:HE1 | 2:L:651:LYS:CE | 1.75 | 1.17 |
| 2:M:650:MET:HE1 | 2:M:651:LYS:CE | 1.75 | 1.17 |
| 2:L:650:MET:HE2 | 2:L:651:LYS:CD | 1.52 | 1.16 |
| 2:I:650:MET:HE3 | 2:I:651:LYS:CD | 1.56 | 1.14 |
| 2:K:650:MET:HE2 | 2:K:651:LYS:CD | 1.49 | 1.14 |
| 2:J:650:MET:HE3 | 2:J:651:LYS:CD | 1.56 | 1.13 |
| 2:H:650:MET:HE2 | 2:H:651:LYS:CD | 1.49 | 1.13 |
| 2:K:225:PHE:HE2 | 2:K:338:LEU:HD11 | 1.16 | 1.10 |
| 2:H:225:PHE:HE2 | 2:H:338:LEU:HD11 | 1.16 | 1.09 |
| 2:I:650:MET:CE | 2:I:651:LYS:CE | 2.30 | 1.08 |
| 2:J:650:MET:CE | 2:J:651:LYS:CE | 2.30 | 1.08 |
| 2:M:650:MET:CE | 2:M:651:LYS:CE | 2.30 | 1.07 |
| 2:I:225:PHE:HE2 | 2:I:338:LEU:HD11 | 1.16 | 1.07 |
| 2:I:650:MET:CE | 2:I:651:LYS:HD3 | 1.78 | 1.07 |
| 2:J:650:MET:CE | 2:J:651:LYS:HD3 | 1.78 | 1.07 |
| 2:L:650:MET:CE | 2:L:651:LYS:CE | 2.30 | 1.07 |
| 2:M:225:PHE:HE2 | 2:M:338:LEU:HD11 | 1.16 | 1.07 |
| 2:J:225:PHE:HE2 | 2:J:338:LEU:HD11 | 1.16 | 1.06 |
| 2:L:225:PHE:HE2 | 2:L:338:LEU:HD11 | 1.16 | 1.06 |
| 2:I:650:MET:HE2 | 2:I:651:LYS:CE | 1.86 | 1.06 |
| 2:J:650:MET:HE2 | 2:J:651:LYS:CE | 1.87 | 1.05 |
| 2:K:650:MET:CE | 2:K:651:LYS:NZ | 2.19 | 1.05 |
| 2:H:650:MET:CE | 2:H:651:LYS:NZ | 2.19 | 1.04 |
| 1:D:560:PHE:O | 3:G:803:BTI:S1 | 2.15 | 1.04 |
| 2:M:650:MET:CE | 2:M:651:LYS:NZ | 2.19 | 1.03 |
| 2:I:650:MET:CE | 2:I:651:LYS:NZ | 2.19 | 1.03 |
| 2:L:650:MET:CE | 2:L:651:LYS:NZ | 2.19 | 1.03 |
| 2:J:650:MET:CE | 2:J:651:LYS:NZ | 2.19 | 1.02 |
| 2:K:650:MET:CE | 2:K:651:LYS:CE | 2.30 | 1.02 |
| 2:H:650:MET:CE | 2:H:651:LYS:CE | 2.30 | 1.02 |
| 2:K:573:MET:CE | 2:M:58:GLU:OE1 | 2.07 | 1.02 |
| 2:I:650:MET:HE1 | 2:I:651:LYS:HD2 | 1.39 | 1.02 |
| 2:J:650:MET:HE1 | 2:J:651:LYS:HD2 | 1.39 | 1.01 |
| 1:F:428:LEU:HB3 | 2:M:606:PHE:HZ | 1.23 | 1.01 |
| 2:H:573:MET:CE | 2:L:58:GLU:OE1 | 2.10 | 1.00 |
| 2:I:573:MET:CE | 2:K:58:GLU:OE1 | 2.10 | 1.00 |
| 2:M:650:MET:HE2 | 2:M:651:LYS:HD3 | 1.02 | 1.00 |
| 2:L:650:MET:HE2 | 2:L:651:LYS:HD3 | 1.02 | 1.00 |
| 2:M:650:MET:HE1 | 2:M:651:LYS:HZ2 | 1.05 | 0.99 |
| 2:J:58:GLU:OE1 | 2:L:573:MET:CE | 2.09 | 0.99 |
| 2:J:650:MET:HE2 | 2:J:651:LYS:CD | 1.67 | 0.98 |

Continued on next page...

Not For Manuscript Review

*Continued from previous page...*

| Atom-1 | Atom-2 | Interatomic distance (Å) | Clash overlap (Å) |
|---|---|---|---|
| 2:L:650:MET:HE1 | 2:L:651:LYS:HZ2 | 1.05 | 0.98 |
| 1:F:560:PHE:O | 3:G:801:BTI:S1 | 2.21 | 0.98 |
| 2:I:650:MET:HE3 | 2:I:651:LYS:HD2 | 1.00 | 0.98 |
| 2:H:58:GLU:OE1 | 2:J:573:MET:CE | 2.11 | 0.98 |
| 2:I:58:GLU:OE1 | 2:M:573:MET:CE | 2.11 | 0.97 |
| 2:I:650:MET:HE2 | 2:I:651:LYS:CD | 1.67 | 0.97 |
| 2:J:650:MET:HE3 | 2:J:651:LYS:HD2 | 0.99 | 0.97 |
| 1:F:428:LEU:HB3 | 2:M:606:PHE:CZ | 2.00 | 0.96 |
| 2:H:650:MET:HE2 | 2:H:651:LYS:HD3 | 1.00 | 0.96 |
| 2:K:650:MET:HE2 | 2:K:651:LYS:HD3 | 1.00 | 0.96 |
| 2:H:34:GLU:OE2 | 2:J:383:ARG:HD2 | 1.65 | 0.96 |
| 2:K:383:ARG:HD2 | 2:M:34:GLU:OE2 | 1.65 | 0.96 |
| 2:J:34:GLU:OE2 | 2:L:383:ARG:HD2 | 1.66 | 0.95 |
| 2:H:383:ARG:HD2 | 2:L:34:GLU:OE2 | 1.66 | 0.95 |
| 1:B:560:PHE:O | 3:G:805:BTI:S1 | 2.24 | 0.95 |
| 2:I:34:GLU:OE2 | 2:M:383:ARG:HD2 | 1.67 | 0.94 |
| 2:J:650:MET:HE2 | 2:J:651:LYS:HD3 | 1.40 | 0.94 |
| 2:J:650:MET:HE1 | 2:J:651:LYS:CD | 1.95 | 0.94 |
| 2:I:650:MET:HE1 | 2:I:651:LYS:CD | 1.95 | 0.94 |
| 2:J:650:MET:HE2 | 2:J:651:LYS:HZ3 | 1.05 | 0.93 |
| 2:I:383:ARG:HD2 | 2:K:34:GLU:OE2 | 1.68 | 0.93 |
| 2:H:225:PHE:CE2 | 2:H:338:LEU:HD11 | 2.04 | 0.93 |
| 2:I:650:MET:HE2 | 2:I:651:LYS:HD3 | 1.41 | 0.93 |
| 2:K:225:PHE:CE2 | 2:K:338:LEU:HD11 | 2.04 | 0.93 |
| 2:I:650:MET:HE2 | 2:I:651:LYS:HZ2 | 1.02 | 0.92 |
| 2:H:650:MET:HE3 | 2:H:651:LYS:HD2 | 0.93 | 0.92 |
| 2:K:650:MET:HE3 | 2:K:651:LYS:HD2 | 0.93 | 0.92 |
| 2:I:225:PHE:CE2 | 2:I:338:LEU:HD11 | 2.04 | 0.92 |
| 2:J:225:PHE:CE2 | 2:J:338:LEU:HD11 | 2.04 | 0.92 |
| 2:L:225:PHE:CE2 | 2:L:338:LEU:HD11 | 2.04 | 0.92 |
| 2:M:225:PHE:CE2 | 2:M:338:LEU:HD11 | 2.04 | 0.91 |
| 2:J:228:LYS:HG3 | 2:J:280:VAL:HG21 | 1.51 | 0.91 |
| 2:I:228:LYS:HG3 | 2:I:280:VAL:HG21 | 1.52 | 0.91 |
| 2:L:233:VAL:HG23 | 2:L:342:GLN:HG2 | 1.53 | 0.91 |
| 2:M:233:VAL:HG23 | 2:M:342:GLN:HG2 | 1.53 | 0.90 |
| 2:H:233:VAL:HG23 | 2:H:342:GLN:HG2 | 1.53 | 0.90 |
| 2:K:565:PHE:HB2 | 2:K:577:VAL:HG23 | 1.54 | 0.90 |
| 2:H:565:PHE:HB2 | 2:H:577:VAL:HG23 | 1.54 | 0.90 |
| 2:K:233:VAL:HG23 | 2:K:342:GLN:HG2 | 1.53 | 0.90 |
| 2:L:228:LYS:HG3 | 2:L:280:VAL:HG21 | 1.52 | 0.89 |
| 2:M:228:LYS:HG3 | 2:M:280:VAL:HG21 | 1.51 | 0.89 |

*Continued on next page...*

Not For Manuscript Review

Continued from previous page...

| Atom-1 | Atom-2 | Interatomic distance (Å) | Clash overlap (Å) |
|---|---|---|---|
| 2:H:650:MET:CE | 2:H:651:LYS:HZ2 | 1.79 | 0.89 |
| 2:K:650:MET:CE | 2:K:651:LYS:HZ2 | 1.79 | 0.89 |
| 2:M:565:PHE:HB2 | 2:M:577:VAL:HG23 | 1.54 | 0.89 |
| 2:K:228:LYS:HG3 | 2:K:280:VAL:HG21 | 1.51 | 0.89 |
| 2:L:565:PHE:HB2 | 2:L:577:VAL:HG23 | 1.54 | 0.89 |
| 2:I:233:VAL:HG23 | 2:I:342:GLN:HG2 | 1.53 | 0.89 |
| 2:H:228:LYS:HG3 | 2:H:280:VAL:HG21 | 1.52 | 0.89 |
| 2:J:233:VAL:HG23 | 2:J:342:GLN:HG2 | 1.53 | 0.88 |
| 2:I:650:MET:CE | 2:I:651:LYS:HZ2 | 1.84 | 0.88 |
| 1:A:669:TYR:HA | 1:A:673:ARG:HE | 1.40 | 0.87 |
| 1:D:669:TYR:HA | 1:D:673:ARG:HE | 1.40 | 0.87 |
| 1:E:669:TYR:HA | 1:E:673:ARG:HE | 1.40 | 0.87 |
| 1:F:669:TYR:HA | 1:F:673:ARG:HE | 1.40 | 0.87 |
| 1:B:669:TYR:HA | 1:B:673:ARG:HE | 1.40 | 0.87 |
| 1:C:669:TYR:HA | 1:C:673:ARG:HE | 1.40 | 0.87 |
| 2:J:565:PHE:HB2 | 2:J:577:VAL:HG23 | 1.55 | 0.86 |
| 2:I:565:PHE:HB2 | 2:I:577:VAL:HG23 | 1.54 | 0.86 |
| 1:C:428:LEU:HB3 | 2:J:606:PHE:HZ | 1.41 | 0.86 |
| 2:K:646:ILE:HD11 | 2:K:653:GLU:HB2 | 1.57 | 0.86 |
| 2:H:646:ILE:HD11 | 2:H:653:GLU:HB2 | 1.57 | 0.86 |
| 2:L:650:MET:HE3 | 2:L:651:LYS:HD2 | 0.87 | 0.86 |
| 2:M:650:MET:HE3 | 2:M:651:LYS:HD2 | 0.86 | 0.86 |
| 2:J:650:MET:CE | 2:J:651:LYS:HZ3 | 1.86 | 0.86 |
| 2:J:513:TYR:HB2 | 2:J:600:LEU:HA | 1.57 | 0.85 |
| 2:I:513:TYR:HB2 | 2:I:600:LEU:HA | 1.57 | 0.85 |
| 2:H:513:TYR:HB2 | 2:H:600:LEU:HA | 1.57 | 0.85 |
| 2:K:513:TYR:HB2 | 2:K:600:LEU:HA | 1.58 | 0.85 |
| 2:M:650:MET:CE | 2:M:651:LYS:HZ2 | 1.82 | 0.85 |
| 1:E:234:PRO:HB2 | 2:L:495:ASN:HD21 | 1.41 | 0.85 |
| 1:A:234:PRO:HB2 | 2:H:495:ASN:HD21 | 1.42 | 0.85 |
| 1:C:428:LEU:HB3 | 2:J:606:PHE:CZ | 2.12 | 0.85 |
| 2:L:318:ILE:HG22 | 2:L:345:VAL:HG22 | 1.58 | 0.85 |
| 2:L:650:MET:CE | 2:L:651:LYS:HZ2 | 1.82 | 0.85 |
| 2:M:318:ILE:HG22 | 2:M:345:VAL:HG22 | 1.58 | 0.85 |
| 1:E:428:LEU:HB3 | 2:L:606:PHE:CZ | 2.12 | 0.84 |
| 2:J:646:ILE:HD11 | 2:J:653:GLU:HB2 | 1.57 | 0.84 |
| 2:I:646:ILE:HD11 | 2:I:653:GLU:HB2 | 1.57 | 0.84 |
| 2:L:650:MET:HE3 | 2:L:651:LYS:CD | 1.72 | 0.84 |
| 2:M:650:MET:HE3 | 2:M:651:LYS:CD | 1.72 | 0.84 |
| 2:L:513:TYR:HB2 | 2:L:600:LEU:HA | 1.58 | 0.84 |
| 2:M:513:TYR:HB2 | 2:M:600:LEU:HA | 1.58 | 0.84 |

Continued on next page...

Not For Manuscript Review

Continued from previous page...

| Atom-1 | Atom-2 | Interatomic distance (Å) | Clash overlap (Å) |
|---|---|---|---|
| 2:J:318:ILE:HG22 | 2:J:345:VAL:HG22 | 1.58 | 0.83 |
| 2:I:318:ILE:HG22 | 2:I:345:VAL:HG22 | 1.59 | 0.83 |
| 2:L:646:ILE:HD11 | 2:L:653:GLU:HB2 | 1.57 | 0.83 |
| 2:M:646:ILE:HD11 | 2:M:653:GLU:HB2 | 1.57 | 0.83 |
| 1:C:234:PRO:HB2 | 2:J:495:ASN:HD21 | 1.42 | 0.83 |
| 1:D:428:LEU:HB3 | 2:K:606:PHE:HZ | 1.42 | 0.83 |
| 2:H:318:ILE:HG22 | 2:H:345:VAL:HG22 | 1.58 | 0.83 |
| 1:B:234:PRO:HB2 | 2:I:495:ASN:HD21 | 1.44 | 0.83 |
| 2:K:318:ILE:HG22 | 2:K:345:VAL:HG22 | 1.59 | 0.82 |
| 2:J:158:GLU:HA | 2:J:161:LYS:HB2 | 1.62 | 0.82 |
| 2:I:158:GLU:HA | 2:I:161:LYS:HB2 | 1.62 | 0.82 |
| 1:E:428:LEU:HB3 | 2:L:606:PHE:HZ | 1.41 | 0.81 |
| 2:H:158:GLU:HA | 2:H:161:LYS:HB2 | 1.62 | 0.81 |
| 2:K:158:GLU:HA | 2:K:161:LYS:HB2 | 1.62 | 0.81 |
| 2:L:223:GLN:HB3 | 2:L:235:PHE:HB2 | 1.63 | 0.81 |
| 1:A:428:LEU:HB3 | 2:H:606:PHE:HZ | 1.44 | 0.81 |
| 1:A:428:LEU:HB3 | 2:H:606:PHE:CZ | 2.14 | 0.80 |
| 2:M:223:GLN:HB3 | 2:M:235:PHE:HB2 | 1.63 | 0.80 |
| 2:J:650:MET:HE1 | 2:J:651:LYS:CE | 2.06 | 0.80 |
| 2:L:158:GLU:HA | 2:L:161:LYS:HB2 | 1.62 | 0.80 |
| 2:M:158:GLU:HA | 2:M:161:LYS:HB2 | 1.62 | 0.80 |
| 1:A:429:HIS:CD2 | 2:H:607:GLY:HA2 | 2.17 | 0.80 |
| 1:D:428:LEU:HB3 | 2:K:606:PHE:CZ | 2.17 | 0.80 |
| 2:I:650:MET:HE1 | 2:I:651:LYS:CE | 2.07 | 0.80 |
| 2:J:223:GLN:HB3 | 2:J:235:PHE:HB2 | 1.63 | 0.79 |
| 2:I:223:GLN:HB3 | 2:I:235:PHE:HB2 | 1.63 | 0.79 |
| 2:K:562:VAL:HG13 | 2:K:580:VAL:HG22 | 1.64 | 0.79 |
| 2:L:562:VAL:HG13 | 2:L:580:VAL:HG22 | 1.64 | 0.79 |
| 2:M:562:VAL:HG13 | 2:M:580:VAL:HG22 | 1.64 | 0.79 |
| 2:H:223:GLN:HB3 | 2:H:235:PHE:HB2 | 1.63 | 0.79 |
| 2:H:562:VAL:HG13 | 2:H:580:VAL:HG22 | 1.64 | 0.79 |
| 2:K:223:GLN:HB3 | 2:K:235:PHE:HB2 | 1.63 | 0.79 |
| 2:J:562:VAL:HG13 | 2:J:580:VAL:HG22 | 1.64 | 0.78 |
| 1:B:428:LEU:HB3 | 2:I:606:PHE:HZ | 1.48 | 0.78 |
| 2:I:562:VAL:HG13 | 2:I:580:VAL:HG22 | 1.64 | 0.78 |
| 1:D:142:TYR:OH | 1:F:234:PRO:HD2 | 1.85 | 0.77 |
| 1:E:429:HIS:CD2 | 2:L:607:GLY:HA2 | 2.20 | 0.77 |
| 1:B:234:PRO:HD2 | 1:F:142:TYR:OH | 1.84 | 0.77 |
| 1:F:179:ARG:HA | 2:M:503:ASN:HB3 | 1.67 | 0.76 |
| 2:K:627:VAL:HG12 | 2:K:647:VAL:HG22 | 1.68 | 0.76 |
| 1:B:142:TYR:OH | 1:D:234:PRO:HD2 | 1.86 | 0.76 |

Continued on next page...

Not For Manuscript Review

Continued from previous page...

| Atom-1 | Atom-2 | Interatomic distance (Å) | Clash overlap (Å) |
|---|---|---|---|
| 2:H:627:VAL:HG12 | 2:H:647:VAL:HG22 | 1.68 | 0.76 |
| 2:J:627:VAL:HG12 | 2:J:647:VAL:HG22 | 1.68 | 0.76 |
| 2:I:627:VAL:HG12 | 2:I:647:VAL:HG22 | 1.68 | 0.76 |
| 1:A:142:TYR:OH | 1:E:234:PRO:HD2 | 1.86 | 0.76 |
| 1:C:429:HIS:CD2 | 2:J:607:GLY:HA2 | 2.19 | 0.76 |
| 2:K:650:MET:HE1 | 2:K:651:LYS:HZ2 | 0.91 | 0.76 |
| 2:I:650:MET:HE3 | 2:I:651:LYS:HD3 | 1.51 | 0.75 |
| 2:H:650:MET:HE1 | 2:H:651:LYS:HZ2 | 0.91 | 0.75 |
| 2:L:627:VAL:HG12 | 2:L:647:VAL:HG22 | 1.67 | 0.75 |
| 2:M:627:VAL:HG12 | 2:M:647:VAL:HG22 | 1.68 | 0.75 |
| 1:C:234:PRO:HD2 | 1:E:142:TYR:OH | 1.86 | 0.75 |
| 1:A:234:PRO:HD2 | 1:C:142:TYR:OH | 1.87 | 0.74 |
| 2:H:573:MET:HE1 | 2:L:58:GLU:OE1 | 1.86 | 0.74 |
| 1:C:142:TYR:HA | 1:C:145:SER:HB3 | 1.70 | 0.74 |
| 1:B:142:TYR:HA | 1:B:145:SER:HB3 | 1.70 | 0.74 |
| 2:J:650:MET:HE3 | 2:J:651:LYS:HD3 | 1.53 | 0.74 |
| 1:D:142:TYR:HA | 1:D:145:SER:HB3 | 1.70 | 0.73 |
| 1:B:558:THR:OG1 | 1:B:598:TYR:O | 2.06 | 0.73 |
| 1:C:558:THR:OG1 | 1:C:598:TYR:O | 2.06 | 0.73 |
| 1:A:142:TYR:HA | 1:A:145:SER:HB3 | 1.70 | 0.73 |
| 1:F:142:TYR:HA | 1:F:145:SER:HB3 | 1.70 | 0.73 |
| 2:H:44:CYS:SG | 2:H:65:PRO:HA | 2.28 | 0.73 |
| 1:E:142:TYR:HA | 1:E:145:SER:HB3 | 1.70 | 0.73 |
| 1:A:558:THR:OG1 | 1:A:598:TYR:O | 2.06 | 0.73 |
| 1:D:429:HIS:CD2 | 2:K:607:GLY:HA2 | 2.24 | 0.73 |
| 1:D:558:THR:OG1 | 1:D:598:TYR:O | 2.06 | 0.73 |
| 2:L:44:CYS:SG | 2:L:65:PRO:HA | 2.28 | 0.73 |
| 2:M:44:CYS:SG | 2:M:65:PRO:HA | 2.28 | 0.73 |
| 2:I:44:CYS:SG | 2:I:65:PRO:HA | 2.28 | 0.73 |
| 2:J:44:CYS:SG | 2:J:65:PRO:HA | 2.28 | 0.73 |
| 2:K:44:CYS:SG | 2:K:65:PRO:HA | 2.28 | 0.73 |
| 2:L:123:LEU:HD21 | 2:L:133:ILE:HG13 | 1.71 | 0.73 |
| 2:M:123:LEU:HD21 | 2:M:133:ILE:HG13 | 1.71 | 0.73 |
| 2:K:573:MET:HE1 | 2:M:58:GLU:OE1 | 1.88 | 0.73 |
| 1:B:428:LEU:HB3 | 2:I:606:PHE:CZ | 2.24 | 0.72 |
| 2:H:123:LEU:HD21 | 2:H:133:ILE:HG13 | 1.71 | 0.72 |
| 2:K:123:LEU:HD21 | 2:K:133:ILE:HG13 | 1.71 | 0.72 |
| 2:L:64:PRO:HD2 | 2:L:69:ASN:HB3 | 1.72 | 0.72 |
| 2:M:64:PRO:HD2 | 2:M:69:ASN:HB3 | 1.72 | 0.72 |
| 1:F:558:THR:OG1 | 1:F:598:TYR:O | 2.06 | 0.72 |
| 1:E:558:THR:OG1 | 1:E:598:TYR:O | 2.06 | 0.72 |

Continued on next page...

Not For Manuscript Review

Continued from previous page...

| Atom-1 | Atom-2 | Interatomic distance (Å) | Clash overlap (Å) |
|---|---|---|---|
| 2:K:64:PRO:HD2 | 2:K:69:ASN:HB3 | 1.72 | 0.71 |
| 2:H:64:PRO:HD2 | 2:H:69:ASN:HB3 | 1.72 | 0.71 |
| 2:I:372:LEU:HB3 | 2:I:375:ILE:HD11 | 1.71 | 0.71 |
| 2:J:372:LEU:HB3 | 2:J:375:ILE:HD11 | 1.71 | 0.71 |
| 1:B:334:ASP:HB3 | 1:B:337:HIS:CE1 | 2.26 | 0.71 |
| 1:C:334:ASP:HB3 | 1:C:337:HIS:CE1 | 2.26 | 0.71 |
| 2:M:372:LEU:HB3 | 2:M:375:ILE:HD11 | 1.71 | 0.71 |
| 2:H:58:GLU:OE1 | 2:J:573:MET:HE1 | 1.90 | 0.71 |
| 2:L:372:LEU:HB3 | 2:L:375:ILE:HD11 | 1.71 | 0.71 |
| 2:H:372:LEU:HB3 | 2:H:375:ILE:HD11 | 1.71 | 0.71 |
| 2:H:34:GLU:OE2 | 2:J:383:ARG:HB2 | 1.90 | 0.71 |
| 2:K:372:LEU:HB3 | 2:K:375:ILE:HD11 | 1.71 | 0.71 |
| 1:A:334:ASP:HB3 | 1:A:337:HIS:CE1 | 2.26 | 0.71 |
| 2:I:64:PRO:HD2 | 2:I:69:ASN:HB3 | 1.72 | 0.71 |
| 2:J:123:LEU:HD21 | 2:J:133:ILE:HG13 | 1.71 | 0.71 |
| 2:J:64:PRO:HD2 | 2:J:69:ASN:HB3 | 1.72 | 0.71 |
| 2:I:123:LEU:HD21 | 2:I:133:ILE:HG13 | 1.71 | 0.71 |
| 2:I:586:VAL:HG23 | 2:I:599:LEU:HD21 | 1.73 | 0.71 |
| 2:J:586:VAL:HG23 | 2:J:599:LEU:HD21 | 1.73 | 0.71 |
| 2:L:529:ASN:OD1 | 2:L:532:LYS:HB3 | 1.91 | 0.71 |
| 2:M:529:ASN:OD1 | 2:M:532:LYS:HB3 | 1.91 | 0.71 |
| 1:D:176:GLU:CD | 1:D:198:LYS:HE3 | 2.12 | 0.70 |
| 1:D:334:ASP:HB3 | 1:D:337:HIS:CE1 | 2.26 | 0.70 |
| 1:A:176:GLU:CD | 1:A:198:LYS:HE3 | 2.12 | 0.70 |
| 1:B:501:LYS:HD3 | 1:D:420:GLY:O | 1.92 | 0.70 |
| 2:I:383:ARG:HB2 | 2:K:34:GLU:OE2 | 1.92 | 0.70 |
| 2:I:573:MET:HE1 | 2:K:58:GLU:OE1 | 1.90 | 0.70 |
| 1:F:176:GLU:CD | 1:F:198:LYS:HE3 | 2.12 | 0.70 |
| 1:B:139:PRO:HG3 | 1:B:144:ARG:NH1 | 2.06 | 0.70 |
| 1:C:139:PRO:HG3 | 1:C:144:ARG:NH1 | 2.06 | 0.70 |
| 1:E:176:GLU:CD | 1:E:198:LYS:HE3 | 2.12 | 0.70 |
| 2:J:58:GLU:OE1 | 2:L:573:MET:HE1 | 1.90 | 0.70 |
| 2:J:34:GLU:OE2 | 2:L:383:ARG:HB2 | 1.91 | 0.70 |
| 1:D:139:PRO:HG3 | 1:D:144:ARG:NH1 | 2.06 | 0.70 |
| 1:E:334:ASP:HB3 | 1:E:337:HIS:CE1 | 2.26 | 0.70 |
| 2:H:383:ARG:HB2 | 2:L:34:GLU:OE2 | 1.92 | 0.70 |
| 1:F:334:ASP:HB3 | 1:F:337:HIS:CE1 | 2.26 | 0.69 |
| 2:K:529:ASN:OD1 | 2:K:532:LYS:HB3 | 1.91 | 0.69 |
| 1:A:139:PRO:HG3 | 1:A:144:ARG:NH1 | 2.07 | 0.69 |
| 2:J:529:ASN:OD1 | 2:J:532:LYS:HB3 | 1.91 | 0.69 |
| 2:K:586:VAL:HG23 | 2:K:599:LEU:HD21 | 1.73 | 0.69 |

Continued on next page...

Not For Manuscript Review

*Continued from previous page...*

| Atom-1 | Atom-2 | Interatomic distance (Å) | Clash overlap (Å) |
|---|---|---|---|
| 1:F:139:PRO:HG3 | 1:F:144:ARG:NH1 | 2.06 | 0.69 |
| 2:I:529:ASN:OD1 | 2:I:532:LYS:HB3 | 1.91 | 0.69 |
| 1:E:139:PRO:HG3 | 1:E:144:ARG:NH1 | 2.06 | 0.69 |
| 2:I:34:GLU:OE2 | 2:M:383:ARG:HB2 | 1.92 | 0.69 |
| 2:J:58:GLU:OE1 | 2:L:573:MET:HE2 | 1.93 | 0.69 |
| 2:H:529:ASN:OD1 | 2:H:532:LYS:HB3 | 1.91 | 0.69 |
| 2:H:586:VAL:HG23 | 2:H:599:LEU:HD21 | 1.74 | 0.69 |
| 2:I:58:GLU:OE1 | 2:M:573:MET:HE1 | 1.90 | 0.69 |
| 2:L:586:VAL:HG23 | 2:L:599:LEU:HD21 | 1.73 | 0.69 |
| 2:M:586:VAL:HG23 | 2:M:599:LEU:HD21 | 1.73 | 0.69 |
| 2:L:562:VAL:HG22 | 2:L:580:VAL:HG13 | 1.74 | 0.69 |
| 2:M:562:VAL:HG22 | 2:M:580:VAL:HG13 | 1.74 | 0.69 |
| 1:C:176:GLU:CD | 1:C:198:LYS:HE3 | 2.12 | 0.69 |
| 2:L:225:PHE:HB2 | 2:L:235:PHE:HE1 | 1.58 | 0.69 |
| 1:F:233:ASP:OD1 | 1:F:435:ARG:NH1 | 2.24 | 0.69 |
| 2:J:562:VAL:HG22 | 2:J:580:VAL:HG13 | 1.74 | 0.69 |
| 2:K:383:ARG:HB2 | 2:M:34:GLU:OE2 | 1.93 | 0.69 |
| 2:M:225:PHE:HB2 | 2:M:235:PHE:HE1 | 1.58 | 0.68 |
| 1:E:233:ASP:OD1 | 1:E:435:ARG:NH1 | 2.24 | 0.68 |
| 2:I:225:PHE:HB2 | 2:I:235:PHE:HE1 | 1.58 | 0.68 |
| 1:B:176:GLU:CD | 1:B:198:LYS:HE3 | 2.12 | 0.68 |
| 2:H:58:GLU:OE1 | 2:J:573:MET:HE2 | 1.94 | 0.68 |
| 2:I:562:VAL:HG22 | 2:I:580:VAL:HG13 | 1.74 | 0.68 |
| 2:J:225:PHE:HB2 | 2:J:235:PHE:HE1 | 1.58 | 0.68 |
| 1:F:429:HIS:CD2 | 2:M:607:GLY:HA2 | 2.27 | 0.68 |
| 2:H:562:VAL:HG22 | 2:H:580:VAL:HG13 | 1.74 | 0.68 |
| 2:H:638:VAL:HG11 | 2:H:644:LEU:HD11 | 1.76 | 0.68 |
| 2:K:562:VAL:HG22 | 2:K:580:VAL:HG13 | 1.74 | 0.68 |
| 2:K:638:VAL:HG11 | 2:K:644:LEU:HD11 | 1.76 | 0.68 |
| 1:E:593:LEU:HD12 | 1:E:617:MET:HE2 | 1.76 | 0.68 |
| 1:F:593:LEU:HD12 | 1:F:617:MET:HE2 | 1.76 | 0.68 |
| 1:B:144:ARG:HG3 | 2:K:606:PHE:CD2 | 2.28 | 0.67 |
| 2:J:638:VAL:HG11 | 2:J:644:LEU:HD11 | 1.76 | 0.67 |
| 1:B:144:ARG:HG2 | 2:K:606:PHE:CE2 | 2.29 | 0.67 |
| 2:I:58:GLU:OE1 | 2:M:573:MET:HE2 | 1.94 | 0.67 |
| 2:I:638:VAL:HG11 | 2:I:644:LEU:HD11 | 1.76 | 0.67 |
| 2:H:254:PRO:HG3 | 2:H:315:VAL:HG21 | 1.76 | 0.67 |
| 2:K:225:PHE:HB2 | 2:K:235:PHE:HE1 | 1.58 | 0.67 |
| 1:A:564:LYS:HE3 | 1:B:406:GLU:HB3 | 1.75 | 0.67 |
| 2:H:225:PHE:HB2 | 2:H:235:PHE:HE1 | 1.58 | 0.67 |
| 2:I:573:MET:HE2 | 2:K:58:GLU:OE1 | 1.94 | 0.67 |

*Continued on next page...*

Not For Manuscript Review

*Continued from previous page...*

| Atom-1 | Atom-2 | Interatomic distance (Å) | Clash overlap (Å) |
|---|---|---|---|
| 2:K:254:PRO:HG3 | 2:K:315:VAL:HG21 | 1.76 | 0.67 |
| 1:A:406:GLU:HB3 | 1:B:564:LYS:HE3 | 1.77 | 0.67 |
| 1:C:564:LYS:HE3 | 1:D:406:GLU:HB3 | 1.76 | 0.67 |
| 2:J:254:PRO:HG3 | 2:J:315:VAL:HG21 | 1.76 | 0.67 |
| 2:M:254:PRO:HG3 | 2:M:315:VAL:HG21 | 1.75 | 0.67 |
| 1:C:593:LEU:HD12 | 1:C:617:MET:HE2 | 1.76 | 0.67 |
| 1:B:420:GLY:O | 1:F:501:LYS:HD3 | 1.94 | 0.67 |
| 2:L:638:VAL:HG11 | 2:L:644:LEU:HD11 | 1.76 | 0.67 |
| 2:M:638:VAL:HG11 | 2:M:644:LEU:HD11 | 1.76 | 0.67 |
| 2:I:254:PRO:HG3 | 2:I:315:VAL:HG21 | 1.76 | 0.67 |
| 2:K:573:MET:HE2 | 2:M:58:GLU:OE1 | 1.91 | 0.67 |
| 1:F:248:ASP:HB2 | 1:F:258:TYR:HD2 | 1.60 | 0.67 |
| 2:K:650:MET:CE | 2:K:651:LYS:HD3 | 1.78 | 0.67 |
| 2:L:254:PRO:HG3 | 2:L:315:VAL:HG21 | 1.76 | 0.67 |
| 1:B:593:LEU:HD12 | 1:B:617:MET:HE2 | 1.77 | 0.66 |
| 1:E:248:ASP:HB2 | 1:E:258:TYR:HD2 | 1.60 | 0.66 |
| 2:H:650:MET:CE | 2:H:651:LYS:HD3 | 1.78 | 0.66 |
| 1:B:248:ASP:HB2 | 1:B:258:TYR:HD2 | 1.60 | 0.66 |
| 1:C:420:GLY:O | 1:E:501:LYS:HD3 | 1.95 | 0.66 |
| 1:D:248:ASP:HB2 | 1:D:258:TYR:HD2 | 1.60 | 0.66 |
| 1:E:564:LYS:HE3 | 1:F:406:GLU:HB3 | 1.76 | 0.66 |
| 1:C:248:ASP:HB2 | 1:C:258:TYR:HD2 | 1.60 | 0.66 |
| 1:C:406:GLU:HB3 | 1:D:564:LYS:HE3 | 1.76 | 0.66 |
| 1:A:142:TYR:O | 1:A:144:ARG:N | 2.29 | 0.66 |
| 1:A:248:ASP:HB2 | 1:A:258:TYR:HD2 | 1.60 | 0.66 |
| 1:C:142:TYR:O | 1:C:144:ARG:N | 2.29 | 0.66 |
| 1:D:142:TYR:O | 1:D:144:ARG:N | 2.29 | 0.66 |
| 1:F:234:PRO:HB2 | 2:M:495:ASN:HD21 | 1.61 | 0.66 |
| 2:K:228:LYS:HG3 | 2:K:280:VAL:CG2 | 2.26 | 0.66 |
| 1:A:150:PRO:HB2 | 1:A:152:ILE:HG22 | 1.78 | 0.66 |
| 1:B:142:TYR:O | 1:B:144:ARG:N | 2.29 | 0.66 |
| 2:H:228:LYS:HG3 | 2:H:280:VAL:CG2 | 2.26 | 0.66 |
| 1:D:150:PRO:HB2 | 1:D:152:ILE:HG22 | 1.78 | 0.66 |
| 1:A:420:GLY:O | 1:C:501:LYS:HD3 | 1.96 | 0.66 |
| 1:F:150:PRO:HB2 | 1:F:152:ILE:HG22 | 1.78 | 0.66 |
| 1:E:150:PRO:HB2 | 1:E:152:ILE:HG22 | 1.78 | 0.65 |
| 1:A:664:GLU:HA | 1:A:669:TYR:CD1 | 2.32 | 0.65 |
| 1:B:207:GLU:OE1 | 1:B:211:ARG:NH2 | 2.29 | 0.65 |
| 1:E:207:GLU:OE1 | 1:E:211:ARG:NH2 | 2.29 | 0.65 |
| 1:F:142:TYR:O | 1:F:144:ARG:N | 2.29 | 0.65 |
| 1:F:207:GLU:OE1 | 1:F:211:ARG:NH2 | 2.29 | 0.65 |

*Continued on next page...*

Not For Manuscript Review

Continued from previous page...

| Atom-1 | Atom-2 | Interatomic distance (Å) | Clash overlap (Å) |
|---|---|---|---|
| 1:C:207:GLU:OE1 | 1:C:211:ARG:NH2 | 2.29 | 0.65 |
| 1:D:664:GLU:HA | 1:D:669:TYR:CD1 | 2.32 | 0.65 |
| 1:E:142:TYR:O | 1:E:144:ARG:N | 2.29 | 0.65 |
| 1:A:501:LYS:HD3 | 1:E:420:GLY:O | 1.96 | 0.65 |
| 1:D:397:ALA:HB2 | 2:H:650:MET:HG2 | 1.79 | 0.65 |
| 1:A:233:ASP:OD1 | 1:A:435:ARG:NH1 | 2.24 | 0.65 |
| 1:A:429:HIS:NE2 | 2:H:607:GLY:HA2 | 2.12 | 0.65 |
| 1:C:664:GLU:HA | 1:C:669:TYR:CD1 | 2.32 | 0.65 |
| 1:A:161:ASP:O | 1:A:165:GLN:HG2 | 1.97 | 0.65 |
| 1:B:608:ARG:HH21 | 1:B:615:LEU:HD13 | 1.62 | 0.65 |
| 1:B:664:GLU:HA | 1:B:669:TYR:CD1 | 2.32 | 0.65 |
| 1:C:608:ARG:HH21 | 1:C:615:LEU:HD13 | 1.62 | 0.65 |
| 1:D:161:ASP:O | 1:D:165:GLN:HG2 | 1.97 | 0.65 |
| 1:D:233:ASP:OD1 | 1:D:435:ARG:NH1 | 2.24 | 0.65 |
| 1:E:406:GLU:HB3 | 1:F:564:LYS:HE3 | 1.78 | 0.64 |
| 1:D:560:PHE:O | 3:G:803:BTI:C6 | 2.44 | 0.64 |
| 1:F:161:ASP:O | 1:F:165:GLN:HG2 | 1.97 | 0.64 |
| 2:H:225:PHE:HB2 | 2:H:235:PHE:CE1 | 2.33 | 0.64 |
| 2:H:378:PRO:HD3 | 2:H:432:TYR:HD1 | 1.63 | 0.64 |
| 2:K:225:PHE:HB2 | 2:K:235:PHE:CE1 | 2.33 | 0.64 |
| 2:K:378:PRO:HD3 | 2:K:432:TYR:HD1 | 1.63 | 0.64 |
| 1:A:593:LEU:HD12 | 1:A:617:MET:HE2 | 1.79 | 0.64 |
| 1:E:161:ASP:O | 1:E:165:GLN:HG2 | 1.97 | 0.64 |
| 1:F:664:GLU:HA | 1:F:669:TYR:CD1 | 2.32 | 0.64 |
| 2:L:228:LYS:HG3 | 2:L:280:VAL:CG2 | 2.26 | 0.64 |
| 1:B:161:ASP:O | 1:B:165:GLN:HG2 | 1.97 | 0.64 |
| 1:C:153:LEU:HA | 1:C:496:ARG:HA | 1.79 | 0.64 |
| 1:C:179:ARG:HA | 2:J:503:ASN:HB3 | 1.80 | 0.64 |
| 1:D:501:LYS:HD3 | 1:F:420:GLY:O | 1.96 | 0.64 |
| 1:E:664:GLU:HA | 1:E:669:TYR:CD1 | 2.32 | 0.64 |
| 2:M:228:LYS:HG3 | 2:M:280:VAL:CG2 | 2.26 | 0.64 |
| 1:A:608:ARG:HH21 | 1:A:615:LEU:HD13 | 1.62 | 0.64 |
| 1:B:153:LEU:HA | 1:B:496:ARG:HA | 1.79 | 0.64 |
| 1:E:630:GLN:HG2 | 3:G:802:BTI:O3 | 1.98 | 0.64 |
| 2:I:225:PHE:HB2 | 2:I:235:PHE:CE1 | 2.33 | 0.64 |
| 1:A:207:GLU:OE1 | 1:A:211:ARG:NH2 | 2.29 | 0.64 |
| 1:B:397:ALA:HB2 | 2:L:650:MET:HG2 | 1.79 | 0.64 |
| 1:C:161:ASP:O | 1:C:165:GLN:HG2 | 1.97 | 0.64 |
| 1:D:608:ARG:HH21 | 1:D:615:LEU:HD13 | 1.62 | 0.64 |
| 1:E:179:ARG:HA | 2:L:503:ASN:HB3 | 1.80 | 0.64 |
| 1:E:608:ARG:HH21 | 1:E:615:LEU:HD13 | 1.62 | 0.64 |

Continued on next page...

Not For Manuscript Review

Continued from previous page...

| Atom-1 | Atom-2 | Interatomic distance (Å) | Clash overlap (Å) |
|---|---|---|---|
| 1:F:608:ARG:HH21 | 1:F:615:LEU:HD13 | 1.62 | 0.64 |
| 2:J:225:PHE:HB2 | 2:J:235:PHE:CE1 | 2.33 | 0.64 |
| 2:L:225:PHE:HB2 | 2:L:235:PHE:CE1 | 2.33 | 0.64 |
| 1:B:150:PRO:HB2 | 1:B:152:ILE:HG22 | 1.77 | 0.64 |
| 1:D:153:LEU:HA | 1:D:496:ARG:HA | 1.79 | 0.64 |
| 1:A:153:LEU:HA | 1:A:496:ARG:HA | 1.79 | 0.64 |
| 1:D:207:GLU:OE1 | 1:D:211:ARG:NH2 | 2.29 | 0.64 |
| 2:I:378:PRO:HD3 | 2:I:432:TYR:HD1 | 1.63 | 0.64 |
| 2:M:225:PHE:HB2 | 2:M:235:PHE:CE1 | 2.33 | 0.64 |
| 1:C:150:PRO:HB2 | 1:C:152:ILE:HG22 | 1.78 | 0.63 |
| 1:C:233:ASP:OD1 | 1:C:435:ARG:NH1 | 2.24 | 0.63 |
| 2:J:378:PRO:HD3 | 2:J:432:TYR:HD1 | 1.63 | 0.63 |
| 1:E:153:LEU:HA | 1:E:496:ARG:HA | 1.79 | 0.63 |
| 2:M:378:PRO:HD3 | 2:M:432:TYR:HD1 | 1.63 | 0.63 |
| 2:L:378:PRO:HD3 | 2:L:432:TYR:HD1 | 1.63 | 0.63 |
| 1:F:153:LEU:HA | 1:F:496:ARG:HA | 1.79 | 0.63 |
| 1:B:233:ASP:OD1 | 1:B:435:ARG:NH1 | 2.24 | 0.63 |
| 1:D:593:LEU:HD12 | 1:D:617:MET:HE2 | 1.80 | 0.63 |
| 1:F:397:ALA:HB2 | 2:J:650:MET:HG2 | 1.81 | 0.63 |
| 1:A:364:THR:HG22 | 1:A:387:PHE:HB2 | 1.81 | 0.63 |
| 1:B:364:THR:HG22 | 1:B:387:PHE:HB2 | 1.81 | 0.62 |
| 1:D:179:ARG:HA | 2:K:503:ASN:HB3 | 1.81 | 0.62 |
| 1:D:364:THR:HG22 | 1:D:387:PHE:HB2 | 1.81 | 0.62 |
| 1:A:179:ARG:HA | 2:H:503:ASN:HB3 | 1.81 | 0.62 |
| 1:C:364:THR:HG22 | 1:C:387:PHE:HB2 | 1.81 | 0.62 |
| 2:M:233:VAL:CG2 | 2:M:342:GLN:HA | 2.30 | 0.62 |
| 1:B:237:ARG:NH1 | 1:B:691:GLU:OE2 | 2.32 | 0.62 |
| 1:C:237:ARG:NH1 | 1:C:691:GLU:OE2 | 2.33 | 0.62 |
| 1:F:237:ARG:NH1 | 1:F:691:GLU:OE2 | 2.32 | 0.62 |
| 2:L:233:VAL:CG2 | 2:L:342:GLN:HA | 2.30 | 0.62 |
| 1:B:144:ARG:CG | 2:K:606:PHE:CD2 | 2.82 | 0.62 |
| 1:E:142:TYR:C | 1:E:145:SER:H | 2.03 | 0.62 |
| 1:E:237:ARG:NH1 | 1:E:691:GLU:OE2 | 2.33 | 0.62 |
| 2:J:228:LYS:HG3 | 2:J:280:VAL:CG2 | 2.26 | 0.62 |
| 1:F:142:TYR:C | 1:F:145:SER:H | 2.03 | 0.62 |
| 2:I:228:LYS:HG3 | 2:I:280:VAL:CG2 | 2.26 | 0.62 |
| 2:K:236:PHE:CD2 | 2:K:316:CYS:HB3 | 2.35 | 0.62 |
| 2:H:236:PHE:CD2 | 2:H:316:CYS:HB3 | 2.35 | 0.62 |
| 1:B:466:GLU:OE1 | 1:B:490:ARG:NH2 | 2.27 | 0.62 |
| 1:B:669:TYR:HA | 1:B:673:ARG:NE | 2.13 | 0.62 |
| 2:H:233:VAL:CG2 | 2:H:342:GLN:HA | 2.30 | 0.62 |

Continued on next page...

Not For Manuscript Review

*Continued from previous page...*

| Atom-1 | Atom-2 | Interatomic distance (Å) | Clash overlap (Å) |
|---|---|---|---|
| 2:J:314:GLU:C | 2:J:317:ARG:HH12 | 2.03 | 0.62 |
| 2:K:233:VAL:CG2 | 2:K:342:GLN:HA | 2.30 | 0.62 |
| 2:M:32:CYS:SG | 2:M:37:ILE:HB | 2.40 | 0.62 |
| 1:C:142:TYR:C | 1:C:145:SER:H | 2.03 | 0.61 |
| 1:C:429:HIS:NE2 | 2:J:607:GLY:HA2 | 2.15 | 0.61 |
| 1:C:669:TYR:HA | 1:C:673:ARG:NE | 2.13 | 0.61 |
| 2:I:314:GLU:C | 2:I:317:ARG:HH12 | 2.03 | 0.61 |
| 2:K:160:LYS:HG3 | 2:K:164:PHE:CE1 | 2.35 | 0.61 |
| 2:L:32:CYS:SG | 2:L:37:ILE:HB | 2.40 | 0.61 |
| 2:L:236:PHE:CD2 | 2:L:316:CYS:HB3 | 2.35 | 0.61 |
| 2:L:314:GLU:C | 2:L:317:ARG:HH12 | 2.03 | 0.61 |
| 2:M:314:GLU:C | 2:M:317:ARG:HH12 | 2.03 | 0.61 |
| 1:D:142:TYR:C | 1:D:145:SER:H | 2.02 | 0.61 |
| 1:F:251:TRP:HZ2 | 1:F:255:LYS:HG3 | 1.65 | 0.61 |
| 2:H:160:LYS:HG3 | 2:H:164:PHE:CE1 | 2.35 | 0.61 |
| 2:I:233:VAL:CG2 | 2:I:342:GLN:HA | 2.30 | 0.61 |
| 2:M:236:PHE:CD2 | 2:M:316:CYS:HB3 | 2.35 | 0.61 |
| 1:C:251:TRP:HZ2 | 1:C:255:LYS:HG3 | 1.65 | 0.61 |
| 1:C:466:GLU:OE1 | 1:C:490:ARG:NH2 | 2.27 | 0.61 |
| 1:D:669:TYR:HA | 1:D:673:ARG:NE | 2.13 | 0.61 |
| 1:A:237:ARG:NH1 | 1:A:691:GLU:OE2 | 2.32 | 0.61 |
| 1:A:251:TRP:HZ2 | 1:A:255:LYS:HG3 | 1.65 | 0.61 |
| 1:E:251:TRP:HZ2 | 1:E:255:LYS:HG3 | 1.65 | 0.61 |
| 2:H:314:GLU:C | 2:H:317:ARG:HH12 | 2.03 | 0.61 |
| 2:I:32:CYS:SG | 2:I:37:ILE:HB | 2.40 | 0.61 |
| 2:I:236:PHE:CD2 | 2:I:316:CYS:HB3 | 2.35 | 0.61 |
| 2:J:233:VAL:CG2 | 2:J:342:GLN:HA | 2.30 | 0.61 |
| 1:A:142:TYR:C | 1:A:145:SER:H | 2.03 | 0.61 |
| 1:B:142:TYR:C | 1:B:145:SER:H | 2.03 | 0.61 |
| 1:B:144:ARG:CG | 2:K:606:PHE:CE2 | 2.82 | 0.61 |
| 1:B:251:TRP:HZ2 | 1:B:255:LYS:HG3 | 1.65 | 0.61 |
| 1:E:429:HIS:NE2 | 2:L:607:GLY:HA2 | 2.15 | 0.61 |
| 2:I:160:LYS:HG3 | 2:I:164:PHE:CE1 | 2.35 | 0.61 |
| 1:A:669:TYR:HA | 1:A:673:ARG:NE | 2.13 | 0.61 |
| 1:D:237:ARG:NH1 | 1:D:691:GLU:OE2 | 2.33 | 0.61 |
| 1:D:251:TRP:HZ2 | 1:D:255:LYS:HG3 | 1.65 | 0.61 |
| 2:J:32:CYS:SG | 2:J:37:ILE:HB | 2.41 | 0.61 |
| 2:J:160:LYS:HG3 | 2:J:164:PHE:CE1 | 2.35 | 0.61 |
| 2:J:236:PHE:CD2 | 2:J:316:CYS:HB3 | 2.35 | 0.61 |
| 2:K:314:GLU:C | 2:K:317:ARG:HH12 | 2.04 | 0.61 |
| 1:E:177:GLN:HG2 | 1:E:198:LYS:HG3 | 1.83 | 0.61 |

*Continued on next page...*

Not For Manuscript Review

Continued from previous page...

| Atom-1 | Atom-2 | Interatomic distance (Å) | Clash overlap (Å) |
|---|---|---|---|
| 1:F:177:GLN:HG2 | 1:F:198:LYS:HG3 | 1.82 | 0.61 |
| 2:L:93:TYR:HA | 2:L:304:ARG:HD2 | 1.83 | 0.61 |
| 2:L:160:LYS:HG3 | 2:L:164:PHE:CE1 | 2.35 | 0.61 |
| 2:L:160:LYS:HG3 | 2:L:164:PHE:HE1 | 1.66 | 0.61 |
| 2:M:93:TYR:HA | 2:M:304:ARG:HD2 | 1.83 | 0.61 |
| 2:L:487:ALA:O | 2:L:491:ASN:HB2 | 2.01 | 0.61 |
| 2:M:160:LYS:HG3 | 2:M:164:PHE:CE1 | 2.35 | 0.61 |
| 2:M:487:ALA:O | 2:M:491:ASN:HB2 | 2.01 | 0.61 |
| 2:M:160:LYS:HG3 | 2:M:164:PHE:HE1 | 1.66 | 0.61 |
| 2:H:93:TYR:HA | 2:H:304:ARG:HD2 | 1.83 | 0.60 |
| 2:J:650:MET:HE3 | 2:J:650:MET:C | 2.20 | 0.60 |
| 2:K:32:CYS:SG | 2:K:37:ILE:HB | 2.40 | 0.60 |
| 2:K:93:TYR:HA | 2:K:304:ARG:HD2 | 1.83 | 0.60 |
| 2:H:32:CYS:SG | 2:H:37:ILE:HB | 2.40 | 0.60 |
| 2:I:93:TYR:HA | 2:I:304:ARG:HD2 | 1.83 | 0.60 |
| 2:K:496:TRP:CH2 | 2:K:524:HIS:HA | 2.36 | 0.60 |
| 1:B:554:LEU:HD22 | 1:B:594:ILE:HD11 | 1.83 | 0.60 |
| 1:C:554:LEU:HD22 | 1:C:594:ILE:HD11 | 1.83 | 0.60 |
| 1:D:177:GLN:HG2 | 1:D:198:LYS:HG3 | 1.83 | 0.60 |
| 1:F:364:THR:HG22 | 1:F:387:PHE:HB2 | 1.81 | 0.60 |
| 2:I:116:PRO:HG3 | 2:I:303:THR:HB | 1.84 | 0.60 |
| 2:I:650:MET:HE3 | 2:I:650:MET:C | 2.20 | 0.60 |
| 2:J:93:TYR:HA | 2:J:304:ARG:HD2 | 1.83 | 0.60 |
| 2:J:116:PRO:HG3 | 2:J:303:THR:HB | 1.84 | 0.60 |
| 1:A:150:PRO:O | 1:A:153:LEU:HG | 2.02 | 0.60 |
| 1:E:364:THR:HG22 | 1:E:387:PHE:HB2 | 1.81 | 0.60 |
| 2:H:496:TRP:CH2 | 2:H:524:HIS:HA | 2.36 | 0.60 |
| 1:A:177:GLN:HG2 | 1:A:198:LYS:HG3 | 1.83 | 0.60 |
| 1:B:177:GLN:HG2 | 1:B:198:LYS:HG3 | 1.83 | 0.60 |
| 2:L:496:TRP:CH2 | 2:L:524:HIS:HA | 2.36 | 0.60 |
| 2:M:496:TRP:CH2 | 2:M:524:HIS:HA | 2.36 | 0.60 |
| 1:D:150:PRO:O | 1:D:153:LEU:HG | 2.02 | 0.60 |
| 1:E:150:PRO:O | 1:E:153:LEU:HG | 2.02 | 0.60 |
| 1:F:560:PHE:O | 3:G:801:BTI:C6 | 2.49 | 0.60 |
| 2:I:98:GLU:OE2 | 2:I:304:ARG:NH1 | 2.35 | 0.60 |
| 2:M:98:GLU:OE2 | 2:M:304:ARG:NH1 | 2.35 | 0.60 |
| 1:B:150:PRO:O | 1:B:153:LEU:HG | 2.02 | 0.60 |
| 1:C:150:PRO:O | 1:C:153:LEU:HG | 2.02 | 0.60 |
| 1:C:177:GLN:HG2 | 1:C:198:LYS:HG3 | 1.83 | 0.60 |
| 1:F:150:PRO:O | 1:F:153:LEU:HG | 2.02 | 0.60 |
| 2:H:573:MET:HE2 | 2:L:58:GLU:OE1 | 1.99 | 0.60 |

Continued on next page...

Not For Manuscript Review

*Continued from previous page...*

| Atom-1 | Atom-2 | Interatomic distance (Å) | Clash overlap (Å) |
|---|---|---|---|
| 2:J:98:GLU:OE2 | 2:J:304:ARG:NH1 | 2.35 | 0.60 |
| 2:K:487:ALA:O | 2:K:491:ASN:HB2 | 2.01 | 0.60 |
| 2:L:98:GLU:OE2 | 2:L:304:ARG:NH1 | 2.35 | 0.60 |
| 2:M:494:ASP:HB2 | 2:M:497:ARG:HB2 | 1.84 | 0.60 |
| 1:D:687:VAL:O | 1:D:691:GLU:HG2 | 2.02 | 0.60 |
| 2:H:98:GLU:OE2 | 2:H:304:ARG:NH1 | 2.35 | 0.60 |
| 2:K:98:GLU:OE2 | 2:K:304:ARG:NH1 | 2.35 | 0.60 |
| 2:L:494:ASP:HB2 | 2:L:497:ARG:HB2 | 1.84 | 0.60 |
| 1:E:669:TYR:HA | 1:E:673:ARG:NE | 2.13 | 0.60 |
| 1:F:466:GLU:OE1 | 1:F:490:ARG:NH2 | 2.27 | 0.60 |
| 1:A:687:VAL:O | 1:A:691:GLU:HG2 | 2.02 | 0.60 |
| 1:C:687:VAL:O | 1:C:691:GLU:HG2 | 2.02 | 0.60 |
| 1:F:669:TYR:HA | 1:F:673:ARG:NE | 2.13 | 0.60 |
| 2:I:487:ALA:O | 2:I:491:ASN:HB2 | 2.01 | 0.60 |
| 2:J:494:ASP:HB2 | 2:J:497:ARG:HB2 | 1.84 | 0.60 |
| 1:B:687:VAL:O | 1:B:691:GLU:HG2 | 2.02 | 0.59 |
| 2:H:116:PRO:HG3 | 2:H:303:THR:HB | 1.84 | 0.59 |
| 2:H:487:ALA:O | 2:H:491:ASN:HB2 | 2.01 | 0.59 |
| 2:I:494:ASP:HB2 | 2:I:497:ARG:HB2 | 1.84 | 0.59 |
| 2:J:487:ALA:O | 2:J:491:ASN:HB2 | 2.01 | 0.59 |
| 2:K:116:PRO:HG3 | 2:K:303:THR:HB | 1.84 | 0.59 |
| 2:L:116:PRO:HG3 | 2:L:303:THR:HB | 1.83 | 0.59 |
| 1:B:429:HIS:CD2 | 2:I:607:GLY:HA2 | 2.36 | 0.59 |
| 1:E:466:GLU:OE1 | 1:E:490:ARG:NH2 | 2.27 | 0.59 |
| 2:J:160:LYS:HG3 | 2:J:164:PHE:HE1 | 1.66 | 0.59 |
| 1:B:152:ILE:HG12 | 1:B:496:ARG:HB3 | 1.83 | 0.59 |
| 2:M:116:PRO:HG3 | 2:M:303:THR:HB | 1.84 | 0.59 |
| 1:E:152:ILE:HG12 | 1:E:496:ARG:HB3 | 1.84 | 0.59 |
| 1:E:554:LEU:HD22 | 1:E:594:ILE:HD11 | 1.83 | 0.59 |
| 1:F:152:ILE:HG12 | 1:F:496:ARG:HB3 | 1.84 | 0.59 |
| 2:I:160:LYS:HG3 | 2:I:164:PHE:HE1 | 1.66 | 0.59 |
| 2:J:496:TRP:CH2 | 2:J:524:HIS:HA | 2.36 | 0.59 |
| 1:D:554:LEU:HD22 | 1:D:594:ILE:HD11 | 1.83 | 0.59 |
| 1:F:137:HIS:CD2 | 2:I:602:LEU:HD21 | 2.38 | 0.59 |
| 1:C:152:ILE:HG12 | 1:C:496:ARG:HB3 | 1.83 | 0.59 |
| 1:E:176:GLU:HB3 | 1:E:198:LYS:NZ | 2.18 | 0.59 |
| 1:F:554:LEU:HD22 | 1:F:594:ILE:HD11 | 1.83 | 0.59 |
| 1:A:554:LEU:HD22 | 1:A:594:ILE:HD11 | 1.83 | 0.59 |
| 1:C:505:GLY:O | 1:C:536:LYS:NZ | 2.36 | 0.59 |
| 1:B:505:GLY:O | 1:B:536:LYS:NZ | 2.36 | 0.59 |
| 1:C:176:GLU:HB3 | 1:C:198:LYS:NZ | 2.18 | 0.59 |

*Continued on next page...*

Not For Manuscript Review

*Continued from previous page...*

| Atom-1 | Atom-2 | Interatomic distance (Å) | Clash overlap (Å) |
|---|---|---|---|
| 1:F:176:GLU:HB3 | 1:F:198:LYS:NZ | 2.18 | 0.59 |
| 2:I:496:TRP:CH2 | 2:I:524:HIS:HA | 2.37 | 0.59 |
| 2:K:160:LYS:HG3 | 2:K:164:PHE:HE1 | 1.66 | 0.59 |
| 1:B:176:GLU:HB3 | 1:B:198:LYS:NZ | 2.18 | 0.59 |
| 1:D:466:GLU:OE1 | 1:D:490:ARG:NH2 | 2.27 | 0.59 |
| 1:A:505:GLY:O | 1:A:536:LYS:NZ | 2.36 | 0.59 |
| 1:D:505:GLY:O | 1:D:536:LYS:NZ | 2.36 | 0.59 |
| 2:H:262:THR:HG23 | 2:H:265:ARG:NH2 | 2.18 | 0.59 |
| 2:M:262:THR:HG23 | 2:M:265:ARG:NH2 | 2.18 | 0.59 |
| 1:D:478:VAL:HG23 | 2:M:651:LYS:CB | 2.33 | 0.58 |
| 1:E:687:VAL:O | 1:E:691:GLU:HG2 | 2.02 | 0.58 |
| 2:H:160:LYS:HG3 | 2:H:164:PHE:HE1 | 1.66 | 0.58 |
| 2:J:262:THR:HG23 | 2:J:265:ARG:NH2 | 2.18 | 0.58 |
| 2:K:262:THR:HG23 | 2:K:265:ARG:NH2 | 2.18 | 0.58 |
| 2:K:494:ASP:HB2 | 2:K:497:ARG:HB2 | 1.84 | 0.58 |
| 1:D:152:ILE:HG12 | 1:D:496:ARG:HB3 | 1.84 | 0.58 |
| 1:F:505:GLY:O | 1:F:536:LYS:NZ | 2.36 | 0.58 |
| 2:I:262:THR:HG23 | 2:I:265:ARG:NH2 | 2.18 | 0.58 |
| 2:J:413:SER:HB3 | 2:J:425:LEU:HG | 1.86 | 0.58 |
| 2:L:262:THR:HG23 | 2:L:265:ARG:NH2 | 2.18 | 0.58 |
| 1:A:152:ILE:HG12 | 1:A:496:ARG:HB3 | 1.84 | 0.58 |
| 1:A:466:GLU:OE1 | 1:A:490:ARG:NH2 | 2.27 | 0.58 |
| 1:B:234:PRO:HD2 | 1:F:142:TYR:CZ | 2.37 | 0.58 |
| 1:D:537:GLY:O | 1:D:541:ILE:HG13 | 2.03 | 0.58 |
| 1:E:505:GLY:O | 1:E:536:LYS:NZ | 2.36 | 0.58 |
| 1:F:687:VAL:O | 1:F:691:GLU:HG2 | 2.02 | 0.58 |
| 2:H:494:ASP:HB2 | 2:H:497:ARG:HB2 | 1.84 | 0.58 |
| 2:H:315:VAL:HG22 | 2:H:347:LEU:HB2 | 1.86 | 0.58 |
| 2:H:496:TRP:HH2 | 2:H:524:HIS:HA | 1.69 | 0.58 |
| 2:K:315:VAL:HG22 | 2:K:347:LEU:HB2 | 1.86 | 0.58 |
| 1:A:537:GLY:O | 1:A:541:ILE:HG13 | 2.04 | 0.58 |
| 1:F:537:GLY:O | 1:F:541:ILE:HG13 | 2.04 | 0.58 |
| 2:H:20:ASN:ND2 | 2:H:93:TYR:O | 2.37 | 0.58 |
| 2:I:413:SER:HB3 | 2:I:425:LEU:HG | 1.86 | 0.58 |
| 2:K:20:ASN:ND2 | 2:K:93:TYR:O | 2.37 | 0.58 |
| 2:K:496:TRP:HH2 | 2:K:524:HIS:HA | 1.69 | 0.58 |
| 2:L:20:ASN:ND2 | 2:L:93:TYR:O | 2.37 | 0.58 |
| 2:M:20:ASN:ND2 | 2:M:93:TYR:O | 2.37 | 0.58 |
| 1:A:608:ARG:HG3 | 1:B:335:GLU:HB2 | 1.86 | 0.58 |
| 1:B:142:TYR:CZ | 1:D:234:PRO:HD2 | 2.38 | 0.58 |
| 1:E:537:GLY:O | 1:E:541:ILE:HG13 | 2.04 | 0.58 |

*Continued on next page...*

Not For Manuscript Review

*Continued from previous page...*

| Atom-1 | Atom-2 | Interatomic distance (Å) | Clash overlap (Å) |
|---|---|---|---|
| 2:J:496:TRP:HH2 | 2:J:524:HIS:HA | 1.68 | 0.58 |
| 2:L:219:HIS:O | 2:L:220:ILE:N | 2.37 | 0.58 |
| 2:M:219:HIS:O | 2:M:220:ILE:N | 2.37 | 0.58 |
| 1:E:231:LEU:O | 1:E:435:ARG:NH2 | 2.37 | 0.58 |
| 1:D:144:ARG:HG2 | 2:M:606:PHE:CE2 | 2.37 | 0.58 |
| 1:F:162:PRO:O | 1:F:166:GLU:HG2 | 2.04 | 0.58 |
| 1:F:231:LEU:O | 1:F:435:ARG:NH2 | 2.37 | 0.58 |
| 2:I:496:TRP:HH2 | 2:I:524:HIS:HA | 1.69 | 0.58 |
| 1:D:176:GLU:HB3 | 1:D:198:LYS:NZ | 2.18 | 0.58 |
| 1:D:234:PRO:HB2 | 2:K:495:ASN:HD21 | 1.68 | 0.58 |
| 1:E:162:PRO:O | 1:E:166:GLU:HG2 | 2.04 | 0.58 |
| 2:I:20:ASN:ND2 | 2:I:93:TYR:O | 2.37 | 0.58 |
| 2:J:20:ASN:ND2 | 2:J:93:TYR:O | 2.37 | 0.58 |
| 2:L:126:SER:HB3 | 2:L:129:GLU:HG2 | 1.86 | 0.58 |
| 1:A:504:TYR:HE1 | 1:E:415:ARG:N | 2.02 | 0.57 |
| 1:B:231:LEU:O | 1:B:435:ARG:NH2 | 2.37 | 0.57 |
| 1:C:162:PRO:O | 1:C:166:GLU:HG2 | 2.04 | 0.57 |
| 1:E:139:PRO:HG3 | 1:E:144:ARG:HH12 | 1.69 | 0.57 |
| 2:K:413:SER:HB3 | 2:K:425:LEU:HG | 1.86 | 0.57 |
| 2:L:448:CYS:HA | 2:L:453:PHE:HE1 | 1.69 | 0.57 |
| 2:M:126:SER:HB3 | 2:M:129:GLU:HG2 | 1.86 | 0.57 |
| 1:C:231:LEU:O | 1:C:435:ARG:NH2 | 2.37 | 0.57 |
| 1:D:231:LEU:O | 1:D:435:ARG:NH2 | 2.37 | 0.57 |
| 2:H:413:SER:HB3 | 2:H:425:LEU:HG | 1.86 | 0.57 |
| 1:A:231:LEU:O | 1:A:435:ARG:NH2 | 2.37 | 0.57 |
| 1:B:162:PRO:O | 1:B:166:GLU:HG2 | 2.05 | 0.57 |
| 1:F:139:PRO:HG3 | 1:F:144:ARG:HH12 | 1.69 | 0.57 |
| 2:H:191:MET:SD | 2:H:192:LEU:N | 2.78 | 0.57 |
| 2:H:219:HIS:O | 2:H:220:ILE:N | 2.37 | 0.57 |
| 2:K:219:HIS:O | 2:K:220:ILE:N | 2.37 | 0.57 |
| 2:L:413:SER:HB3 | 2:L:425:LEU:HG | 1.85 | 0.57 |
| 2:L:620:VAL:HB | 2:L:677:ASP:HA | 1.86 | 0.57 |
| 2:M:448:CYS:HA | 2:M:453:PHE:HE1 | 1.69 | 0.57 |
| 1:A:162:PRO:O | 1:A:166:GLU:HG2 | 2.04 | 0.57 |
| 1:A:176:GLU:HB3 | 1:A:198:LYS:NZ | 2.18 | 0.57 |
| 1:D:162:PRO:O | 1:D:166:GLU:HG2 | 2.05 | 0.57 |
| 2:I:191:MET:SD | 2:I:192:LEU:N | 2.78 | 0.57 |
| 2:J:191:MET:SD | 2:J:192:LEU:N | 2.78 | 0.57 |
| 2:K:191:MET:SD | 2:K:192:LEU:N | 2.78 | 0.57 |
| 2:M:620:VAL:HB | 2:M:677:ASP:HA | 1.86 | 0.57 |
| 1:A:266:ILE:HD11 | 1:A:299:LYS:HB3 | 1.86 | 0.57 |

*Continued on next page...*

Not For Manuscript Review

Continued from previous page...

| Atom-1 | Atom-2 | Interatomic distance (Å) | Clash overlap (Å) |
|---|---|---|---|
| 1:D:142:TYR:CZ | 1:F:234:PRO:HD2 | 2.39 | 0.57 |
| 2:H:126:SER:HB3 | 2:H:129:GLU:HG2 | 1.86 | 0.57 |
| 2:K:126:SER:HB3 | 2:K:129:GLU:HG2 | 1.86 | 0.57 |
| 1:A:615:LEU:HB3 | 1:A:676:ASP:OD2 | 2.04 | 0.57 |
| 2:H:258:LEU:HB3 | 2:H:263:ARG:HE | 1.69 | 0.57 |
| 2:I:315:VAL:HG22 | 2:I:347:LEU:HB2 | 1.86 | 0.57 |
| 2:M:315:VAL:HG22 | 2:M:347:LEU:HB2 | 1.86 | 0.57 |
| 1:B:537:GLY:O | 1:B:541:ILE:HG13 | 2.04 | 0.57 |
| 1:C:234:PRO:HD2 | 1:E:142:TYR:CZ | 2.39 | 0.57 |
| 1:D:615:LEU:HB3 | 1:D:676:ASP:OD2 | 2.04 | 0.57 |
| 2:J:126:SER:HB3 | 2:J:129:GLU:HG2 | 1.86 | 0.57 |
| 2:M:413:SER:HB3 | 2:M:425:LEU:HG | 1.86 | 0.57 |
| 2:M:496:TRP:HH2 | 2:M:524:HIS:HA | 1.68 | 0.57 |
| 1:C:537:GLY:O | 1:C:541:ILE:HG13 | 2.04 | 0.57 |
| 1:D:139:PRO:HG3 | 1:D:144:ARG:HH12 | 1.69 | 0.57 |
| 1:D:144:ARG:CG | 2:M:606:PHE:CD2 | 2.88 | 0.57 |
| 1:D:266:ILE:HD11 | 1:D:299:LYS:HB3 | 1.86 | 0.57 |
| 1:F:266:ILE:HD11 | 1:F:299:LYS:HB3 | 1.86 | 0.57 |
| 2:I:217:SER:OG | 2:I:460:THR:OG1 | 2.23 | 0.57 |
| 2:I:258:LEU:HB3 | 2:I:263:ARG:HE | 1.69 | 0.57 |
| 2:J:258:LEU:HB3 | 2:J:263:ARG:HE | 1.69 | 0.57 |
| 2:J:315:VAL:HG22 | 2:J:347:LEU:HB2 | 1.86 | 0.57 |
| 2:K:258:LEU:HB3 | 2:K:263:ARG:HE | 1.69 | 0.57 |
| 2:L:315:VAL:HG22 | 2:L:347:LEU:HB2 | 1.86 | 0.57 |
| 2:L:496:TRP:HH2 | 2:L:524:HIS:HA | 1.69 | 0.57 |
| 2:M:191:MET:SD | 2:M:192:LEU:N | 2.78 | 0.57 |
| 1:E:266:ILE:HD11 | 1:E:299:LYS:HB3 | 1.86 | 0.57 |
| 2:I:448:CYS:HA | 2:I:453:PHE:HE1 | 1.69 | 0.57 |
| 2:L:191:MET:SD | 2:L:192:LEU:N | 2.78 | 0.57 |
| 2:I:219:HIS:O | 2:I:220:ILE:N | 2.37 | 0.57 |
| 2:L:258:LEU:HB3 | 2:L:263:ARG:HE | 1.69 | 0.57 |
| 1:A:139:PRO:HG3 | 1:A:144:ARG:HH12 | 1.69 | 0.56 |
| 1:A:317:LEU:HD13 | 1:A:358:VAL:HB | 1.87 | 0.56 |
| 1:D:317:LEU:HD13 | 1:D:358:VAL:HB | 1.88 | 0.56 |
| 1:F:615:LEU:HB3 | 1:F:676:ASP:OD2 | 2.04 | 0.56 |
| 2:H:448:CYS:HA | 2:H:453:PHE:HE1 | 1.69 | 0.56 |
| 2:J:219:HIS:O | 2:J:220:ILE:N | 2.37 | 0.56 |
| 2:J:448:CYS:HA | 2:J:453:PHE:HE1 | 1.69 | 0.56 |
| 1:A:193:GLU:HG3 | 1:A:195:GLU:H | 1.70 | 0.56 |
| 1:A:335:GLU:HB2 | 1:B:608:ARG:HG3 | 1.87 | 0.56 |
| 1:D:504:TYR:HE1 | 1:F:415:ARG:N | 2.03 | 0.56 |

Continued on next page...

Not For Manuscript Review

*Continued from previous page...*

| Atom-1 | Atom-2 | Interatomic distance (Å) | Clash overlap (Å) |
|---|---|---|---|
| 1:E:608:ARG:HG3 | 1:F:335:GLU:HB2 | 1.87 | 0.56 |
| 1:E:615:LEU:HB3 | 1:E:676:ASP:OD2 | 2.04 | 0.56 |
| 2:I:126:SER:HB3 | 2:I:129:GLU:HG2 | 1.86 | 0.56 |
| 1:B:617:MET:HB2 | 1:B:676:ASP:OD1 | 2.05 | 0.56 |
| 1:C:335:GLU:HB2 | 1:D:608:ARG:HG3 | 1.88 | 0.56 |
| 1:C:617:MET:HB2 | 1:C:676:ASP:OD1 | 2.05 | 0.56 |
| 2:I:314:GLU:HG2 | 2:I:414:TRP:HB2 | 1.87 | 0.56 |
| 2:J:314:GLU:HG2 | 2:J:414:TRP:HB2 | 1.87 | 0.56 |
| 2:J:600:LEU:HD12 | 2:J:601:PRO:HD2 | 1.87 | 0.56 |
| 2:K:448:CYS:HA | 2:K:453:PHE:HE1 | 1.69 | 0.56 |
| 2:L:605:GLY:O | 2:L:606:PHE:C | 2.43 | 0.56 |
| 2:M:258:LEU:HB3 | 2:M:263:ARG:HE | 1.70 | 0.56 |
| 2:M:605:GLY:O | 2:M:606:PHE:C | 2.43 | 0.56 |
| 1:B:266:ILE:HD11 | 1:B:299:LYS:HB3 | 1.86 | 0.56 |
| 1:D:193:GLU:HG3 | 1:D:195:GLU:H | 1.70 | 0.56 |
| 1:E:335:GLU:HB2 | 1:F:608:ARG:HG3 | 1.88 | 0.56 |
| 1:C:608:ARG:HG3 | 1:D:335:GLU:HB2 | 1.88 | 0.56 |
| 1:C:615:LEU:HB3 | 1:C:676:ASP:OD2 | 2.04 | 0.56 |
| 1:D:560:PHE:O | 3:G:803:BTI:H62 | 2.04 | 0.56 |
| 1:E:397:ALA:HB2 | 2:I:650:MET:HG2 | 1.87 | 0.56 |
| 1:E:617:MET:HB2 | 1:E:676:ASP:OD1 | 2.05 | 0.56 |
| 1:F:560:PHE:O | 3:G:801:BTI:H62 | 2.06 | 0.56 |
| 2:I:600:LEU:HD12 | 2:I:601:PRO:HD2 | 1.87 | 0.56 |
| 1:A:142:TYR:CZ | 1:E:234:PRO:HD2 | 2.40 | 0.56 |
| 1:B:135:TYR:CE2 | 1:D:220:GLY:HA2 | 2.41 | 0.56 |
| 1:B:615:LEU:HB3 | 1:B:676:ASP:OD2 | 2.04 | 0.56 |
| 1:C:317:LEU:HD13 | 1:C:358:VAL:HB | 1.87 | 0.56 |
| 1:D:413:HIS:CE1 | 1:D:415:ARG:HB3 | 2.40 | 0.56 |
| 2:I:418:ARG:HH12 | 2:I:454:ALA:HA | 1.71 | 0.56 |
| 2:M:217:SER:OG | 2:M:460:THR:OG1 | 2.23 | 0.56 |
| 1:F:617:MET:HB2 | 1:F:676:ASP:OD1 | 2.05 | 0.56 |
| 2:J:418:ARG:HH12 | 2:J:454:ALA:HA | 1.71 | 0.56 |
| 2:J:620:VAL:HB | 2:J:677:ASP:HA | 1.86 | 0.56 |
| 2:K:620:VAL:HB | 2:K:677:ASP:HA | 1.86 | 0.56 |
| 2:L:217:SER:OG | 2:L:460:THR:OG1 | 2.23 | 0.56 |
| 1:A:413:HIS:CE1 | 1:A:415:ARG:HB3 | 2.40 | 0.56 |
| 1:B:283:ASN:HD22 | 1:B:318:VAL:HA | 1.71 | 0.56 |
| 1:B:317:LEU:HD13 | 1:B:358:VAL:HB | 1.88 | 0.56 |
| 1:B:413:HIS:CE1 | 1:B:415:ARG:HB3 | 2.40 | 0.56 |
| 1:C:413:HIS:CE1 | 1:C:415:ARG:HB3 | 2.40 | 0.56 |
| 1:F:137:HIS:HD2 | 2:I:602:LEU:HD21 | 1.69 | 0.56 |

*Continued on next page...*

Not For Manuscript Review

*Continued from previous page...*

| Atom-1 | Atom-2 | Interatomic distance (Å) | Clash overlap (Å) |
|---|---|---|---|
| 2:I:620:VAL:HB | 2:I:677:ASP:HA | 1.86 | 0.56 |
| 1:A:415:ARG:N | 1:C:504:TYR:HE1 | 2.04 | 0.56 |
| 1:B:144:ARG:HD2 | 2:K:606:PHE:CG | 2.40 | 0.56 |
| 1:C:193:GLU:HG3 | 1:C:195:GLU:H | 1.70 | 0.56 |
| 1:C:266:ILE:HD11 | 1:C:299:LYS:HB3 | 1.86 | 0.56 |
| 1:C:283:ASN:HD22 | 1:C:318:VAL:HA | 1.71 | 0.56 |
| 1:F:413:HIS:CE1 | 1:F:415:ARG:HB3 | 2.40 | 0.56 |
| 2:L:600:LEU:HD12 | 2:L:601:PRO:HD2 | 1.86 | 0.56 |
| 1:A:617:MET:HB2 | 1:A:676:ASP:OD1 | 2.05 | 0.56 |
| 1:D:144:ARG:HG3 | 2:M:606:PHE:CD2 | 2.40 | 0.56 |
| 1:E:413:HIS:CE1 | 1:E:415:ARG:HB3 | 2.40 | 0.56 |
| 2:H:600:LEU:HD12 | 2:H:601:PRO:HD2 | 1.87 | 0.56 |
| 2:H:620:VAL:HB | 2:H:677:ASP:HA | 1.86 | 0.56 |
| 2:M:600:LEU:HD12 | 2:M:601:PRO:HD2 | 1.87 | 0.56 |
| 1:A:266:ILE:HG22 | 1:A:283:ASN:HA | 1.88 | 0.55 |
| 1:B:193:GLU:HG3 | 1:B:195:GLU:H | 1.70 | 0.55 |
| 2:L:314:GLU:HG2 | 2:L:414:TRP:HB2 | 1.87 | 0.55 |
| 1:A:283:ASN:HD22 | 1:A:318:VAL:HA | 1.71 | 0.55 |
| 1:C:266:ILE:HG22 | 1:C:283:ASN:HA | 1.88 | 0.55 |
| 1:D:266:ILE:HG22 | 1:D:283:ASN:HA | 1.88 | 0.55 |
| 1:D:617:MET:HB2 | 1:D:676:ASP:OD1 | 2.06 | 0.55 |
| 2:I:376:ARG:NH1 | 2:I:376:ARG:HA | 2.21 | 0.55 |
| 2:I:605:GLY:O | 2:I:606:PHE:C | 2.43 | 0.55 |
| 2:K:600:LEU:HD12 | 2:K:601:PRO:HD2 | 1.87 | 0.55 |
| 2:M:314:GLU:HG2 | 2:M:414:TRP:HB2 | 1.87 | 0.55 |
| 1:B:266:ILE:HG22 | 1:B:283:ASN:HA | 1.88 | 0.55 |
| 1:C:139:PRO:HG3 | 1:C:144:ARG:HH12 | 1.69 | 0.55 |
| 1:C:415:ARG:N | 1:E:504:TYR:HE1 | 2.03 | 0.55 |
| 1:E:193:GLU:HG3 | 1:E:195:GLU:H | 1.70 | 0.55 |
| 2:H:314:GLU:HG2 | 2:H:414:TRP:HB2 | 1.87 | 0.55 |
| 2:J:17:LEU:HB2 | 2:J:86:VAL:HG11 | 1.88 | 0.55 |
| 2:J:376:ARG:HA | 2:J:376:ARG:NH1 | 2.21 | 0.55 |
| 2:M:418:ARG:HH12 | 2:M:454:ALA:HA | 1.71 | 0.55 |
| 1:D:283:ASN:HD22 | 1:D:318:VAL:HA | 1.71 | 0.55 |
| 1:E:283:ASN:HD22 | 1:E:318:VAL:HA | 1.71 | 0.55 |
| 2:H:605:GLY:O | 2:H:606:PHE:C | 2.43 | 0.55 |
| 2:J:605:GLY:O | 2:J:606:PHE:C | 2.43 | 0.55 |
| 2:K:314:GLU:HG2 | 2:K:414:TRP:HB2 | 1.87 | 0.55 |
| 2:K:605:GLY:O | 2:K:606:PHE:C | 2.43 | 0.55 |
| 1:A:234:PRO:HD2 | 1:C:142:TYR:CZ | 2.41 | 0.55 |
| 1:E:266:ILE:HG22 | 1:E:283:ASN:HA | 1.88 | 0.55 |

*Continued on next page...*

Not For Manuscript Review

Continued from previous page...

| Atom-1 | Atom-2 | Interatomic distance (Å) | Clash overlap (Å) |
|---|---|---|---|
| 1:E:317:LEU:HD13 | 1:E:358:VAL:HB | 1.87 | 0.55 |
| 1:F:193:GLU:HG3 | 1:F:195:GLU:H | 1.70 | 0.55 |
| 2:I:17:LEU:HB2 | 2:I:86:VAL:HG11 | 1.88 | 0.55 |
| 2:L:418:ARG:HH12 | 2:L:454:ALA:HA | 1.71 | 0.55 |
| 1:A:416:ILE:HD11 | 1:B:568:GLU:OE1 | 2.06 | 0.55 |
| 1:A:630:GLN:HG2 | 3:G:806:BTI:O3 | 2.07 | 0.55 |
| 1:B:139:PRO:HG3 | 1:B:144:ARG:HH12 | 1.69 | 0.55 |
| 1:F:266:ILE:HG22 | 1:F:283:ASN:HA | 1.88 | 0.55 |
| 1:F:283:ASN:HD22 | 1:F:318:VAL:HA | 1.71 | 0.55 |
| 1:F:317:LEU:HD13 | 1:F:358:VAL:HB | 1.87 | 0.55 |
| 2:H:226:PHE:CD1 | 2:H:279:TYR:HD2 | 2.25 | 0.55 |
| 2:L:376:ARG:NH1 | 2:L:376:ARG:HA | 2.21 | 0.55 |
| 2:H:418:ARG:HH12 | 2:H:454:ALA:HA | 1.71 | 0.55 |
| 2:K:226:PHE:CD1 | 2:K:279:TYR:HD2 | 2.25 | 0.55 |
| 2:M:376:ARG:NH1 | 2:M:376:ARG:HA | 2.21 | 0.55 |
| 1:D:478:VAL:HG23 | 2:M:651:LYS:HB3 | 1.89 | 0.55 |
| 2:K:383:ARG:CD | 2:M:34:GLU:OE2 | 2.49 | 0.55 |
| 2:K:418:ARG:HH12 | 2:K:454:ALA:HA | 1.71 | 0.55 |
| 1:E:641:ASN:HB3 | 1:E:653:PHE:HE2 | 1.72 | 0.55 |
| 2:H:34:GLU:OE2 | 2:J:383:ARG:CD | 2.48 | 0.55 |
| 2:J:636:ASP:HA | 2:J:663:ARG:HH11 | 1.71 | 0.55 |
| 2:K:376:ARG:NH1 | 2:K:376:ARG:HA | 2.21 | 0.55 |
| 1:B:143:GLU:N | 1:B:145:SER:O | 2.40 | 0.55 |
| 1:F:641:ASN:HB3 | 1:F:653:PHE:HE2 | 1.72 | 0.55 |
| 2:I:636:ASP:HA | 2:I:663:ARG:HH11 | 1.71 | 0.55 |
| 1:A:525:ASN:HD21 | 1:A:529:LEU:HD11 | 1.72 | 0.54 |
| 1:B:142:TYR:C | 1:B:144:ARG:N | 2.60 | 0.54 |
| 1:C:142:TYR:C | 1:C:144:ARG:N | 2.60 | 0.54 |
| 1:C:143:GLU:N | 1:C:145:SER:O | 2.40 | 0.54 |
| 1:B:525:ASN:HD21 | 1:B:529:LEU:HD11 | 1.72 | 0.54 |
| 1:C:525:ASN:HD21 | 1:C:529:LEU:HD11 | 1.72 | 0.54 |
| 2:H:17:LEU:HB2 | 2:H:86:VAL:HG11 | 1.88 | 0.54 |
| 2:L:44:CYS:SG | 2:L:45:GLU:N | 2.80 | 0.54 |
| 1:B:144:ARG:HD2 | 2:K:606:PHE:CD1 | 2.42 | 0.54 |
| 1:B:478:VAL:HG23 | 2:K:651:LYS:HA | 1.88 | 0.54 |
| 1:C:428:LEU:HD11 | 1:E:139:PRO:HD2 | 1.89 | 0.54 |
| 2:H:44:CYS:SG | 2:H:45:GLU:N | 2.80 | 0.54 |
| 2:H:217:SER:OG | 2:H:460:THR:OG1 | 2.23 | 0.54 |
| 2:H:376:ARG:NH1 | 2:H:376:ARG:HA | 2.21 | 0.54 |
| 2:K:44:CYS:SG | 2:K:45:GLU:N | 2.80 | 0.54 |
| 2:L:17:LEU:HB2 | 2:L:86:VAL:HG11 | 1.88 | 0.54 |

Continued on next page...

Not For Manuscript Review

*Continued from previous page...*

| Atom-1 | Atom-2 | Interatomic distance (Å) | Clash overlap (Å) |
|---|---|---|---|
| 2:M:17:LEU:HB2 | 2:M:86:VAL:HG11 | 1.88 | 0.54 |
| 1:C:641:ASN:HB3 | 1:C:653:PHE:HE2 | 1.72 | 0.54 |
| 1:F:177:GLN:HA | 1:F:180:ALA:HB3 | 1.90 | 0.54 |
| 2:H:636:ASP:HA | 2:H:663:ARG:HH11 | 1.71 | 0.54 |
| 2:J:621:SER:HB3 | 2:J:647:VAL:HG21 | 1.90 | 0.54 |
| 2:K:17:LEU:HB2 | 2:K:86:VAL:HG11 | 1.88 | 0.54 |
| 2:M:44:CYS:SG | 2:M:45:GLU:N | 2.80 | 0.54 |
| 1:E:177:GLN:HA | 1:E:180:ALA:HB3 | 1.90 | 0.54 |
| 1:F:525:ASN:HD21 | 1:F:529:LEU:HD11 | 1.72 | 0.54 |
| 2:H:383:ARG:CD | 2:L:34:GLU:OE2 | 2.49 | 0.54 |
| 1:B:248:ASP:HB2 | 1:B:258:TYR:CD2 | 2.42 | 0.54 |
| 1:B:641:ASN:HB3 | 1:B:653:PHE:HE2 | 1.72 | 0.54 |
| 1:D:144:ARG:HD2 | 2:M:606:PHE:CG | 2.43 | 0.54 |
| 1:D:525:ASN:HD21 | 1:D:529:LEU:HD11 | 1.73 | 0.54 |
| 2:K:217:SER:OG | 2:K:460:THR:OG1 | 2.23 | 0.54 |
| 2:K:636:ASP:HA | 2:K:663:ARG:HH11 | 1.72 | 0.54 |
| 2:M:636:ASP:HA | 2:M:663:ARG:HH11 | 1.71 | 0.54 |
| 1:A:142:TYR:C | 1:A:144:ARG:N | 2.60 | 0.54 |
| 1:B:504:TYR:HE1 | 1:D:415:ARG:N | 2.06 | 0.54 |
| 1:E:524:ALA:HB2 | 1:E:554:LEU:HD12 | 1.90 | 0.54 |
| 2:I:621:SER:HB3 | 2:I:647:VAL:HG21 | 1.90 | 0.54 |
| 1:C:248:ASP:HB2 | 1:C:258:TYR:CD2 | 2.42 | 0.54 |
| 1:D:142:TYR:C | 1:D:144:ARG:N | 2.60 | 0.54 |
| 1:E:525:ASN:HD21 | 1:E:529:LEU:HD11 | 1.72 | 0.54 |
| 1:F:524:ALA:HB2 | 1:F:554:LEU:HD12 | 1.90 | 0.54 |
| 2:I:491:ASN:O | 2:I:561:ILE:HD13 | 2.08 | 0.54 |
| 2:J:226:PHE:CD1 | 2:J:279:TYR:HD2 | 2.25 | 0.54 |
| 2:J:491:ASN:O | 2:J:561:ILE:HD13 | 2.08 | 0.54 |
| 2:L:636:ASP:HA | 2:L:663:ARG:HH11 | 1.71 | 0.54 |
| 2:M:226:PHE:CD1 | 2:M:279:TYR:HD2 | 2.25 | 0.54 |
| 2:M:621:SER:HB3 | 2:M:647:VAL:HG21 | 1.90 | 0.54 |
| 1:A:641:ASN:HB3 | 1:A:653:PHE:HE2 | 1.72 | 0.54 |
| 1:B:415:ARG:N | 1:F:504:TYR:HE1 | 2.05 | 0.54 |
| 1:E:143:GLU:N | 1:E:145:SER:O | 2.40 | 0.54 |
| 2:L:226:PHE:CD1 | 2:L:279:TYR:HD2 | 2.25 | 0.54 |
| 2:L:621:SER:HB3 | 2:L:647:VAL:HG21 | 1.90 | 0.54 |
| 1:A:139:PRO:HD2 | 1:E:428:LEU:HD11 | 1.89 | 0.54 |
| 1:B:139:PRO:HD2 | 1:D:428:LEU:HD11 | 1.88 | 0.54 |
| 1:D:137:HIS:HD2 | 2:M:602:LEU:HD21 | 1.71 | 0.54 |
| 1:D:284:ASP:OD1 | 1:D:285:ALA:N | 2.41 | 0.54 |
| 1:D:641:ASN:HB3 | 1:D:653:PHE:HE2 | 1.72 | 0.54 |

*Continued on next page...*

Not For Manuscript Review

*Continued from previous page...*

| Atom-1 | Atom-2 | Interatomic distance (Å) | Clash overlap (Å) |
|---|---|---|---|
| 2:K:262:THR:O | 2:K:266:ILE:HG13 | 2.08 | 0.54 |
| 1:A:284:ASP:OD1 | 1:A:285:ALA:N | 2.41 | 0.53 |
| 1:A:524:ALA:HB2 | 1:A:554:LEU:HD12 | 1.90 | 0.53 |
| 1:C:177:GLN:HA | 1:C:180:ALA:HB3 | 1.90 | 0.53 |
| 1:F:143:GLU:N | 1:F:145:SER:O | 2.40 | 0.53 |
| 2:H:262:THR:O | 2:H:266:ILE:HG13 | 2.08 | 0.53 |
| 2:I:226:PHE:CD1 | 2:I:279:TYR:HD2 | 2.25 | 0.53 |
| 2:L:262:THR:O | 2:L:266:ILE:HG13 | 2.08 | 0.53 |
| 1:A:154:ARG:HH22 | 1:A:497:PHE:HB3 | 1.73 | 0.53 |
| 1:B:154:ARG:HH22 | 1:B:497:PHE:HB3 | 1.73 | 0.53 |
| 1:D:154:ARG:HH22 | 1:D:497:PHE:HB3 | 1.73 | 0.53 |
| 1:D:524:ALA:HB2 | 1:D:554:LEU:HD12 | 1.90 | 0.53 |
| 1:E:154:ARG:HH22 | 1:E:497:PHE:HB3 | 1.73 | 0.53 |
| 1:F:154:ARG:HH22 | 1:F:497:PHE:HB3 | 1.73 | 0.53 |
| 1:B:177:GLN:HA | 1:B:180:ALA:HB3 | 1.90 | 0.53 |
| 1:D:177:GLN:HA | 1:D:180:ALA:HB3 | 1.90 | 0.53 |
| 2:H:621:SER:HB3 | 2:H:647:VAL:HG21 | 1.90 | 0.53 |
| 2:M:262:THR:O | 2:M:266:ILE:HG13 | 2.08 | 0.53 |
| 1:A:143:GLU:N | 1:A:145:SER:O | 2.40 | 0.53 |
| 1:A:177:GLN:HA | 1:A:180:ALA:HB3 | 1.90 | 0.53 |
| 1:A:529:LEU:N | 1:A:559:GLY:O | 2.42 | 0.53 |
| 1:B:428:LEU:HD11 | 1:F:139:PRO:HD2 | 1.90 | 0.53 |
| 1:C:154:ARG:HH22 | 1:C:497:PHE:HB3 | 1.73 | 0.53 |
| 1:C:170:ARG:O | 1:C:174:LEU:HG | 2.08 | 0.53 |
| 1:C:524:ALA:HB2 | 1:C:554:LEU:HD12 | 1.90 | 0.53 |
| 2:K:621:SER:HB3 | 2:K:647:VAL:HG21 | 1.90 | 0.53 |
| 2:M:233:VAL:HG23 | 2:M:342:GLN:HA | 1.91 | 0.53 |
| 1:B:524:ALA:HB2 | 1:B:554:LEU:HD12 | 1.90 | 0.53 |
| 1:D:143:GLU:N | 1:D:145:SER:O | 2.40 | 0.53 |
| 1:D:529:LEU:N | 1:D:559:GLY:O | 2.42 | 0.53 |
| 1:E:142:TYR:C | 1:E:144:ARG:N | 2.60 | 0.53 |
| 1:F:160:SER:HA | 1:F:164:TYR:CD2 | 2.44 | 0.53 |
| 1:B:170:ARG:O | 1:B:174:LEU:HG | 2.08 | 0.53 |
| 1:C:161:ASP:HB3 | 1:C:162:PRO:HD2 | 1.91 | 0.53 |
| 1:E:160:SER:HA | 1:E:164:TYR:CD2 | 2.44 | 0.53 |
| 2:J:262:THR:O | 2:J:266:ILE:HG13 | 2.08 | 0.53 |
| 2:M:654:HIS:ND1 | 2:M:654:HIS:O | 2.42 | 0.53 |
| 1:B:179:ARG:HB3 | 1:B:179:ARG:NH1 | 2.24 | 0.53 |
| 1:D:179:ARG:NH1 | 1:D:179:ARG:HB3 | 2.24 | 0.53 |
| 1:E:529:LEU:N | 1:E:559:GLY:O | 2.42 | 0.53 |
| 1:F:142:TYR:C | 1:F:144:ARG:N | 2.60 | 0.53 |

*Continued on next page...*

Not For Manuscript Review

Continued from previous page...

| Atom-1 | Atom-2 | Interatomic distance (Å) | Clash overlap (Å) |
|---|---|---|---|
| 2:H:491:ASN:O | 2:H:561:ILE:HD13 | 2.08 | 0.53 |
| 2:I:262:THR:O | 2:I:266:ILE:HG13 | 2.08 | 0.53 |
| 2:L:233:VAL:HG23 | 2:L:342:GLN:HA | 1.91 | 0.53 |
| 2:L:653:GLU:O | 2:L:654:HIS:C | 2.47 | 0.53 |
| 2:L:654:HIS:ND1 | 2:L:654:HIS:O | 2.42 | 0.53 |
| 2:M:653:GLU:O | 2:M:654:HIS:C | 2.47 | 0.53 |
| 1:B:179:ARG:HA | 2:I:503:ASN:HB3 | 1.90 | 0.53 |
| 2:I:44:CYS:SG | 2:I:45:GLU:N | 2.80 | 0.53 |
| 1:A:179:ARG:NH1 | 1:A:179:ARG:HB3 | 2.24 | 0.53 |
| 1:C:179:ARG:HB3 | 1:C:179:ARG:NH1 | 2.24 | 0.53 |
| 1:D:170:ARG:O | 1:D:174:LEU:HG | 2.08 | 0.53 |
| 1:D:283:ASN:ND2 | 1:D:318:VAL:HA | 2.24 | 0.53 |
| 1:E:416:ILE:HD11 | 1:F:568:GLU:OE1 | 2.08 | 0.53 |
| 2:J:44:CYS:SG | 2:J:45:GLU:N | 2.80 | 0.53 |
| 2:J:654:HIS:ND1 | 2:J:654:HIS:O | 2.42 | 0.53 |
| 2:K:491:ASN:O | 2:K:561:ILE:HD13 | 2.08 | 0.53 |
| 2:M:236:PHE:HD2 | 2:M:316:CYS:HB3 | 1.74 | 0.53 |
| 1:A:135:TYR:CE2 | 1:E:220:GLY:HA2 | 2.44 | 0.53 |
| 1:A:161:ASP:HB3 | 1:A:162:PRO:HD2 | 1.91 | 0.53 |
| 1:B:161:ASP:HB3 | 1:B:162:PRO:HD2 | 1.91 | 0.53 |
| 1:D:161:ASP:HB3 | 1:D:162:PRO:HD2 | 1.91 | 0.53 |
| 1:E:284:ASP:OD1 | 1:E:285:ALA:N | 2.41 | 0.53 |
| 1:F:358:VAL:HG22 | 1:F:378:ILE:HD12 | 1.91 | 0.53 |
| 2:I:233:VAL:HG23 | 2:I:342:GLN:HA | 1.91 | 0.53 |
| 2:I:654:HIS:ND1 | 2:I:654:HIS:O | 2.42 | 0.53 |
| 1:A:170:ARG:O | 1:A:174:LEU:HG | 2.08 | 0.52 |
| 1:A:283:ASN:ND2 | 1:A:318:VAL:HA | 2.25 | 0.52 |
| 1:E:358:VAL:HG22 | 1:E:378:ILE:HD12 | 1.91 | 0.52 |
| 1:F:284:ASP:OD1 | 1:F:285:ALA:N | 2.41 | 0.52 |
| 2:H:236:PHE:HD2 | 2:H:316:CYS:HB3 | 1.74 | 0.52 |
| 2:J:233:VAL:HG23 | 2:J:342:GLN:HA | 1.91 | 0.52 |
| 2:K:233:VAL:HG23 | 2:K:342:GLN:HA | 1.91 | 0.52 |
| 2:K:261:GLU:HG3 | 2:K:262:THR:N | 2.25 | 0.52 |
| 2:L:236:PHE:HD2 | 2:L:316:CYS:HB3 | 1.74 | 0.52 |
| 2:M:491:ASN:O | 2:M:561:ILE:HD13 | 2.08 | 0.52 |
| 1:B:220:GLY:HA2 | 1:F:135:TYR:CE2 | 2.44 | 0.52 |
| 1:C:284:ASP:OD1 | 1:C:285:ALA:N | 2.41 | 0.52 |
| 2:H:261:GLU:HG3 | 2:H:262:THR:N | 2.25 | 0.52 |
| 2:I:100:ALA:HB1 | 2:I:118:ALA:HB1 | 1.92 | 0.52 |
| 2:K:236:PHE:HD2 | 2:K:316:CYS:HB3 | 1.74 | 0.52 |
| 2:L:491:ASN:O | 2:L:561:ILE:HD13 | 2.08 | 0.52 |

Continued on next page...

Not For Manuscript Review

*Continued from previous page...*

| Atom-1 | Atom-2 | Interatomic distance (Å) | Clash overlap (Å) |
|---|---|---|---|
| 1:A:160:SER:HA | 1:A:164:TYR:CD2 | 2.44 | 0.52 |
| 1:A:190:VAL:HA | 1:A:206:ILE:HG23 | 1.91 | 0.52 |
| 1:B:284:ASP:OD1 | 1:B:285:ALA:N | 2.42 | 0.52 |
| 1:C:220:GLY:HA2 | 1:E:135:TYR:CE2 | 2.44 | 0.52 |
| 1:D:190:VAL:HA | 1:D:206:ILE:HG23 | 1.91 | 0.52 |
| 1:D:358:VAL:HG22 | 1:D:378:ILE:HD12 | 1.91 | 0.52 |
| 1:E:170:ARG:O | 1:E:174:LEU:HG | 2.08 | 0.52 |
| 1:E:288:LYS:H | 1:E:288:LYS:HD2 | 1.74 | 0.52 |
| 1:F:248:ASP:HB2 | 1:F:258:TYR:CD2 | 2.42 | 0.52 |
| 1:F:288:LYS:H | 1:F:288:LYS:HD2 | 1.74 | 0.52 |
| 2:K:653:GLU:O | 2:K:654:HIS:C | 2.47 | 0.52 |
| 1:A:358:VAL:HG22 | 1:A:378:ILE:HD12 | 1.91 | 0.52 |
| 1:B:170:ARG:O | 1:B:173:GLY:N | 2.43 | 0.52 |
| 1:D:137:HIS:CD2 | 2:M:602:LEU:HD21 | 2.44 | 0.52 |
| 1:E:248:ASP:HB2 | 1:E:258:TYR:CD2 | 2.42 | 0.52 |
| 1:F:170:ARG:O | 1:F:174:LEU:HG | 2.08 | 0.52 |
| 2:H:653:GLU:O | 2:H:654:HIS:C | 2.48 | 0.52 |
| 2:J:100:ALA:HB1 | 2:J:118:ALA:HB1 | 1.92 | 0.52 |
| 1:C:160:SER:HA | 1:C:164:TYR:CD2 | 2.44 | 0.52 |
| 1:C:170:ARG:O | 1:C:173:GLY:N | 2.43 | 0.52 |
| 1:D:160:SER:HA | 1:D:164:TYR:CD2 | 2.44 | 0.52 |
| 1:E:170:ARG:O | 1:E:173:GLY:N | 2.43 | 0.52 |
| 1:F:179:ARG:NH1 | 1:F:179:ARG:HB3 | 2.24 | 0.52 |
| 2:H:654:HIS:ND1 | 2:H:654:HIS:O | 2.42 | 0.52 |
| 2:K:654:HIS:ND1 | 2:K:654:HIS:O | 2.42 | 0.52 |
| 1:A:170:ARG:O | 1:A:173:GLY:N | 2.43 | 0.52 |
| 1:B:160:SER:HA | 1:B:164:TYR:CD2 | 2.44 | 0.52 |
| 1:C:358:VAL:HG22 | 1:C:378:ILE:HD12 | 1.91 | 0.52 |
| 1:D:170:ARG:O | 1:D:173:GLY:N | 2.43 | 0.52 |
| 2:H:233:VAL:HG23 | 2:H:342:GLN:HA | 1.91 | 0.52 |
| 1:C:190:VAL:HA | 1:C:206:ILE:HG23 | 1.91 | 0.52 |
| 1:C:416:ILE:HD11 | 1:D:568:GLU:OE1 | 2.10 | 0.52 |
| 1:E:179:ARG:HB3 | 1:E:179:ARG:NH1 | 2.24 | 0.52 |
| 1:F:170:ARG:O | 1:F:173:GLY:N | 2.43 | 0.52 |
| 2:H:223:GLN:HG2 | 2:H:235:PHE:CD1 | 2.44 | 0.52 |
| 2:I:223:GLN:HG2 | 2:I:235:PHE:CD1 | 2.44 | 0.52 |
| 2:L:223:GLN:HG2 | 2:L:235:PHE:CD1 | 2.44 | 0.52 |
| 1:A:178:LEU:O | 1:A:182:VAL:HG23 | 2.10 | 0.52 |
| 1:A:226:ASP:HA | 1:A:229:GLU:OE2 | 2.10 | 0.52 |
| 1:A:248:ASP:HB2 | 1:A:258:TYR:CD2 | 2.42 | 0.52 |
| 1:F:161:ASP:HB3 | 1:F:162:PRO:HD2 | 1.91 | 0.52 |

*Continued on next page...*

Not For Manuscript Review

Continued from previous page...

| Atom-1 | Atom-2 | Interatomic distance (Å) | Clash overlap (Å) |
|---|---|---|---|
| 1:F:283:ASN:ND2 | 1:F:318:VAL:HA | 2.24 | 0.52 |
| 1:A:328:GLN:HG3 | 1:B:625:VAL:HG12 | 1.92 | 0.52 |
| 1:B:190:VAL:HA | 1:B:206:ILE:HG23 | 1.91 | 0.52 |
| 1:B:358:VAL:HG22 | 1:B:378:ILE:HD12 | 1.91 | 0.52 |
| 1:B:397:ALA:CB | 2:L:650:MET:HG2 | 2.40 | 0.52 |
| 1:D:226:ASP:HA | 1:D:229:GLU:OE2 | 2.10 | 0.52 |
| 1:D:248:ASP:HB2 | 1:D:258:TYR:CD2 | 2.42 | 0.52 |
| 1:E:283:ASN:ND2 | 1:E:318:VAL:HA | 2.25 | 0.52 |
| 2:H:100:ALA:HB1 | 2:H:118:ALA:HB1 | 1.92 | 0.52 |
| 2:J:653:GLU:O | 2:J:654:HIS:C | 2.47 | 0.52 |
| 2:K:100:ALA:HB1 | 2:K:118:ALA:HB1 | 1.92 | 0.52 |
| 2:K:223:GLN:HG2 | 2:K:235:PHE:CD1 | 2.45 | 0.52 |
| 2:M:100:ALA:HB1 | 2:M:118:ALA:HB1 | 1.92 | 0.52 |
| 1:A:576:ALA:HB2 | 1:B:372:ALA:HB1 | 1.91 | 0.52 |
| 1:B:226:ASP:HA | 1:B:229:GLU:OE2 | 2.10 | 0.52 |
| 1:D:178:LEU:O | 1:D:182:VAL:HG23 | 2.10 | 0.52 |
| 1:E:161:ASP:HB3 | 1:E:162:PRO:HD2 | 1.91 | 0.52 |
| 2:I:568:LEU:HD12 | 2:I:573:MET:O | 2.10 | 0.52 |
| 2:I:653:GLU:O | 2:I:654:HIS:C | 2.48 | 0.52 |
| 2:L:100:ALA:HB1 | 2:L:118:ALA:HB1 | 1.92 | 0.52 |
| 2:M:223:GLN:HG2 | 2:M:235:PHE:CD1 | 2.44 | 0.52 |
| 1:B:178:LEU:O | 1:B:182:VAL:HG23 | 2.10 | 0.51 |
| 1:B:529:LEU:N | 1:B:559:GLY:O | 2.42 | 0.51 |
| 1:C:178:LEU:O | 1:C:182:VAL:HG23 | 2.10 | 0.51 |
| 1:C:226:ASP:HA | 1:C:229:GLU:OE2 | 2.10 | 0.51 |
| 1:C:568:GLU:OE1 | 1:D:416:ILE:HD11 | 2.09 | 0.51 |
| 2:J:223:GLN:HG2 | 2:J:235:PHE:CD1 | 2.45 | 0.51 |
| 2:J:236:PHE:HD2 | 2:J:316:CYS:HB3 | 1.74 | 0.51 |
| 1:A:568:GLU:OE1 | 1:B:416:ILE:HD11 | 2.10 | 0.51 |
| 1:B:560:PHE:O | 3:G:805:BTI:C6 | 2.58 | 0.51 |
| 2:J:568:LEU:HD12 | 2:J:573:MET:O | 2.11 | 0.51 |
| 1:B:191:VAL:HG12 | 1:B:193:GLU:HG2 | 1.93 | 0.51 |
| 1:B:283:ASN:ND2 | 1:B:318:VAL:HA | 2.24 | 0.51 |
| 1:C:191:VAL:HG12 | 1:C:193:GLU:HG2 | 1.93 | 0.51 |
| 1:C:377:ASN:ND2 | 1:C:420:GLY:HA2 | 2.26 | 0.51 |
| 1:C:529:LEU:N | 1:C:559:GLY:O | 2.42 | 0.51 |
| 1:D:139:PRO:HD2 | 1:F:428:LEU:HD11 | 1.91 | 0.51 |
| 1:D:191:VAL:HG12 | 1:D:193:GLU:HG2 | 1.93 | 0.51 |
| 1:E:190:VAL:HA | 1:E:206:ILE:HG23 | 1.91 | 0.51 |
| 1:E:568:GLU:OE1 | 1:F:416:ILE:HD11 | 2.09 | 0.51 |
| 2:J:217:SER:OG | 2:J:460:THR:OG1 | 2.23 | 0.51 |

Continued on next page...

Not For Manuscript Review

Continued from previous page...

| Atom-1 | Atom-2 | Interatomic distance (Å) | Clash overlap (Å) |
|---|---|---|---|
| 2:M:261:GLU:HG3 | 2:M:262:THR:N | 2.25 | 0.51 |
| 1:A:191:VAL:HG12 | 1:A:193:GLU:HG2 | 1.93 | 0.51 |
| 1:A:288:LYS:HD2 | 1:A:288:LYS:H | 1.75 | 0.51 |
| 1:B:377:ASN:ND2 | 1:B:420:GLY:HA2 | 2.26 | 0.51 |
| 1:D:288:LYS:H | 1:D:288:LYS:HD2 | 1.75 | 0.51 |
| 1:F:190:VAL:HA | 1:F:206:ILE:HG23 | 1.91 | 0.51 |
| 2:I:236:PHE:HD2 | 2:I:316:CYS:HB3 | 1.74 | 0.51 |
| 2:L:261:GLU:HG3 | 2:L:262:THR:N | 2.25 | 0.51 |
| 2:M:568:LEU:HD12 | 2:M:573:MET:O | 2.11 | 0.51 |
| 1:B:504:TYR:CE2 | 1:D:423:TYR:HE2 | 2.29 | 0.51 |
| 1:C:283:ASN:ND2 | 1:C:318:VAL:HA | 2.25 | 0.51 |
| 1:E:576:ALA:HB2 | 1:F:372:ALA:HB1 | 1.92 | 0.51 |
| 2:L:568:LEU:HD12 | 2:L:573:MET:O | 2.11 | 0.51 |
| 1:A:220:GLY:HA2 | 1:C:135:TYR:CE2 | 2.46 | 0.51 |
| 1:A:428:LEU:HD11 | 1:C:139:PRO:HD2 | 1.91 | 0.51 |
| 1:F:142:TYR:O | 1:F:143:GLU:HB2 | 2.11 | 0.51 |
| 2:H:496:TRP:CZ3 | 2:H:506:THR:HB | 2.46 | 0.51 |
| 2:K:496:TRP:CZ3 | 2:K:506:THR:HB | 2.46 | 0.51 |
| 1:A:137:HIS:CD2 | 2:L:602:LEU:HD21 | 2.46 | 0.51 |
| 1:F:335:GLU:OE2 | 1:F:336:GLN:HG2 | 2.11 | 0.51 |
| 1:A:372:ALA:HB1 | 1:B:576:ALA:HB2 | 1.91 | 0.51 |
| 1:C:176:GLU:CG | 1:C:198:LYS:HE3 | 2.41 | 0.51 |
| 1:E:142:TYR:O | 1:E:143:GLU:HB2 | 2.11 | 0.51 |
| 1:E:304:GLN:NE2 | 1:E:345:GLU:OE2 | 2.44 | 0.51 |
| 1:F:304:GLN:NE2 | 1:F:345:GLU:OE2 | 2.44 | 0.51 |
| 1:A:142:TYR:O | 1:A:143:GLU:HB2 | 2.11 | 0.51 |
| 1:B:179:ARG:HB3 | 1:B:179:ARG:CZ | 2.41 | 0.51 |
| 1:C:152:ILE:HG23 | 1:C:496:ARG:HG2 | 1.93 | 0.51 |
| 1:E:335:GLU:OE2 | 1:E:336:GLN:HG2 | 2.11 | 0.51 |
| 1:E:460:PRO:HD3 | 1:E:686:ARG:NH1 | 2.26 | 0.51 |
| 1:E:473:ASP:O | 1:E:480:LYS:HE3 | 2.11 | 0.51 |
| 1:F:473:ASP:O | 1:F:480:LYS:HE3 | 2.11 | 0.51 |
| 2:J:512:PHE:O | 2:J:519:VAL:N | 2.37 | 0.51 |
| 1:B:152:ILE:HG23 | 1:B:496:ARG:HG2 | 1.93 | 0.50 |
| 1:B:176:GLU:CG | 1:B:198:LYS:HE3 | 2.41 | 0.50 |
| 1:D:135:TYR:CE2 | 1:F:220:GLY:HA2 | 2.46 | 0.50 |
| 1:D:142:TYR:O | 1:D:143:GLU:HB2 | 2.11 | 0.50 |
| 1:E:226:ASP:HA | 1:E:229:GLU:OE2 | 2.10 | 0.50 |
| 1:E:372:ALA:HB1 | 1:F:576:ALA:HB2 | 1.92 | 0.50 |
| 1:E:377:ASN:ND2 | 1:E:420:GLY:HA2 | 2.26 | 0.50 |
| 1:E:491:ILE:HG23 | 1:E:686:ARG:HB3 | 1.94 | 0.50 |

Continued on next page...

Not For Manuscript Review

Continued from previous page...

| Atom-1 | Atom-2 | Interatomic distance (Å) | Clash overlap (Å) |
|---|---|---|---|
| 1:E:560:PHE:O | 3:G:802:BTI:H72 | 2.11 | 0.50 |
| 1:F:460:PRO:HD3 | 1:F:686:ARG:NH1 | 2.26 | 0.50 |
| 1:F:491:ILE:HG23 | 1:F:686:ARG:HB3 | 1.94 | 0.50 |
| 1:D:335:GLU:OE2 | 1:D:336:GLN:HG2 | 2.11 | 0.50 |
| 1:E:137:HIS:CD2 | 2:J:602:LEU:HD21 | 2.46 | 0.50 |
| 1:F:226:ASP:HA | 1:F:229:GLU:OE2 | 2.10 | 0.50 |
| 2:I:261:GLU:HG3 | 2:I:262:THR:N | 2.25 | 0.50 |
| 1:A:304:GLN:NE2 | 1:A:345:GLU:OE2 | 2.44 | 0.50 |
| 1:A:460:PRO:HD3 | 1:A:686:ARG:NH1 | 2.26 | 0.50 |
| 1:B:288:LYS:H | 1:B:288:LYS:HD2 | 1.75 | 0.50 |
| 1:C:179:ARG:HB3 | 1:C:179:ARG:CZ | 2.42 | 0.50 |
| 1:C:288:LYS:H | 1:C:288:LYS:HD2 | 1.75 | 0.50 |
| 1:C:473:ASP:O | 1:C:480:LYS:HE3 | 2.11 | 0.50 |
| 1:D:377:ASN:ND2 | 1:D:420:GLY:HA2 | 2.26 | 0.50 |
| 1:D:460:PRO:HD3 | 1:D:686:ARG:NH1 | 2.26 | 0.50 |
| 2:M:496:TRP:CZ3 | 2:M:506:THR:HB | 2.46 | 0.50 |
| 1:A:429:HIS:NE2 | 2:H:607:GLY:CA | 2.73 | 0.50 |
| 1:B:423:TYR:HE2 | 1:F:504:TYR:CE2 | 2.30 | 0.50 |
| 1:B:473:ASP:O | 1:B:480:LYS:HE3 | 2.11 | 0.50 |
| 1:C:460:PRO:HD3 | 1:C:686:ARG:NH1 | 2.26 | 0.50 |
| 1:D:176:GLU:CG | 1:D:198:LYS:HE3 | 2.41 | 0.50 |
| 1:D:304:GLN:NE2 | 1:D:345:GLU:OE2 | 2.44 | 0.50 |
| 1:E:179:ARG:HB3 | 1:E:179:ARG:CZ | 2.41 | 0.50 |
| 1:F:377:ASN:ND2 | 1:F:420:GLY:HA2 | 2.26 | 0.50 |
| 2:I:512:PHE:O | 2:I:519:VAL:N | 2.37 | 0.50 |
| 2:L:236:PHE:CE1 | 2:L:347:LEU:HB3 | 2.47 | 0.50 |
| 2:L:496:TRP:CZ3 | 2:L:506:THR:HB | 2.46 | 0.50 |
| 1:A:335:GLU:OE2 | 1:A:336:GLN:HG2 | 2.11 | 0.50 |
| 2:H:129:GLU:O | 2:H:133:ILE:HG12 | 2.12 | 0.50 |
| 2:I:544:PHE:CB | 2:I:569:PHE:HA | 2.42 | 0.50 |
| 2:J:544:PHE:CB | 2:J:569:PHE:HA | 2.42 | 0.50 |
| 2:K:129:GLU:O | 2:K:133:ILE:HG12 | 2.12 | 0.50 |
| 2:K:568:LEU:HD12 | 2:K:573:MET:O | 2.11 | 0.50 |
| 2:M:236:PHE:CE1 | 2:M:347:LEU:HB3 | 2.47 | 0.50 |
| 1:A:176:GLU:CG | 1:A:198:LYS:HE3 | 2.41 | 0.50 |
| 1:A:377:ASN:ND2 | 1:A:420:GLY:HA2 | 2.26 | 0.50 |
| 1:A:560:PHE:O | 3:G:806:BTI:H72 | 2.11 | 0.50 |
| 1:B:304:GLN:NE2 | 1:B:345:GLU:OE2 | 2.44 | 0.50 |
| 1:B:591:THR:HG22 | 1:B:615:LEU:HG | 1.93 | 0.50 |
| 1:C:137:HIS:CD2 | 2:H:602:LEU:HD21 | 2.47 | 0.50 |
| 1:C:304:GLN:NE2 | 1:C:345:GLU:OE2 | 2.44 | 0.50 |

Continued on next page...

Not For Manuscript Review

Continued from previous page...

| Atom-1 | Atom-2 | Interatomic distance (Å) | Clash overlap (Å) |
|---|---|---|---|
| 1:E:176:GLU:CG | 1:E:198:LYS:HE3 | 2.41 | 0.50 |
| 1:E:191:VAL:HG12 | 1:E:193:GLU:HG2 | 1.93 | 0.50 |
| 1:F:178:LEU:O | 1:F:182:VAL:HG23 | 2.10 | 0.50 |
| 1:F:179:ARG:HB3 | 1:F:179:ARG:CZ | 2.42 | 0.50 |
| 2:H:253:ALA:HB3 | 2:H:254:PRO:HD3 | 1.94 | 0.50 |
| 2:H:568:LEU:HD12 | 2:H:573:MET:O | 2.10 | 0.50 |
| 2:J:261:GLU:HG3 | 2:J:262:THR:N | 2.25 | 0.50 |
| 2:K:490:LEU:HD12 | 2:K:497:ARG:HE | 1.77 | 0.50 |
| 1:A:473:ASP:O | 1:A:480:LYS:HE3 | 2.11 | 0.50 |
| 1:B:460:PRO:HD3 | 1:B:686:ARG:NH1 | 2.26 | 0.50 |
| 1:C:591:THR:HG22 | 1:C:615:LEU:HG | 1.93 | 0.50 |
| 1:D:179:ARG:HB3 | 1:D:179:ARG:CZ | 2.41 | 0.50 |
| 1:E:178:LEU:O | 1:E:182:VAL:HG23 | 2.10 | 0.50 |
| 1:F:176:GLU:CG | 1:F:198:LYS:HE3 | 2.41 | 0.50 |
| 1:F:591:THR:HG22 | 1:F:615:LEU:HG | 1.93 | 0.50 |
| 2:H:490:LEU:HD12 | 2:H:497:ARG:HE | 1.77 | 0.50 |
| 2:J:129:GLU:O | 2:J:133:ILE:HG12 | 2.12 | 0.50 |
| 2:J:496:TRP:CZ3 | 2:J:506:THR:HB | 2.46 | 0.50 |
| 2:M:544:PHE:CB | 2:M:569:PHE:HA | 2.42 | 0.50 |
| 1:C:216:HIS:ND1 | 1:C:223:LEU:HB2 | 2.27 | 0.50 |
| 1:D:180:ALA:O | 1:D:183:ARG:N | 2.45 | 0.50 |
| 1:D:473:ASP:O | 1:D:480:LYS:HE3 | 2.11 | 0.50 |
| 1:F:191:VAL:HG12 | 1:F:193:GLU:HG2 | 1.93 | 0.50 |
| 2:I:96:LEU:HD22 | 2:I:99:ASN:HD22 | 1.77 | 0.50 |
| 2:I:383:ARG:CD | 2:K:34:GLU:OE2 | 2.51 | 0.50 |
| 2:I:496:TRP:CZ3 | 2:I:506:THR:HB | 2.46 | 0.50 |
| 2:J:96:LEU:HD22 | 2:J:99:ASN:HD22 | 1.77 | 0.50 |
| 2:L:650:MET:HE2 | 2:L:651:LYS:NZ | 2.18 | 0.50 |
| 1:A:179:ARG:HB3 | 1:A:179:ARG:CZ | 2.41 | 0.50 |
| 1:A:180:ALA:O | 1:A:183:ARG:N | 2.45 | 0.50 |
| 1:B:216:HIS:ND1 | 1:B:223:LEU:HB2 | 2.27 | 0.50 |
| 1:D:216:HIS:ND1 | 1:D:223:LEU:HB2 | 2.27 | 0.50 |
| 2:K:236:PHE:CE1 | 2:K:347:LEU:HB3 | 2.47 | 0.50 |
| 2:K:253:ALA:HB3 | 2:K:254:PRO:HD3 | 1.94 | 0.50 |
| 2:L:124:MET:HE1 | 2:L:301:MET:HB3 | 1.94 | 0.50 |
| 2:L:544:PHE:CB | 2:L:569:PHE:HA | 2.42 | 0.50 |
| 2:M:124:MET:HE1 | 2:M:301:MET:HB3 | 1.94 | 0.50 |
| 2:M:650:MET:HE2 | 2:M:651:LYS:NZ | 2.18 | 0.50 |
| 1:C:142:TYR:O | 1:C:143:GLU:HB2 | 2.11 | 0.49 |
| 1:E:137:HIS:HA | 2:J:602:LEU:HD11 | 1.94 | 0.49 |
| 1:E:591:THR:HG22 | 1:E:615:LEU:HG | 1.94 | 0.49 |

Continued on next page...

Not For Manuscript Review

Continued from previous page...

| Atom-1 | Atom-2 | Interatomic distance (Å) | Clash overlap (Å) |
|---|---|---|---|
| 2:H:236:PHE:CE1 | 2:H:347:LEU:HB3 | 2.47 | 0.49 |
| 2:I:129:GLU:O | 2:I:133:ILE:HG12 | 2.12 | 0.49 |
| 2:I:490:LEU:HD12 | 2:I:497:ARG:HE | 1.77 | 0.49 |
| 2:J:490:LEU:HD12 | 2:J:497:ARG:HE | 1.77 | 0.49 |
| 2:J:650:MET:HE1 | 2:J:651:LYS:NZ | 2.12 | 0.49 |
| 2:K:633:LYS:NZ | 2:K:634:SER:OG | 2.45 | 0.49 |
| 1:A:216:HIS:ND1 | 1:A:223:LEU:HB2 | 2.27 | 0.49 |
| 1:B:142:TYR:O | 1:B:143:GLU:HB2 | 2.11 | 0.49 |
| 1:C:491:ILE:HG23 | 1:C:686:ARG:HB3 | 1.94 | 0.49 |
| 1:D:152:ILE:HG23 | 1:D:496:ARG:HG2 | 1.93 | 0.49 |
| 2:H:633:LYS:NZ | 2:H:634:SER:OG | 2.45 | 0.49 |
| 2:J:573:MET:HG2 | 2:J:574:HIS:H | 1.77 | 0.49 |
| 2:K:573:MET:HG2 | 2:K:574:HIS:H | 1.77 | 0.49 |
| 2:L:490:LEU:HD12 | 2:L:497:ARG:HE | 1.77 | 0.49 |
| 1:B:491:ILE:HG23 | 1:B:686:ARG:HB3 | 1.94 | 0.49 |
| 1:C:372:ALA:HB1 | 1:D:576:ALA:HB2 | 1.93 | 0.49 |
| 1:F:397:ALA:CB | 2:J:650:MET:HG2 | 2.42 | 0.49 |
| 2:H:573:MET:HG2 | 2:H:574:HIS:H | 1.77 | 0.49 |
| 2:I:573:MET:HG2 | 2:I:574:HIS:H | 1.77 | 0.49 |
| 2:J:425:LEU:O | 2:J:426:ARG:C | 2.51 | 0.49 |
| 2:M:490:LEU:HD12 | 2:M:497:ARG:HE | 1.77 | 0.49 |
| 1:A:152:ILE:HG23 | 1:A:496:ARG:HG2 | 1.94 | 0.49 |
| 1:C:625:VAL:HG12 | 1:D:328:GLN:HG3 | 1.94 | 0.49 |
| 1:D:167:ASN:HB3 | 1:D:471:ILE:O | 2.13 | 0.49 |
| 1:D:397:ALA:CB | 2:H:650:MET:HG2 | 2.42 | 0.49 |
| 1:E:156:PRO:HG2 | 1:E:490:ARG:NH1 | 2.28 | 0.49 |
| 2:I:238:ARG:HG2 | 2:I:253:ALA:HA | 1.94 | 0.49 |
| 2:I:425:LEU:O | 2:I:426:ARG:C | 2.51 | 0.49 |
| 1:A:167:ASN:HB3 | 1:A:471:ILE:O | 2.13 | 0.49 |
| 1:B:335:GLU:OE2 | 1:B:336:GLN:HG2 | 2.11 | 0.49 |
| 1:C:137:HIS:HA | 2:H:602:LEU:HD11 | 1.92 | 0.49 |
| 1:C:180:ALA:O | 1:C:183:ARG:N | 2.45 | 0.49 |
| 1:C:335:GLU:OE2 | 1:C:336:GLN:HG2 | 2.11 | 0.49 |
| 1:C:576:ALA:HB2 | 1:D:372:ALA:HB1 | 1.93 | 0.49 |
| 1:D:491:ILE:HG23 | 1:D:686:ARG:HB3 | 1.94 | 0.49 |
| 1:E:328:GLN:HG3 | 1:F:625:VAL:HG12 | 1.95 | 0.49 |
| 2:J:236:PHE:CE1 | 2:J:347:LEU:HB3 | 2.47 | 0.49 |
| 2:J:238:ARG:HG2 | 2:J:253:ALA:HA | 1.94 | 0.49 |
| 1:A:491:ILE:HG23 | 1:A:686:ARG:HB3 | 1.94 | 0.49 |
| 1:E:625:VAL:HG12 | 1:F:328:GLN:HG3 | 1.95 | 0.49 |
| 1:F:156:PRO:HG2 | 1:F:490:ARG:NH1 | 2.28 | 0.49 |

Continued on next page...

Not For Manuscript Review

Continued from previous page...

| Atom-1 | Atom-2 | Interatomic distance (Å) | Clash overlap (Å) |
|---|---|---|---|
| 1:F:167:ASN:HB3 | 1:F:471:ILE:O | 2.13 | 0.49 |
| 2:H:257:HIS:NE2 | 2:H:455:ARG:O | 2.43 | 0.49 |
| 2:I:236:PHE:CE1 | 2:I:347:LEU:HB3 | 2.47 | 0.49 |
| 1:A:156:PRO:HG2 | 1:A:490:ARG:NH1 | 2.28 | 0.49 |
| 1:B:346:ALA:O | 1:B:350:ILE:HG12 | 2.13 | 0.49 |
| 1:B:429:HIS:NE2 | 2:I:607:GLY:HA2 | 2.28 | 0.49 |
| 1:C:156:PRO:HG2 | 1:C:490:ARG:NH1 | 2.28 | 0.49 |
| 1:C:346:ALA:O | 1:C:350:ILE:HG12 | 2.13 | 0.49 |
| 1:F:235:GLY:HA3 | 2:M:494:ASP:OD1 | 2.13 | 0.49 |
| 1:F:529:LEU:N | 1:F:559:GLY:O | 2.42 | 0.49 |
| 2:L:238:ARG:HG2 | 2:L:253:ALA:HA | 1.94 | 0.49 |
| 2:M:238:ARG:HG2 | 2:M:253:ALA:HA | 1.94 | 0.49 |
| 1:B:156:PRO:HG2 | 1:B:490:ARG:NH1 | 2.28 | 0.49 |
| 1:B:180:ALA:O | 1:B:183:ARG:N | 2.45 | 0.49 |
| 1:D:156:PRO:HG2 | 1:D:490:ARG:NH1 | 2.28 | 0.49 |
| 1:E:167:ASN:HB3 | 1:E:471:ILE:O | 2.13 | 0.49 |
| 2:I:650:MET:HE1 | 2:I:651:LYS:NZ | 2.14 | 0.49 |
| 2:K:257:HIS:NE2 | 2:K:455:ARG:O | 2.43 | 0.49 |
| 2:L:96:LEU:HD22 | 2:L:99:ASN:HD22 | 1.77 | 0.49 |
| 1:B:415:ARG:HD2 | 1:B:417:SER:HB2 | 1.95 | 0.49 |
| 1:C:415:ARG:HD2 | 1:C:417:SER:HB2 | 1.95 | 0.49 |
| 2:H:96:LEU:HD22 | 2:H:99:ASN:HD22 | 1.77 | 0.49 |
| 2:H:155:LEU:O | 2:H:159:ALA:HB2 | 2.13 | 0.49 |
| 2:H:544:PHE:CB | 2:H:569:PHE:HA | 2.42 | 0.49 |
| 2:J:34:GLU:OE2 | 2:L:383:ARG:CD | 2.49 | 0.49 |
| 2:J:529:ASN:CG | 2:J:532:LYS:HB3 | 2.33 | 0.49 |
| 2:K:155:LEU:O | 2:K:159:ALA:HB2 | 2.13 | 0.49 |
| 2:L:155:LEU:O | 2:L:159:ALA:HB2 | 2.13 | 0.49 |
| 2:L:425:LEU:O | 2:L:426:ARG:C | 2.51 | 0.49 |
| 2:M:155:LEU:O | 2:M:159:ALA:HB2 | 2.13 | 0.49 |
| 1:A:137:HIS:HA | 2:L:602:LEU:HD11 | 1.94 | 0.49 |
| 1:A:591:THR:HG22 | 1:A:615:LEU:HG | 1.93 | 0.49 |
| 2:J:619:ILE:HD11 | 2:J:681:ALA:HB3 | 1.95 | 0.49 |
| 2:K:356:VAL:HG23 | 2:K:408:LEU:HB2 | 1.95 | 0.49 |
| 2:K:544:PHE:CB | 2:K:569:PHE:HA | 2.42 | 0.49 |
| 2:M:129:GLU:O | 2:M:133:ILE:HG12 | 2.12 | 0.49 |
| 1:C:411:ASP:O | 1:E:504:TYR:OH | 2.31 | 0.48 |
| 1:D:591:THR:HG22 | 1:D:615:LEU:HG | 1.93 | 0.48 |
| 2:H:356:VAL:HG23 | 2:H:408:LEU:HB2 | 1.95 | 0.48 |
| 2:H:565:PHE:O | 2:H:576:THR:HA | 2.13 | 0.48 |
| 2:I:529:ASN:CG | 2:I:532:LYS:HB3 | 2.33 | 0.48 |

Continued on next page...

Not For Manuscript Review

Continued from previous page...

| Atom-1 | Atom-2 | Interatomic distance (Å) | Clash overlap (Å) |
|---|---|---|---|
| 2:J:552:THR:OG1 | 2:J:563:ASN:OD1 | 2.29 | 0.48 |
| 2:K:96:LEU:HD22 | 2:K:99:ASN:HD22 | 1.77 | 0.48 |
| 2:K:565:PHE:O | 2:K:576:THR:HA | 2.13 | 0.48 |
| 2:M:96:LEU:HD22 | 2:M:99:ASN:HD22 | 1.77 | 0.48 |
| 1:A:346:ALA:O | 1:A:350:ILE:HG12 | 2.13 | 0.48 |
| 1:D:346:ALA:O | 1:D:350:ILE:HG12 | 2.13 | 0.48 |
| 1:E:429:HIS:NE2 | 2:L:607:GLY:CA | 2.76 | 0.48 |
| 1:F:152:ILE:HG23 | 1:F:496:ARG:HG2 | 1.93 | 0.48 |
| 2:H:169:LYS:NZ | 2:H:173:GLY:O | 2.37 | 0.48 |
| 2:I:378:PRO:HD3 | 2:I:432:TYR:CD1 | 2.47 | 0.48 |
| 2:I:619:ILE:HD11 | 2:I:681:ALA:HB3 | 1.96 | 0.48 |
| 2:J:378:PRO:HD3 | 2:J:432:TYR:CD1 | 2.47 | 0.48 |
| 2:L:129:GLU:O | 2:L:133:ILE:HG12 | 2.12 | 0.48 |
| 2:L:356:VAL:HG23 | 2:L:408:LEU:HB2 | 1.95 | 0.48 |
| 2:M:565:PHE:O | 2:M:576:THR:HA | 2.13 | 0.48 |
| 2:M:633:LYS:NZ | 2:M:634:SER:OG | 2.45 | 0.48 |
| 1:B:411:ASP:O | 1:F:504:TYR:OH | 2.31 | 0.48 |
| 1:C:423:TYR:HE2 | 1:E:504:TYR:CE2 | 2.32 | 0.48 |
| 1:E:152:ILE:HG23 | 1:E:496:ARG:HG2 | 1.93 | 0.48 |
| 2:H:529:ASN:CG | 2:H:532:LYS:HB3 | 2.33 | 0.48 |
| 2:K:169:LYS:NZ | 2:K:173:GLY:O | 2.37 | 0.48 |
| 2:K:529:ASN:CG | 2:K:532:LYS:HB3 | 2.33 | 0.48 |
| 2:L:565:PHE:O | 2:L:576:THR:HA | 2.14 | 0.48 |
| 2:L:573:MET:HG2 | 2:L:574:HIS:H | 1.77 | 0.48 |
| 2:L:633:LYS:NZ | 2:L:634:SER:OG | 2.45 | 0.48 |
| 2:M:356:VAL:HG23 | 2:M:408:LEU:HB2 | 1.95 | 0.48 |
| 2:M:425:LEU:O | 2:M:426:ARG:C | 2.51 | 0.48 |
| 2:M:573:MET:HG2 | 2:M:574:HIS:H | 1.77 | 0.48 |
| 1:B:181:ARG:NH1 | 1:B:247:TRP:HE1 | 2.11 | 0.48 |
| 1:E:346:ALA:O | 1:E:350:ILE:HG12 | 2.13 | 0.48 |
| 2:H:378:PRO:HD3 | 2:H:432:TYR:CD1 | 2.47 | 0.48 |
| 2:H:652:MET:HG3 | 2:H:654:HIS:CD2 | 2.49 | 0.48 |
| 2:I:652:MET:HG3 | 2:I:654:HIS:CD2 | 2.49 | 0.48 |
| 2:J:652:MET:HG3 | 2:J:654:HIS:CD2 | 2.49 | 0.48 |
| 2:K:378:PRO:HD3 | 2:K:432:TYR:CD1 | 2.47 | 0.48 |
| 2:K:652:MET:HG3 | 2:K:654:HIS:CD2 | 2.49 | 0.48 |
| 2:M:253:ALA:HB3 | 2:M:254:PRO:HD3 | 1.94 | 0.48 |
| 1:B:504:TYR:OH | 1:D:411:ASP:O | 2.30 | 0.48 |
| 1:C:181:ARG:NH1 | 1:C:247:TRP:HE1 | 2.11 | 0.48 |
| 1:D:144:ARG:CG | 2:M:606:PHE:CE2 | 2.96 | 0.48 |
| 1:E:182:VAL:HG21 | 2:L:503:ASN:HB2 | 1.96 | 0.48 |

Continued on next page...

Not For Manuscript Review

Continued from previous page...

| Atom-1 | Atom-2 | Interatomic distance (Å) | Clash overlap (Å) |
|---|---|---|---|
| 1:F:242:SER:HB3 | 1:F:299:LYS:HE3 | 1.95 | 0.48 |
| 2:I:552:THR:OG1 | 2:I:563:ASN:OD1 | 2.30 | 0.48 |
| 2:I:633:LYS:NZ | 2:I:634:SER:OG | 2.45 | 0.48 |
| 2:J:253:ALA:HB3 | 2:J:254:PRO:HD3 | 1.94 | 0.48 |
| 2:J:633:LYS:NZ | 2:J:634:SER:OG | 2.45 | 0.48 |
| 2:L:253:ALA:HB3 | 2:L:254:PRO:HD3 | 1.94 | 0.48 |
| 2:L:376:ARG:HA | 2:L:376:ARG:HH11 | 1.79 | 0.48 |
| 2:M:652:MET:O | 2:M:654:HIS:CD2 | 2.67 | 0.48 |
| 1:A:328:GLN:CG | 1:B:625:VAL:HG12 | 2.43 | 0.48 |
| 1:A:625:VAL:HG12 | 1:B:328:GLN:HG3 | 1.94 | 0.48 |
| 1:C:182:VAL:HG21 | 2:J:503:ASN:HB2 | 1.96 | 0.48 |
| 1:D:181:ARG:NH1 | 1:D:247:TRP:HE1 | 2.11 | 0.48 |
| 1:E:242:SER:HB3 | 1:E:299:LYS:HE3 | 1.95 | 0.48 |
| 1:F:346:ALA:O | 1:F:350:ILE:HG12 | 2.13 | 0.48 |
| 2:I:253:ALA:HB3 | 2:I:254:PRO:HD3 | 1.94 | 0.48 |
| 2:I:565:PHE:O | 2:I:576:THR:HA | 2.14 | 0.48 |
| 2:J:257:HIS:NE2 | 2:J:455:ARG:O | 2.43 | 0.48 |
| 2:J:565:PHE:O | 2:J:576:THR:HA | 2.13 | 0.48 |
| 2:L:529:ASN:CG | 2:L:532:LYS:HB3 | 2.33 | 0.48 |
| 2:L:652:MET:O | 2:L:654:HIS:CD2 | 2.67 | 0.48 |
| 2:M:17:LEU:HD23 | 2:M:89:ILE:HG12 | 1.96 | 0.48 |
| 2:M:376:ARG:HA | 2:M:376:ARG:HH11 | 1.79 | 0.48 |
| 1:A:181:ARG:NH1 | 1:A:247:TRP:HE1 | 2.11 | 0.48 |
| 1:A:423:TYR:HE2 | 1:C:504:TYR:CE2 | 2.32 | 0.48 |
| 1:D:429:HIS:NE2 | 2:K:607:GLY:HA2 | 2.27 | 0.48 |
| 1:E:216:HIS:ND1 | 1:E:223:LEU:HB2 | 2.27 | 0.48 |
| 1:F:216:HIS:ND1 | 1:F:223:LEU:HB2 | 2.27 | 0.48 |
| 2:I:155:LEU:O | 2:I:159:ALA:HB2 | 2.13 | 0.48 |
| 2:I:257:HIS:NE2 | 2:I:455:ARG:O | 2.43 | 0.48 |
| 2:J:155:LEU:O | 2:J:159:ALA:HB2 | 2.13 | 0.48 |
| 2:L:652:MET:HG3 | 2:L:654:HIS:CD2 | 2.49 | 0.48 |
| 2:M:30:ARG:HA | 2:M:33:ARG:NH2 | 2.29 | 0.48 |
| 1:C:251:TRP:HE1 | 1:C:255:LYS:HA | 1.79 | 0.48 |
| 2:H:553:SER:HB3 | 2:H:561:ILE:HG13 | 1.96 | 0.48 |
| 2:I:356:VAL:HG23 | 2:I:408:LEU:HB2 | 1.95 | 0.48 |
| 2:L:17:LEU:HD23 | 2:L:89:ILE:HG12 | 1.96 | 0.48 |
| 2:M:529:ASN:CG | 2:M:532:LYS:HB3 | 2.33 | 0.48 |
| 2:M:553:SER:HB3 | 2:M:561:ILE:HG13 | 1.96 | 0.48 |
| 2:M:652:MET:HG3 | 2:M:654:HIS:CD2 | 2.49 | 0.48 |
| 1:A:251:TRP:HE1 | 1:A:255:LYS:HA | 1.79 | 0.48 |
| 1:B:242:SER:HB3 | 1:B:299:LYS:HE3 | 1.95 | 0.48 |

Continued on next page...

Not For Manuscript Review

Continued from previous page...

| Atom-1 | Atom-2 | Interatomic distance (Å) | Clash overlap (Å) |
|---|---|---|---|
| 1:B:251:TRP:HE1 | 1:B:255:LYS:HA | 1.79 | 0.48 |
| 1:C:242:SER:HB3 | 1:C:299:LYS:HE3 | 1.95 | 0.48 |
| 1:C:328:GLN:HG3 | 1:D:625:VAL:HG12 | 1.95 | 0.48 |
| 1:C:429:HIS:NE2 | 2:J:607:GLY:CA | 2.76 | 0.48 |
| 1:C:493:ASP:OD1 | 1:C:686:ARG:NH2 | 2.47 | 0.48 |
| 1:E:476:SER:O | 1:E:477:ASP:C | 2.52 | 0.48 |
| 2:I:652:MET:O | 2:I:654:HIS:CD2 | 2.67 | 0.48 |
| 2:J:165:PRO:HB2 | 2:J:212:ARG:HA | 1.96 | 0.48 |
| 2:J:652:MET:O | 2:J:654:HIS:CD2 | 2.67 | 0.48 |
| 2:K:553:SER:HB3 | 2:K:561:ILE:HG13 | 1.96 | 0.48 |
| 2:L:553:SER:HB3 | 2:L:561:ILE:HG13 | 1.96 | 0.48 |
| 1:A:332:PHE:HB3 | 1:A:333:PRO:HD3 | 1.96 | 0.48 |
| 1:A:415:ARG:HD2 | 1:A:417:SER:HB2 | 1.95 | 0.48 |
| 1:A:504:TYR:CE1 | 1:E:415:ARG:N | 2.82 | 0.48 |
| 1:B:167:ASN:HB3 | 1:B:471:ILE:O | 2.13 | 0.48 |
| 1:B:493:ASP:OD1 | 1:B:686:ARG:NH2 | 2.47 | 0.48 |
| 1:C:177:GLN:CG | 1:C:198:LYS:HZ2 | 2.27 | 0.48 |
| 1:D:251:TRP:HE1 | 1:D:255:LYS:HA | 1.79 | 0.48 |
| 1:D:332:PHE:HB3 | 1:D:333:PRO:HD3 | 1.96 | 0.48 |
| 1:D:415:ARG:HD2 | 1:D:417:SER:HB2 | 1.95 | 0.48 |
| 1:E:251:TRP:HE1 | 1:E:255:LYS:HA | 1.79 | 0.48 |
| 1:F:251:TRP:HE1 | 1:F:255:LYS:HA | 1.79 | 0.48 |
| 1:F:476:SER:O | 1:F:477:ASP:C | 2.52 | 0.48 |
| 2:H:376:ARG:HA | 2:H:376:ARG:HH11 | 1.79 | 0.48 |
| 2:I:165:PRO:HB2 | 2:I:212:ARG:HA | 1.96 | 0.48 |
| 2:J:356:VAL:HG23 | 2:J:408:LEU:HB2 | 1.95 | 0.48 |
| 2:L:30:ARG:HA | 2:L:33:ARG:NH2 | 2.29 | 0.48 |
| 2:M:165:PRO:HB2 | 2:M:212:ARG:HA | 1.96 | 0.48 |
| 1:A:476:SER:O | 1:A:477:ASP:C | 2.52 | 0.47 |
| 1:C:167:ASN:HB3 | 1:C:471:ILE:O | 2.13 | 0.47 |
| 1:D:476:SER:O | 1:D:477:ASP:C | 2.52 | 0.47 |
| 1:F:385:THR:HA | 1:F:409:GLY:HA2 | 1.96 | 0.47 |
| 2:H:238:ARG:HG2 | 2:H:253:ALA:HA | 1.94 | 0.47 |
| 2:H:652:MET:O | 2:H:654:HIS:CD2 | 2.67 | 0.47 |
| 2:I:30:ARG:HA | 2:I:33:ARG:NH2 | 2.29 | 0.47 |
| 2:J:30:ARG:HA | 2:J:33:ARG:NH2 | 2.29 | 0.47 |
| 2:J:604:ASP:O | 2:J:605:GLY:C | 2.53 | 0.47 |
| 2:K:376:ARG:HA | 2:K:376:ARG:HH11 | 1.79 | 0.47 |
| 2:K:652:MET:O | 2:K:654:HIS:CD2 | 2.67 | 0.47 |
| 1:A:140:ILE:O | 1:A:141:ASP:HB2 | 2.15 | 0.47 |
| 1:A:328:GLN:NE2 | 1:B:625:VAL:HA | 2.30 | 0.47 |

Continued on next page...

Not For Manuscript Review

*Continued from previous page...*

| Atom-1 | Atom-2 | Interatomic distance (Å) | Clash overlap (Å) |
|---|---|---|---|
| 1:B:476:SER:O | 1:B:477:ASP:C | 2.52 | 0.47 |
| 1:C:476:SER:O | 1:C:477:ASP:C | 2.52 | 0.47 |
| 1:D:478:VAL:HG23 | 2:M:651:LYS:HA | 1.96 | 0.47 |
| 1:E:385:THR:HA | 1:E:409:GLY:HA2 | 1.96 | 0.47 |
| 2:H:17:LEU:HD23 | 2:H:89:ILE:HG12 | 1.96 | 0.47 |
| 2:H:165:PRO:HB2 | 2:H:212:ARG:HA | 1.96 | 0.47 |
| 2:H:425:LEU:O | 2:H:426:ARG:C | 2.51 | 0.47 |
| 2:H:604:ASP:O | 2:H:605:GLY:C | 2.53 | 0.47 |
| 2:I:34:GLU:OE2 | 2:M:383:ARG:CD | 2.51 | 0.47 |
| 2:I:604:ASP:O | 2:I:605:GLY:C | 2.53 | 0.47 |
| 2:K:238:ARG:HG2 | 2:K:253:ALA:HA | 1.95 | 0.47 |
| 2:K:604:ASP:O | 2:K:605:GLY:C | 2.53 | 0.47 |
| 2:L:165:PRO:HB2 | 2:L:212:ARG:HA | 1.96 | 0.47 |
| 2:L:604:ASP:O | 2:L:605:GLY:C | 2.53 | 0.47 |
| 2:M:604:ASP:O | 2:M:605:GLY:C | 2.53 | 0.47 |
| 1:D:140:ILE:O | 1:D:141:ASP:HB2 | 2.15 | 0.47 |
| 1:E:140:ILE:O | 1:E:141:ASP:HB2 | 2.15 | 0.47 |
| 1:F:140:ILE:O | 1:F:141:ASP:HB2 | 2.15 | 0.47 |
| 1:F:450:ARG:HD2 | 1:F:450:ARG:HA | 1.68 | 0.47 |
| 2:I:553:SER:HB3 | 2:I:561:ILE:HG13 | 1.96 | 0.47 |
| 2:K:17:LEU:HD23 | 2:K:89:ILE:HG12 | 1.96 | 0.47 |
| 2:K:165:PRO:HB2 | 2:K:212:ARG:HA | 1.96 | 0.47 |
| 2:L:182:VAL:HG13 | 2:L:182:VAL:O | 2.15 | 0.47 |
| 2:M:182:VAL:O | 2:M:182:VAL:HG13 | 2.15 | 0.47 |
| 2:M:257:HIS:NE2 | 2:M:455:ARG:O | 2.43 | 0.47 |
| 1:C:332:PHE:HB3 | 1:C:333:PRO:HD3 | 1.96 | 0.47 |
| 1:D:242:SER:HB3 | 1:D:299:LYS:HE3 | 1.95 | 0.47 |
| 2:L:512:PHE:O | 2:L:519:VAL:N | 2.37 | 0.47 |
| 2:M:233:VAL:HG21 | 2:M:342:GLN:HA | 1.96 | 0.47 |
| 1:A:137:HIS:HD2 | 2:L:602:LEU:HD21 | 1.79 | 0.47 |
| 1:E:181:ARG:NH1 | 1:E:247:TRP:HE1 | 2.11 | 0.47 |
| 2:H:347:LEU:C | 2:H:347:LEU:HD12 | 2.35 | 0.47 |
| 2:I:585:ASP:HA | 2:I:599:LEU:HD23 | 1.97 | 0.47 |
| 2:J:553:SER:HB3 | 2:J:561:ILE:HG13 | 1.96 | 0.47 |
| 2:J:565:PHE:HB2 | 2:J:577:VAL:CG2 | 2.38 | 0.47 |
| 2:J:585:ASP:HA | 2:J:599:LEU:HD23 | 1.97 | 0.47 |
| 2:K:425:LEU:O | 2:K:426:ARG:C | 2.51 | 0.47 |
| 2:L:233:VAL:HG21 | 2:L:342:GLN:HA | 1.96 | 0.47 |
| 2:M:117:PRO:HD3 | 2:M:280:VAL:HG12 | 1.96 | 0.47 |
| 1:A:242:SER:HB3 | 1:A:299:LYS:HE3 | 1.95 | 0.47 |
| 1:B:332:PHE:HB3 | 1:B:333:PRO:HD3 | 1.96 | 0.47 |

*Continued on next page...*

Not For Manuscript Review

*Continued from previous page...*

| Atom-1 | Atom-2 | Interatomic distance (Å) | Clash overlap (Å) |
|---|---|---|---|
| 1:C:415:ARG:N | 1:E:504:TYR:CE1 | 2.83 | 0.47 |
| 1:D:656:LYS:O | 1:D:660:LYS:HG2 | 2.14 | 0.47 |
| 1:F:181:ARG:NH1 | 1:F:247:TRP:HE1 | 2.12 | 0.47 |
| 1:F:243:GLN:HE22 | 2:M:502:LEU:HD13 | 1.79 | 0.47 |
| 2:H:552:THR:OG1 | 2:H:563:ASN:OD1 | 2.29 | 0.47 |
| 2:I:233:VAL:HG21 | 2:I:342:GLN:HA | 1.96 | 0.47 |
| 2:I:565:PHE:HB2 | 2:I:577:VAL:CG2 | 2.38 | 0.47 |
| 2:J:233:VAL:HG21 | 2:J:342:GLN:HA | 1.96 | 0.47 |
| 2:K:233:VAL:HG21 | 2:K:342:GLN:HA | 1.96 | 0.47 |
| 2:K:347:LEU:C | 2:K:347:LEU:HD12 | 2.35 | 0.47 |
| 2:K:585:ASP:HA | 2:K:599:LEU:HD23 | 1.97 | 0.47 |
| 2:K:619:ILE:HD11 | 2:K:681:ALA:HB3 | 1.96 | 0.47 |
| 2:L:117:PRO:HD3 | 2:L:280:VAL:HG12 | 1.96 | 0.47 |
| 2:L:257:HIS:NE2 | 2:L:455:ARG:O | 2.44 | 0.47 |
| 2:L:378:PRO:HD3 | 2:L:432:TYR:CD1 | 2.47 | 0.47 |
| 2:L:619:ILE:HD11 | 2:L:681:ALA:HB3 | 1.95 | 0.47 |
| 2:M:512:PHE:O | 2:M:519:VAL:N | 2.37 | 0.47 |
| 1:B:656:LYS:O | 1:B:660:LYS:HG2 | 2.14 | 0.47 |
| 1:C:328:GLN:CG | 1:D:625:VAL:HG12 | 2.45 | 0.47 |
| 1:C:656:LYS:O | 1:C:660:LYS:HG2 | 2.14 | 0.47 |
| 1:F:415:ARG:HD2 | 1:F:417:SER:HB2 | 1.95 | 0.47 |
| 2:H:585:ASP:HA | 2:H:599:LEU:HD23 | 1.97 | 0.47 |
| 2:J:117:PRO:HD3 | 2:J:280:VAL:HG12 | 1.96 | 0.47 |
| 2:L:585:ASP:HA | 2:L:599:LEU:HD23 | 1.97 | 0.47 |
| 2:M:619:ILE:HD11 | 2:M:681:ALA:HB3 | 1.95 | 0.47 |
| 1:A:625:VAL:HG12 | 1:B:328:GLN:CG | 2.45 | 0.47 |
| 1:E:415:ARG:HD2 | 1:E:417:SER:HB2 | 1.95 | 0.47 |
| 2:H:30:ARG:HA | 2:H:33:ARG:NH2 | 2.29 | 0.47 |
| 2:H:233:VAL:HG21 | 2:H:342:GLN:HA | 1.96 | 0.47 |
| 2:H:619:ILE:HD11 | 2:H:681:ALA:HB3 | 1.96 | 0.47 |
| 2:I:117:PRO:HD3 | 2:I:280:VAL:HG12 | 1.96 | 0.47 |
| 2:J:376:ARG:HA | 2:J:376:ARG:HH11 | 1.79 | 0.47 |
| 2:K:552:THR:OG1 | 2:K:563:ASN:OD1 | 2.29 | 0.47 |
| 2:M:585:ASP:HA | 2:M:599:LEU:HD23 | 1.97 | 0.47 |
| 1:A:656:LYS:O | 1:A:660:LYS:HG2 | 2.14 | 0.47 |
| 1:C:625:VAL:HA | 1:D:328:GLN:NE2 | 2.30 | 0.47 |
| 1:E:184:TYR:CE2 | 1:E:201:VAL:HG21 | 2.50 | 0.47 |
| 1:F:184:TYR:CE2 | 1:F:201:VAL:HG21 | 2.50 | 0.47 |
| 1:F:478:VAL:HG23 | 2:I:651:LYS:HA | 1.97 | 0.47 |
| 2:I:17:LEU:HD23 | 2:I:89:ILE:HG12 | 1.96 | 0.47 |
| 2:M:532:LYS:HE3 | 2:M:534:PHE:HE1 | 1.80 | 0.47 |

*Continued on next page...*

Not For Manuscript Review

Continued from previous page...

| Atom-1 | Atom-2 | Interatomic distance (Å) | Clash overlap (Å) |
|---|---|---|---|
| 1:A:385:THR:HA | 1:A:409:GLY:HA2 | 1.96 | 0.46 |
| 1:B:385:THR:HA | 1:B:409:GLY:HA2 | 1.96 | 0.46 |
| 1:C:653:PHE:O | 1:C:657:VAL:HG22 | 2.15 | 0.46 |
| 1:D:653:PHE:O | 1:D:657:VAL:HG22 | 2.15 | 0.46 |
| 1:E:450:ARG:HD2 | 1:E:450:ARG:HA | 1.68 | 0.46 |
| 1:E:653:PHE:O | 1:E:657:VAL:HG22 | 2.16 | 0.46 |
| 2:H:652:MET:HG3 | 2:H:654:HIS:HB3 | 1.98 | 0.46 |
| 2:I:376:ARG:HA | 2:I:376:ARG:HH11 | 1.79 | 0.46 |
| 2:J:309:HIS:ND1 | 2:J:310:PRO:HD3 | 2.30 | 0.46 |
| 2:K:30:ARG:HA | 2:K:33:ARG:NH2 | 2.29 | 0.46 |
| 2:L:532:LYS:HE3 | 2:L:534:PHE:HE1 | 1.80 | 0.46 |
| 2:M:378:PRO:HD3 | 2:M:432:TYR:CD1 | 2.47 | 0.46 |
| 1:A:182:VAL:HG21 | 2:H:503:ASN:HB2 | 1.97 | 0.46 |
| 1:A:504:TYR:OH | 1:E:411:ASP:O | 2.32 | 0.46 |
| 1:A:653:PHE:O | 1:A:657:VAL:HG22 | 2.16 | 0.46 |
| 1:B:140:ILE:O | 1:B:141:ASP:HB2 | 2.15 | 0.46 |
| 1:B:653:PHE:O | 1:B:657:VAL:HG22 | 2.16 | 0.46 |
| 1:C:140:ILE:O | 1:C:141:ASP:HB2 | 2.15 | 0.46 |
| 1:C:599:GLY:HA3 | 1:C:626:MET:HA | 1.97 | 0.46 |
| 1:D:385:THR:HA | 1:D:409:GLY:HA2 | 1.96 | 0.46 |
| 1:E:180:ALA:O | 1:E:183:ARG:N | 2.45 | 0.46 |
| 2:I:227:ASP:HB3 | 2:I:338:LEU:CD2 | 2.45 | 0.46 |
| 2:J:17:LEU:HD23 | 2:J:89:ILE:HG12 | 1.96 | 0.46 |
| 2:L:309:HIS:ND1 | 2:L:310:PRO:HD3 | 2.30 | 0.46 |
| 2:L:347:LEU:C | 2:L:347:LEU:HD12 | 2.36 | 0.46 |
| 2:L:571:ASN:OD1 | 2:L:572:GLY:N | 2.48 | 0.46 |
| 2:M:571:ASN:OD1 | 2:M:572:GLY:N | 2.48 | 0.46 |
| 1:A:411:ASP:O | 1:C:504:TYR:OH | 2.32 | 0.46 |
| 1:B:184:TYR:CE2 | 1:B:201:VAL:HG21 | 2.50 | 0.46 |
| 1:B:599:GLY:HA3 | 1:B:626:MET:HA | 1.97 | 0.46 |
| 1:C:184:TYR:CE2 | 1:C:201:VAL:HG21 | 2.50 | 0.46 |
| 1:C:625:VAL:HG12 | 1:D:328:GLN:CG | 2.45 | 0.46 |
| 1:D:504:TYR:CE2 | 1:F:423:TYR:HE2 | 2.32 | 0.46 |
| 1:F:332:PHE:HB3 | 1:F:333:PRO:HD3 | 1.96 | 0.46 |
| 2:H:182:VAL:O | 2:H:182:VAL:HG13 | 2.15 | 0.46 |
| 2:I:48:ARG:HH21 | 2:M:574:HIS:CD2 | 2.34 | 0.46 |
| 2:I:182:VAL:HG13 | 2:I:182:VAL:O | 2.15 | 0.46 |
| 2:J:227:ASP:HB3 | 2:J:338:LEU:CD2 | 2.45 | 0.46 |
| 2:K:182:VAL:O | 2:K:182:VAL:HG13 | 2.15 | 0.46 |
| 2:M:347:LEU:C | 2:M:347:LEU:HD12 | 2.36 | 0.46 |
| 1:C:328:GLN:NE2 | 1:D:625:VAL:HA | 2.31 | 0.46 |

Continued on next page...

Not For Manuscript Review

*Continued from previous page...*

| Atom-1 | Atom-2 | Interatomic distance (Å) | Clash overlap (Å) |
|---|---|---|---|
| 1:C:385:THR:HA | 1:C:409:GLY:HA2 | 1.96 | 0.46 |
| 1:D:504:TYR:OH | 1:F:411:ASP:O | 2.32 | 0.46 |
| 1:D:599:GLY:HA3 | 1:D:626:MET:HA | 1.97 | 0.46 |
| 1:E:137:HIS:HD2 | 2:J:602:LEU:HD21 | 1.79 | 0.46 |
| 1:F:653:PHE:O | 1:F:657:VAL:HG22 | 2.16 | 0.46 |
| 2:I:309:HIS:ND1 | 2:I:310:PRO:HD3 | 2.30 | 0.46 |
| 2:K:619:ILE:HA | 2:K:656:VAL:HG11 | 1.97 | 0.46 |
| 2:L:227:ASP:HB3 | 2:L:338:LEU:CD2 | 2.45 | 0.46 |
| 2:M:309:HIS:ND1 | 2:M:310:PRO:HD3 | 2.30 | 0.46 |
| 1:A:625:VAL:HA | 1:B:328:GLN:NE2 | 2.31 | 0.46 |
| 1:B:478:VAL:HG23 | 2:K:651:LYS:CA | 2.45 | 0.46 |
| 1:D:504:TYR:CE1 | 1:F:415:ARG:N | 2.83 | 0.46 |
| 1:E:328:GLN:NE2 | 1:F:625:VAL:HA | 2.31 | 0.46 |
| 1:F:599:GLY:CA | 1:F:626:MET:HA | 2.46 | 0.46 |
| 2:H:619:ILE:HA | 2:H:656:VAL:HG11 | 1.97 | 0.46 |
| 2:K:652:MET:HG3 | 2:K:654:HIS:HB3 | 1.98 | 0.46 |
| 2:M:227:ASP:HB3 | 2:M:338:LEU:CD2 | 2.45 | 0.46 |
| 1:A:177:GLN:CG | 1:A:198:LYS:HZ2 | 2.29 | 0.46 |
| 1:A:428:LEU:C | 2:H:606:PHE:CZ | 2.89 | 0.46 |
| 1:C:599:GLY:CA | 1:C:626:MET:HA | 2.46 | 0.46 |
| 1:E:332:PHE:HB3 | 1:E:333:PRO:HD3 | 1.96 | 0.46 |
| 1:E:599:GLY:CA | 1:E:626:MET:HA | 2.46 | 0.46 |
| 1:E:625:VAL:HA | 1:F:328:GLN:NE2 | 2.31 | 0.46 |
| 1:F:180:ALA:O | 1:F:183:ARG:N | 2.45 | 0.46 |
| 2:J:182:VAL:HG13 | 2:J:182:VAL:O | 2.15 | 0.46 |
| 2:K:513:TYR:HB2 | 2:K:600:LEU:CA | 2.38 | 0.46 |
| 2:M:236:PHE:CZ | 2:M:347:LEU:HB3 | 2.51 | 0.46 |
| 1:A:415:ARG:N | 1:C:504:TYR:CE1 | 2.84 | 0.46 |
| 1:A:428:LEU:C | 2:H:606:PHE:HZ | 2.18 | 0.46 |
| 1:A:492:VAL:HG21 | 1:A:496:ARG:O | 2.16 | 0.46 |
| 1:B:599:GLY:CA | 1:B:626:MET:HA | 2.46 | 0.46 |
| 1:C:492:VAL:HG21 | 1:C:496:ARG:O | 2.16 | 0.46 |
| 1:E:428:LEU:C | 2:L:606:PHE:HZ | 2.19 | 0.46 |
| 1:E:428:LEU:C | 2:L:606:PHE:CZ | 2.89 | 0.46 |
| 1:E:599:GLY:HA3 | 1:E:626:MET:HA | 1.98 | 0.46 |
| 1:F:599:GLY:HA3 | 1:F:626:MET:HA | 1.97 | 0.46 |
| 1:F:656:LYS:O | 1:F:660:LYS:HG2 | 2.14 | 0.46 |
| 2:H:117:PRO:HD3 | 2:H:280:VAL:HG12 | 1.96 | 0.46 |
| 2:H:426:ARG:HD3 | 2:H:448:CYS:SG | 2.56 | 0.46 |
| 2:J:347:LEU:C | 2:J:347:LEU:HD12 | 2.35 | 0.46 |
| 2:L:236:PHE:CZ | 2:L:347:LEU:HB3 | 2.51 | 0.46 |

*Continued on next page...*

Not For Manuscript Review

Continued from previous page...

| Atom-1 | Atom-2 | Interatomic distance (Å) | Clash overlap (Å) |
|---|---|---|---|
| 1:A:599:GLY:HA3 | 1:A:626:MET:HA | 1.98 | 0.46 |
| 1:B:492:VAL:HG21 | 1:B:496:ARG:O | 2.16 | 0.46 |
| 1:D:184:TYR:CE2 | 1:D:201:VAL:HG21 | 2.50 | 0.46 |
| 1:E:656:LYS:O | 1:E:660:LYS:HG2 | 2.15 | 0.46 |
| 2:H:490:LEU:CD1 | 2:H:497:ARG:HE | 2.29 | 0.46 |
| 2:I:169:LYS:NZ | 2:I:173:GLY:O | 2.37 | 0.46 |
| 2:I:236:PHE:CZ | 2:I:347:LEU:HB3 | 2.51 | 0.46 |
| 2:I:619:ILE:HA | 2:I:656:VAL:HG11 | 1.97 | 0.46 |
| 2:J:253:ALA:CB | 2:J:312:THR:HA | 2.46 | 0.46 |
| 2:J:532:LYS:HE3 | 2:J:534:PHE:HE1 | 1.80 | 0.46 |
| 2:K:117:PRO:HD3 | 2:K:280:VAL:HG12 | 1.97 | 0.46 |
| 2:K:328:LEU:O | 2:K:332:THR:N | 2.40 | 0.46 |
| 2:K:490:LEU:CD1 | 2:K:497:ARG:HE | 2.29 | 0.46 |
| 1:A:608:ARG:HE | 1:A:615:LEU:HD13 | 1.81 | 0.46 |
| 1:D:492:VAL:HG21 | 1:D:496:ARG:O | 2.16 | 0.46 |
| 1:E:328:GLN:CG | 1:F:625:VAL:HG12 | 2.45 | 0.46 |
| 2:H:236:PHE:CZ | 2:H:347:LEU:HB3 | 2.51 | 0.46 |
| 2:H:513:TYR:HB2 | 2:H:600:LEU:CA | 2.38 | 0.46 |
| 2:J:652:MET:HG3 | 2:J:654:HIS:HB3 | 1.98 | 0.46 |
| 2:K:227:ASP:HB3 | 2:K:338:LEU:CD2 | 2.45 | 0.46 |
| 2:K:236:PHE:CZ | 2:K:347:LEU:HB3 | 2.51 | 0.46 |
| 2:K:253:ALA:CB | 2:K:312:THR:HA | 2.46 | 0.46 |
| 2:K:254:PRO:HG2 | 2:K:350:SER:HA | 1.98 | 0.46 |
| 2:K:309:HIS:ND1 | 2:K:310:PRO:HD3 | 2.30 | 0.46 |
| 2:K:426:ARG:HD3 | 2:K:448:CYS:SG | 2.56 | 0.46 |
| 2:L:253:ALA:CB | 2:L:312:THR:HA | 2.46 | 0.46 |
| 2:L:619:ILE:HA | 2:L:656:VAL:HG11 | 1.97 | 0.46 |
| 2:L:621:SER:HB3 | 2:L:680:LEU:HD11 | 1.97 | 0.46 |
| 2:M:253:ALA:CB | 2:M:312:THR:HA | 2.46 | 0.46 |
| 2:M:490:LEU:CD1 | 2:M:497:ARG:HE | 2.29 | 0.46 |
| 2:M:636:ASP:HA | 2:M:663:ARG:NH1 | 2.31 | 0.46 |
| 1:B:254:LYS:HB2 | 1:B:257:GLU:HB3 | 1.98 | 0.46 |
| 1:C:254:LYS:HB2 | 1:C:257:GLU:HB3 | 1.98 | 0.46 |
| 1:C:428:LEU:C | 2:J:606:PHE:CZ | 2.89 | 0.46 |
| 1:C:428:LEU:C | 2:J:606:PHE:HZ | 2.19 | 0.46 |
| 1:E:625:VAL:HG12 | 1:F:328:GLN:CG | 2.46 | 0.46 |
| 2:H:328:LEU:O | 2:H:332:THR:N | 2.40 | 0.46 |
| 2:I:253:ALA:CB | 2:I:312:THR:HA | 2.46 | 0.46 |
| 2:I:347:LEU:C | 2:I:347:LEU:HD12 | 2.35 | 0.46 |
| 2:I:532:LYS:HE3 | 2:I:534:PHE:HE1 | 1.80 | 0.46 |
| 2:J:236:PHE:CZ | 2:J:347:LEU:HB3 | 2.51 | 0.46 |

Continued on next page...

Not For Manuscript Review

*Continued from previous page...*

| Atom-1 | Atom-2 | Interatomic distance (Å) | Clash overlap (Å) |
|---|---|---|---|
| 2:J:619:ILE:HA | 2:J:656:VAL:HG11 | 1.97 | 0.46 |
| 2:L:160:LYS:N | 2:L:160:LYS:HD2 | 2.31 | 0.46 |
| 2:L:490:LEU:CD1 | 2:L:497:ARG:HE | 2.29 | 0.46 |
| 2:L:636:ASP:HA | 2:L:663:ARG:NH1 | 2.31 | 0.46 |
| 2:M:160:LYS:HD2 | 2:M:160:LYS:N | 2.31 | 0.46 |
| 2:M:169:LYS:NZ | 2:M:173:GLY:O | 2.37 | 0.46 |
| 2:M:619:ILE:HA | 2:M:656:VAL:HG11 | 1.97 | 0.46 |
| 1:A:184:TYR:CE2 | 1:A:201:VAL:HG21 | 2.50 | 0.45 |
| 1:A:282:ALA:HA | 1:A:317:LEU:HB2 | 1.98 | 0.45 |
| 1:B:478:VAL:HG23 | 2:K:651:LYS:CB | 2.46 | 0.45 |
| 1:C:142:TYR:HA | 1:C:145:SER:CB | 2.44 | 0.45 |
| 1:C:282:ALA:HA | 1:C:317:LEU:HB2 | 1.98 | 0.45 |
| 1:D:608:ARG:HE | 1:D:615:LEU:HD13 | 1.81 | 0.45 |
| 2:H:227:ASP:HB3 | 2:H:338:LEU:CD2 | 2.45 | 0.45 |
| 2:H:253:ALA:CB | 2:H:312:THR:HA | 2.46 | 0.45 |
| 2:H:309:HIS:ND1 | 2:H:310:PRO:HD3 | 2.30 | 0.45 |
| 2:H:532:LYS:HE3 | 2:H:534:PHE:HE1 | 1.80 | 0.45 |
| 2:H:621:SER:HB3 | 2:H:680:LEU:HD11 | 1.97 | 0.45 |
| 2:I:160:LYS:N | 2:I:160:LYS:HD2 | 2.31 | 0.45 |
| 2:I:652:MET:HG3 | 2:I:654:HIS:HB3 | 1.98 | 0.45 |
| 2:J:570:GLU:O | 2:J:571:ASN:C | 2.55 | 0.45 |
| 2:J:641:GLY:O | 2:J:657:ARG:NH2 | 2.49 | 0.45 |
| 2:K:621:SER:HB3 | 2:K:680:LEU:HD11 | 1.97 | 0.45 |
| 2:L:169:LYS:NZ | 2:L:173:GLY:O | 2.37 | 0.45 |
| 1:A:659:LYS:O | 1:A:663:LYS:HB2 | 2.17 | 0.45 |
| 1:B:282:ALA:HA | 1:B:317:LEU:HB2 | 1.98 | 0.45 |
| 1:B:659:LYS:O | 1:B:663:LYS:HB2 | 2.17 | 0.45 |
| 1:D:282:ALA:HA | 1:D:317:LEU:HB2 | 1.98 | 0.45 |
| 1:D:478:VAL:HG23 | 2:M:651:LYS:CA | 2.46 | 0.45 |
| 1:D:599:GLY:CA | 1:D:626:MET:HA | 2.46 | 0.45 |
| 2:H:254:PRO:HG2 | 2:H:350:SER:HA | 1.98 | 0.45 |
| 2:I:570:GLU:O | 2:I:571:ASN:C | 2.55 | 0.45 |
| 2:J:160:LYS:HD2 | 2:J:160:LYS:N | 2.31 | 0.45 |
| 2:J:426:ARG:HD3 | 2:J:448:CYS:SG | 2.56 | 0.45 |
| 2:M:621:SER:HB3 | 2:M:680:LEU:HD11 | 1.98 | 0.45 |
| 1:A:599:GLY:CA | 1:A:626:MET:HA | 2.46 | 0.45 |
| 1:B:144:ARG:NE | 1:B:144:ARG:HA | 2.32 | 0.45 |
| 1:B:171:MET:HG2 | 1:B:468:GLY:CA | 2.47 | 0.45 |
| 1:C:182:VAL:HG21 | 2:J:503:ASN:CB | 2.47 | 0.45 |
| 1:C:659:LYS:O | 1:C:663:LYS:HB2 | 2.17 | 0.45 |
| 1:D:659:LYS:O | 1:D:663:LYS:HB2 | 2.17 | 0.45 |

*Continued on next page...*

Not For Manuscript Review

Continued from previous page...

| Atom-1 | Atom-2 | Interatomic distance (Å) | Clash overlap (Å) |
|---|---|---|---|
| 1:E:492:VAL:HG21 | 1:E:496:ARG:O | 2.16 | 0.45 |
| 2:H:413:SER:O | 2:H:413:SER:OG | 2.35 | 0.45 |
| 2:I:641:GLY:O | 2:I:657:ARG:NH2 | 2.49 | 0.45 |
| 2:K:413:SER:O | 2:K:413:SER:OG | 2.35 | 0.45 |
| 2:M:650:MET:HE3 | 2:M:650:MET:HA | 1.98 | 0.45 |
| 1:A:504:TYR:CE2 | 1:E:423:TYR:HE2 | 2.33 | 0.45 |
| 1:C:144:ARG:NE | 1:C:144:ARG:HA | 2.32 | 0.45 |
| 1:C:171:MET:HG2 | 1:C:468:GLY:CA | 2.47 | 0.45 |
| 1:D:144:ARG:NE | 1:D:144:ARG:HA | 2.31 | 0.45 |
| 1:D:171:MET:HG2 | 1:D:468:GLY:CA | 2.47 | 0.45 |
| 1:E:433:LEU:HD23 | 1:E:433:LEU:HA | 1.81 | 0.45 |
| 2:I:636:ASP:HA | 2:I:663:ARG:NH1 | 2.31 | 0.45 |
| 2:J:331:LYS:HB3 | 2:J:336:LYS:HD2 | 1.99 | 0.45 |
| 2:J:636:ASP:HA | 2:J:663:ARG:NH1 | 2.31 | 0.45 |
| 2:K:532:LYS:HE3 | 2:K:534:PHE:HE1 | 1.80 | 0.45 |
| 2:M:562:VAL:HG22 | 2:M:580:VAL:HG22 | 1.99 | 0.45 |
| 1:A:171:MET:HG2 | 1:A:468:GLY:CA | 2.47 | 0.45 |
| 1:A:433:LEU:HD23 | 1:A:433:LEU:HA | 1.81 | 0.45 |
| 1:A:582:VAL:O | 1:A:589:LYS:NZ | 2.49 | 0.45 |
| 1:B:142:TYR:HA | 1:B:145:SER:CB | 2.44 | 0.45 |
| 1:E:493:ASP:OD1 | 1:E:686:ARG:NH2 | 2.47 | 0.45 |
| 1:F:492:VAL:HG21 | 1:F:496:ARG:O | 2.16 | 0.45 |
| 2:H:160:LYS:HD2 | 2:H:160:LYS:N | 2.31 | 0.45 |
| 2:I:331:LYS:HB3 | 2:I:336:LYS:HD2 | 1.98 | 0.45 |
| 2:I:426:ARG:HD3 | 2:I:448:CYS:SG | 2.56 | 0.45 |
| 2:J:169:LYS:NZ | 2:J:173:GLY:O | 2.37 | 0.45 |
| 2:K:160:LYS:N | 2:K:160:LYS:HD2 | 2.31 | 0.45 |
| 2:L:650:MET:HE3 | 2:L:650:MET:HA | 1.98 | 0.45 |
| 1:A:625:VAL:HG23 | 1:A:627:GLY:H | 1.82 | 0.45 |
| 1:B:415:ARG:N | 1:F:504:TYR:CE1 | 2.85 | 0.45 |
| 1:B:504:TYR:CE1 | 1:D:415:ARG:N | 2.84 | 0.45 |
| 1:B:608:ARG:HE | 1:B:615:LEU:HD13 | 1.81 | 0.45 |
| 1:D:219:ARG:O | 1:D:382:ARG:NH2 | 2.50 | 0.45 |
| 1:D:582:VAL:O | 1:D:589:LYS:NZ | 2.49 | 0.45 |
| 1:E:254:LYS:HB2 | 1:E:257:GLU:HB3 | 1.99 | 0.45 |
| 2:J:327:LYS:HE2 | 2:J:331:LYS:NZ | 2.32 | 0.45 |
| 2:J:621:SER:HB3 | 2:J:680:LEU:HD11 | 1.97 | 0.45 |
| 2:L:562:VAL:HG22 | 2:L:580:VAL:HG22 | 1.99 | 0.45 |
| 1:A:144:ARG:NE | 1:A:144:ARG:HA | 2.32 | 0.45 |
| 1:B:363:CYS:SG | 1:B:386:ILE:HG22 | 2.57 | 0.45 |
| 1:D:625:VAL:HG23 | 1:D:627:GLY:H | 1.82 | 0.45 |

Continued on next page...

Not For Manuscript Review

Continued from previous page...

| Atom-1 | Atom-2 | Interatomic distance (Å) | Clash overlap (Å) |
|---|---|---|---|
| 1:F:254:LYS:HB2 | 1:F:257:GLU:HB3 | 1.99 | 0.45 |
| 1:F:282:ALA:HA | 1:F:317:LEU:HB2 | 1.98 | 0.45 |
| 1:F:493:ASP:OD1 | 1:F:686:ARG:NH2 | 2.47 | 0.45 |
| 2:H:173:GLY:HA3 | 2:H:202:PHE:HZ | 1.82 | 0.45 |
| 2:I:327:LYS:HE2 | 2:I:331:LYS:NZ | 2.32 | 0.45 |
| 2:I:490:LEU:CD1 | 2:I:497:ARG:HE | 2.29 | 0.45 |
| 2:J:490:LEU:CD1 | 2:J:497:ARG:HE | 2.29 | 0.45 |
| 2:K:512:PHE:O | 2:K:519:VAL:N | 2.37 | 0.45 |
| 2:M:173:GLY:HA3 | 2:M:202:PHE:HZ | 1.82 | 0.45 |
| 1:A:144:ARG:HG3 | 2:L:606:PHE:CD2 | 2.52 | 0.45 |
| 1:A:219:ARG:O | 1:A:382:ARG:NH2 | 2.50 | 0.45 |
| 1:C:363:CYS:SG | 1:C:386:ILE:HG22 | 2.57 | 0.45 |
| 1:E:219:ARG:O | 1:E:382:ARG:NH2 | 2.50 | 0.45 |
| 1:E:282:ALA:HA | 1:E:317:LEU:HB2 | 1.98 | 0.45 |
| 1:F:428:LEU:CB | 2:M:606:PHE:CZ | 2.88 | 0.45 |
| 2:I:621:SER:HB3 | 2:I:680:LEU:HD11 | 1.97 | 0.45 |
| 2:J:254:PRO:HG2 | 2:J:350:SER:HA | 1.98 | 0.45 |
| 2:K:173:GLY:HA3 | 2:K:202:PHE:HZ | 1.82 | 0.45 |
| 2:K:570:GLU:O | 2:K:571:ASN:C | 2.55 | 0.45 |
| 2:L:254:PRO:HG2 | 2:L:350:SER:HA | 1.98 | 0.45 |
| 2:L:327:LYS:HE2 | 2:L:331:LYS:NZ | 2.32 | 0.45 |
| 2:L:485:ALA:HB2 | 2:L:512:PHE:HE2 | 1.82 | 0.45 |
| 2:M:327:LYS:HE2 | 2:M:331:LYS:NZ | 2.32 | 0.45 |
| 2:M:426:ARG:HD3 | 2:M:448:CYS:SG | 2.56 | 0.45 |
| 2:M:485:ALA:HB2 | 2:M:512:PHE:HE2 | 1.82 | 0.45 |
| 1:B:625:VAL:HG23 | 1:B:627:GLY:H | 1.82 | 0.45 |
| 1:C:219:ARG:O | 1:C:382:ARG:NH2 | 2.50 | 0.45 |
| 1:C:608:ARG:HE | 1:C:615:LEU:HD13 | 1.81 | 0.45 |
| 1:C:625:VAL:HG23 | 1:C:627:GLY:H | 1.82 | 0.45 |
| 1:D:473:ASP:C | 1:D:480:LYS:HE3 | 2.37 | 0.45 |
| 1:E:144:ARG:HG3 | 2:J:606:PHE:CD2 | 2.52 | 0.45 |
| 1:F:219:ARG:O | 1:F:382:ARG:NH2 | 2.50 | 0.45 |
| 1:F:363:CYS:SG | 1:F:386:ILE:HG22 | 2.57 | 0.45 |
| 2:H:562:VAL:HG22 | 2:H:580:VAL:HG22 | 1.99 | 0.45 |
| 2:H:570:GLU:O | 2:H:571:ASN:C | 2.55 | 0.45 |
| 2:H:636:ASP:HA | 2:H:663:ARG:NH1 | 2.31 | 0.45 |
| 2:I:254:PRO:HG2 | 2:I:350:SER:HA | 1.98 | 0.45 |
| 2:L:413:SER:O | 2:L:413:SER:OG | 2.35 | 0.45 |
| 2:L:652:MET:HG3 | 2:L:654:HIS:HB3 | 1.98 | 0.45 |
| 2:M:254:PRO:HG2 | 2:M:350:SER:HA | 1.98 | 0.45 |
| 2:M:513:TYR:HB2 | 2:M:600:LEU:CA | 2.38 | 0.45 |

Continued on next page...

Not For Manuscript Review

Continued from previous page...

| Atom-1 | Atom-2 | Interatomic distance (Å) | Clash overlap (Å) |
|---|---|---|---|
| 2:M:570:GLU:O | 2:M:571:ASN:C | 2.55 | 0.45 |
| 2:M:650:MET:HE2 | 2:M:651:LYS:CE | 2.23 | 0.45 |
| 2:M:652:MET:HG3 | 2:M:654:HIS:HB3 | 1.98 | 0.45 |
| 1:A:160:SER:HA | 1:A:164:TYR:CG | 2.52 | 0.45 |
| 1:A:473:ASP:C | 1:A:480:LYS:HE3 | 2.37 | 0.45 |
| 1:B:219:ARG:O | 1:B:382:ARG:NH2 | 2.50 | 0.45 |
| 1:B:582:VAL:O | 1:B:589:LYS:NZ | 2.49 | 0.45 |
| 1:D:160:SER:HA | 1:D:164:TYR:CG | 2.52 | 0.45 |
| 1:E:182:VAL:HG21 | 2:L:503:ASN:CB | 2.47 | 0.45 |
| 1:E:363:CYS:SG | 1:E:386:ILE:HG22 | 2.57 | 0.45 |
| 1:F:608:ARG:HE | 1:F:615:LEU:HD13 | 1.81 | 0.45 |
| 2:I:485:ALA:HB2 | 2:I:512:PHE:HE2 | 1.82 | 0.45 |
| 2:J:485:ALA:HB2 | 2:J:512:PHE:HE2 | 1.82 | 0.45 |
| 2:L:173:GLY:HA3 | 2:L:202:PHE:HZ | 1.82 | 0.45 |
| 2:L:426:ARG:HD3 | 2:L:448:CYS:SG | 2.56 | 0.45 |
| 2:L:513:TYR:HB2 | 2:L:600:LEU:CA | 2.38 | 0.45 |
| 2:M:413:SER:O | 2:M:413:SER:OG | 2.35 | 0.45 |
| 1:D:433:LEU:HD23 | 1:D:433:LEU:HA | 1.81 | 0.44 |
| 1:F:171:MET:HG2 | 1:F:468:GLY:CA | 2.47 | 0.44 |
| 1:F:433:LEU:HD23 | 1:F:433:LEU:HA | 1.81 | 0.44 |
| 2:H:512:PHE:O | 2:H:519:VAL:N | 2.37 | 0.44 |
| 2:I:562:VAL:HG22 | 2:I:580:VAL:HG22 | 1.99 | 0.44 |
| 2:J:562:VAL:HG22 | 2:J:580:VAL:HG22 | 1.99 | 0.44 |
| 2:K:562:VAL:HG22 | 2:K:580:VAL:HG22 | 1.99 | 0.44 |
| 2:K:641:GLY:O | 2:K:657:ARG:NH2 | 2.49 | 0.44 |
| 1:A:254:LYS:HB2 | 1:A:257:GLU:HB3 | 1.99 | 0.44 |
| 1:B:160:SER:HA | 1:B:164:TYR:CG | 2.52 | 0.44 |
| 1:C:473:ASP:C | 1:C:480:LYS:HE3 | 2.37 | 0.44 |
| 1:C:582:VAL:O | 1:C:589:LYS:NZ | 2.49 | 0.44 |
| 1:D:254:LYS:HB2 | 1:D:257:GLU:HB3 | 1.99 | 0.44 |
| 1:F:182:VAL:HG21 | 2:M:503:ASN:HB2 | 1.98 | 0.44 |
| 2:H:641:GLY:O | 2:H:657:ARG:NH2 | 2.49 | 0.44 |
| 2:I:571:ASN:OD1 | 2:I:572:GLY:N | 2.48 | 0.44 |
| 2:J:571:ASN:OD1 | 2:J:572:GLY:N | 2.48 | 0.44 |
| 2:K:571:ASN:OD1 | 2:K:572:GLY:N | 2.48 | 0.44 |
| 2:K:636:ASP:HA | 2:K:663:ARG:NH1 | 2.31 | 0.44 |
| 2:L:570:GLU:O | 2:L:571:ASN:C | 2.55 | 0.44 |
| 2:L:641:GLY:O | 2:L:657:ARG:NH2 | 2.49 | 0.44 |
| 1:B:473:ASP:C | 1:B:480:LYS:HE3 | 2.37 | 0.44 |
| 1:C:160:SER:HA | 1:C:164:TYR:CG | 2.52 | 0.44 |
| 1:E:171:MET:HG2 | 1:E:468:GLY:CA | 2.47 | 0.44 |

Continued on next page...

Not For Manuscript Review

*Continued from previous page...*

| Atom-1 | Atom-2 | Interatomic distance (Å) | Clash overlap (Å) |
|---|---|---|---|
| 2:H:485:ALA:HB2 | 2:H:512:PHE:HE2 | 1.81 | 0.44 |
| 2:H:571:ASN:OD1 | 2:H:572:GLY:N | 2.48 | 0.44 |
| 2:L:331:LYS:HB3 | 2:L:336:LYS:HD2 | 1.98 | 0.44 |
| 2:L:552:THR:OG1 | 2:L:563:ASN:OD1 | 2.30 | 0.44 |
| 2:L:650:MET:HE2 | 2:L:651:LYS:CE | 2.23 | 0.44 |
| 2:M:331:LYS:HB3 | 2:M:336:LYS:HD2 | 1.98 | 0.44 |
| 1:A:500:PHE:CE2 | 1:A:540:PHE:CD1 | 3.06 | 0.44 |
| 1:A:578:LEU:O | 1:A:582:VAL:HG23 | 2.18 | 0.44 |
| 1:E:473:ASP:C | 1:E:480:LYS:HE3 | 2.37 | 0.44 |
| 1:E:608:ARG:HE | 1:E:615:LEU:HD13 | 1.82 | 0.44 |
| 1:F:659:LYS:O | 1:F:663:LYS:HB2 | 2.17 | 0.44 |
| 2:H:48:ARG:HH21 | 2:J:574:HIS:CD2 | 2.35 | 0.44 |
| 2:I:253:ALA:HB1 | 2:I:312:THR:HA | 2.00 | 0.44 |
| 2:J:253:ALA:HB1 | 2:J:312:THR:HA | 2.00 | 0.44 |
| 2:M:641:GLY:O | 2:M:657:ARG:NH2 | 2.49 | 0.44 |
| 1:A:363:CYS:SG | 1:A:386:ILE:HG22 | 2.57 | 0.44 |
| 1:D:363:CYS:SG | 1:D:386:ILE:HG22 | 2.57 | 0.44 |
| 1:D:500:PHE:CE2 | 1:D:540:PHE:CD1 | 3.06 | 0.44 |
| 1:E:142:TYR:HA | 1:E:145:SER:CB | 2.44 | 0.44 |
| 1:E:659:LYS:O | 1:E:663:LYS:HB2 | 2.17 | 0.44 |
| 1:F:137:HIS:HA | 2:I:602:LEU:HD11 | 1.99 | 0.44 |
| 1:F:142:TYR:O | 1:F:143:GLU:CB | 2.65 | 0.44 |
| 2:I:173:GLY:HA3 | 2:I:202:PHE:HZ | 1.82 | 0.44 |
| 2:J:385:PRO:HG2 | 2:J:420:GLU:CD | 2.38 | 0.44 |
| 2:M:552:THR:OG1 | 2:M:563:ASN:OD1 | 2.30 | 0.44 |
| 1:C:433:LEU:HD23 | 1:C:433:LEU:HA | 1.81 | 0.44 |
| 1:D:578:LEU:O | 1:D:582:VAL:HG23 | 2.18 | 0.44 |
| 1:D:629:THR:HA | 1:D:658:LYS:NZ | 2.33 | 0.44 |
| 1:E:142:TYR:O | 1:E:143:GLU:CB | 2.66 | 0.44 |
| 2:H:574:HIS:CD2 | 2:L:48:ARG:HH21 | 2.35 | 0.44 |
| 2:I:513:TYR:HB2 | 2:I:600:LEU:CA | 2.38 | 0.44 |
| 2:J:48:ARG:HH21 | 2:L:574:HIS:CD2 | 2.36 | 0.44 |
| 2:J:429:LEU:HD13 | 2:J:429:LEU:HA | 1.90 | 0.44 |
| 2:J:446:ARG:HD2 | 2:J:446:ARG:HA | 1.83 | 0.44 |
| 2:K:485:ALA:HB2 | 2:K:512:PHE:HE2 | 1.82 | 0.44 |
| 2:K:490:LEU:HD13 | 2:K:490:LEU:HA | 1.86 | 0.44 |
| 1:A:274:ASN:ND2 | 1:A:439:ALA:HB2 | 2.33 | 0.44 |
| 1:A:488:ILE:HG13 | 1:A:489:ALA:N | 2.33 | 0.44 |
| 1:A:629:THR:HA | 1:A:658:LYS:NZ | 2.33 | 0.44 |
| 1:C:144:ARG:HG3 | 2:H:606:PHE:CD2 | 2.53 | 0.44 |
| 1:D:274:ASN:ND2 | 1:D:439:ALA:HB2 | 2.33 | 0.44 |

*Continued on next page...*

Not For Manuscript Review

*Continued from previous page...*

| Atom-1 | Atom-2 | Interatomic distance (Å) | Clash overlap (Å) |
|---|---|---|---|
| 1:E:488:ILE:HG13 | 1:E:489:ALA:N | 2.33 | 0.44 |
| 1:F:142:TYR:HA | 1:F:145:SER:CB | 2.44 | 0.44 |
| 1:F:274:ASN:ND2 | 1:F:439:ALA:HB2 | 2.33 | 0.44 |
| 1:F:473:ASP:C | 1:F:480:LYS:HE3 | 2.37 | 0.44 |
| 1:F:488:ILE:HG13 | 1:F:489:ALA:N | 2.33 | 0.44 |
| 1:F:500:PHE:CE2 | 1:F:540:PHE:CD1 | 3.06 | 0.44 |
| 2:H:327:LYS:HE2 | 2:H:331:LYS:NZ | 2.32 | 0.44 |
| 2:H:566:THR:HA | 2:H:575:HIS:O | 2.18 | 0.44 |
| 2:I:385:PRO:HG2 | 2:I:420:GLU:CD | 2.38 | 0.44 |
| 2:J:169:LYS:HB3 | 2:J:170:ALA:H | 1.53 | 0.44 |
| 2:J:173:GLY:HA3 | 2:J:202:PHE:HZ | 1.82 | 0.44 |
| 2:J:196:LYS:HD3 | 2:J:196:LYS:HA | 1.78 | 0.44 |
| 2:K:327:LYS:HE2 | 2:K:331:LYS:NZ | 2.32 | 0.44 |
| 2:K:331:LYS:HB3 | 2:K:336:LYS:HD2 | 1.98 | 0.44 |
| 2:K:566:THR:HA | 2:K:575:HIS:O | 2.18 | 0.44 |
| 2:L:565:PHE:HB2 | 2:L:577:VAL:CG2 | 2.38 | 0.44 |
| 2:L:607:GLY:O | 2:L:609:SER:N | 2.51 | 0.44 |
| 2:M:607:GLY:O | 2:M:609:SER:N | 2.51 | 0.44 |
| 1:B:500:PHE:CE2 | 1:B:540:PHE:CD1 | 3.06 | 0.44 |
| 1:B:560:PHE:O | 3:G:805:BTI:H62 | 2.17 | 0.44 |
| 1:C:578:LEU:O | 1:C:582:VAL:HG23 | 2.18 | 0.44 |
| 1:D:177:GLN:CG | 1:D:198:LYS:HZ2 | 2.31 | 0.44 |
| 1:D:341:ILE:O | 1:D:345:GLU:HG2 | 2.18 | 0.44 |
| 1:D:488:ILE:HG13 | 1:D:489:ALA:N | 2.33 | 0.44 |
| 1:E:184:TYR:O | 1:E:188:GLY:HA2 | 2.18 | 0.44 |
| 1:E:341:ILE:O | 1:E:345:GLU:HG2 | 2.18 | 0.44 |
| 1:E:500:PHE:CE2 | 1:E:540:PHE:CD1 | 3.06 | 0.44 |
| 1:F:341:ILE:O | 1:F:345:GLU:HG2 | 2.18 | 0.44 |
| 1:F:385:THR:HB | 1:F:387:PHE:CE2 | 2.53 | 0.44 |
| 2:H:221:GLU:HB3 | 2:H:286:GLU:HG3 | 1.99 | 0.44 |
| 2:H:490:LEU:HD13 | 2:H:490:LEU:HA | 1.86 | 0.44 |
| 2:I:196:LYS:HD3 | 2:I:196:LYS:HA | 1.78 | 0.44 |
| 2:I:221:GLU:HB3 | 2:I:286:GLU:HG3 | 2.00 | 0.44 |
| 2:K:652:MET:HG3 | 2:K:654:HIS:HD2 | 1.83 | 0.44 |
| 1:A:341:ILE:O | 1:A:345:GLU:HG2 | 2.18 | 0.44 |
| 1:B:578:LEU:O | 1:B:582:VAL:HG23 | 2.18 | 0.44 |
| 1:C:500:PHE:CE2 | 1:C:540:PHE:CD1 | 3.06 | 0.44 |
| 1:E:274:ASN:ND2 | 1:E:439:ALA:HB2 | 2.33 | 0.44 |
| 1:E:582:VAL:O | 1:E:589:LYS:NZ | 2.49 | 0.44 |
| 2:H:279:TYR:OH | 2:H:282:ALA:O | 2.35 | 0.44 |
| 2:H:331:LYS:HB3 | 2:H:336:LYS:HD2 | 1.99 | 0.44 |

*Continued on next page...*

Not For Manuscript Review

*Continued from previous page...*

| Atom-1 | Atom-2 | Interatomic distance (Å) | Clash overlap (Å) |
|---|---|---|---|
| 2:H:652:MET:HG3 | 2:H:654:HIS:HD2 | 1.83 | 0.44 |
| 2:J:221:GLU:HB3 | 2:J:286:GLU:HG3 | 2.00 | 0.44 |
| 2:M:429:LEU:O | 2:M:441:ILE:HD11 | 2.18 | 0.44 |
| 1:A:142:TYR:O | 1:A:143:GLU:CB | 2.65 | 0.43 |
| 1:A:170:ARG:HB2 | 1:A:473:ASP:OD1 | 2.18 | 0.43 |
| 1:A:226:ASP:HA | 1:A:229:GLU:CD | 2.38 | 0.43 |
| 1:A:385:THR:HB | 1:A:387:PHE:CE2 | 2.53 | 0.43 |
| 1:B:170:ARG:HB2 | 1:B:473:ASP:OD1 | 2.18 | 0.43 |
| 1:B:629:THR:HA | 1:B:658:LYS:NZ | 2.33 | 0.43 |
| 1:C:274:ASN:ND2 | 1:C:439:ALA:HB2 | 2.33 | 0.43 |
| 1:D:170:ARG:HB2 | 1:D:473:ASP:OD1 | 2.18 | 0.43 |
| 1:D:385:THR:HB | 1:D:387:PHE:CE2 | 2.53 | 0.43 |
| 1:E:385:THR:HB | 1:E:387:PHE:CE2 | 2.53 | 0.43 |
| 2:I:169:LYS:HB3 | 2:I:170:ALA:H | 1.52 | 0.43 |
| 2:I:314:GLU:HB3 | 2:I:414:TRP:HD1 | 1.83 | 0.43 |
| 2:J:413:SER:O | 2:J:413:SER:OG | 2.35 | 0.43 |
| 2:J:513:TYR:HB2 | 2:J:600:LEU:CA | 2.38 | 0.43 |
| 2:K:221:GLU:HB3 | 2:K:286:GLU:HG3 | 2.00 | 0.43 |
| 2:L:429:LEU:O | 2:L:441:ILE:HD11 | 2.18 | 0.43 |
| 2:M:253:ALA:HB1 | 2:M:312:THR:HA | 2.00 | 0.43 |
| 1:A:182:VAL:HG21 | 2:H:503:ASN:CB | 2.48 | 0.43 |
| 1:B:433:LEU:HD23 | 1:B:433:LEU:HA | 1.81 | 0.43 |
| 1:C:629:THR:HA | 1:C:658:LYS:NZ | 2.33 | 0.43 |
| 1:D:226:ASP:HA | 1:D:229:GLU:CD | 2.39 | 0.43 |
| 1:E:625:VAL:HG23 | 1:E:627:GLY:H | 1.82 | 0.43 |
| 1:F:184:TYR:O | 1:F:188:GLY:HA2 | 2.18 | 0.43 |
| 1:F:582:VAL:O | 1:F:589:LYS:NZ | 2.49 | 0.43 |
| 2:I:376:ARG:HB2 | 2:I:433:LYS:H | 1.83 | 0.43 |
| 2:I:413:SER:O | 2:I:413:SER:OG | 2.35 | 0.43 |
| 2:J:314:GLU:HB3 | 2:J:414:TRP:HD1 | 1.83 | 0.43 |
| 2:K:279:TYR:OH | 2:K:282:ALA:O | 2.35 | 0.43 |
| 2:L:196:LYS:HD3 | 2:L:196:LYS:HA | 1.78 | 0.43 |
| 2:L:253:ALA:HB1 | 2:L:312:THR:HA | 2.00 | 0.43 |
| 2:M:565:PHE:HB2 | 2:M:577:VAL:CG2 | 2.38 | 0.43 |
| 1:A:149:SER:N | 1:A:150:PRO:HD3 | 2.33 | 0.43 |
| 1:A:184:TYR:HD1 | 1:A:191:VAL:HG21 | 1.83 | 0.43 |
| 1:B:149:SER:N | 1:B:150:PRO:HD3 | 2.33 | 0.43 |
| 1:D:142:TYR:O | 1:D:143:GLU:CB | 2.66 | 0.43 |
| 1:E:160:SER:HA | 1:E:164:TYR:CG | 2.52 | 0.43 |
| 1:E:629:THR:HA | 1:E:658:LYS:NZ | 2.33 | 0.43 |
| 1:F:510:CYS:HB3 | 1:F:540:PHE:CD2 | 2.53 | 0.43 |

*Continued on next page...*

Not For Manuscript Review

Continued from previous page...

| Atom-1 | Atom-2 | Interatomic distance (Å) | Clash overlap (Å) |
|---|---|---|---|
| 1:F:629:THR:HA | 1:F:658:LYS:NZ | 2.33 | 0.43 |
| 2:H:429:LEU:HB2 | 2:H:445:LYS:HD3 | 2.00 | 0.43 |
| 2:K:429:LEU:HB2 | 2:K:445:LYS:HD3 | 2.00 | 0.43 |
| 2:L:652:MET:HG3 | 2:L:654:HIS:HD2 | 1.83 | 0.43 |
| 1:B:184:TYR:CD1 | 1:B:191:VAL:HG21 | 2.53 | 0.43 |
| 1:B:274:ASN:ND2 | 1:B:439:ALA:HB2 | 2.33 | 0.43 |
| 1:B:341:ILE:O | 1:B:345:GLU:HG2 | 2.18 | 0.43 |
| 1:C:149:SER:N | 1:C:150:PRO:HD3 | 2.33 | 0.43 |
| 1:C:170:ARG:HB2 | 1:C:473:ASP:OD1 | 2.19 | 0.43 |
| 1:D:184:TYR:HD1 | 1:D:191:VAL:HG21 | 1.84 | 0.43 |
| 1:D:184:TYR:CD1 | 1:D:191:VAL:HG21 | 2.53 | 0.43 |
| 1:D:493:ASP:OD1 | 1:D:686:ARG:NH2 | 2.47 | 0.43 |
| 1:E:184:TYR:HD1 | 1:E:191:VAL:HG21 | 1.84 | 0.43 |
| 1:E:428:LEU:HB3 | 2:L:606:PHE:CE1 | 2.53 | 0.43 |
| 1:E:510:CYS:HB3 | 1:E:540:PHE:CD2 | 2.54 | 0.43 |
| 2:H:385:PRO:HG2 | 2:H:420:GLU:CD | 2.38 | 0.43 |
| 2:I:429:LEU:HD13 | 2:I:429:LEU:HA | 1.90 | 0.43 |
| 2:I:574:HIS:CD2 | 2:K:48:ARG:HH21 | 2.36 | 0.43 |
| 2:J:376:ARG:HB2 | 2:J:433:LYS:H | 1.83 | 0.43 |
| 2:K:314:GLU:HB3 | 2:K:414:TRP:HD1 | 1.82 | 0.43 |
| 2:K:385:PRO:HG2 | 2:K:420:GLU:CD | 2.38 | 0.43 |
| 2:K:477:THR:HG22 | 2:K:480:VAL:HG23 | 2.01 | 0.43 |
| 2:M:376:ARG:HB2 | 2:M:433:LYS:H | 1.83 | 0.43 |
| 1:A:184:TYR:CD1 | 1:A:191:VAL:HG21 | 2.53 | 0.43 |
| 1:A:493:ASP:OD1 | 1:A:686:ARG:NH2 | 2.47 | 0.43 |
| 1:A:654:LYS:O | 1:A:658:LYS:HG2 | 2.19 | 0.43 |
| 1:B:385:THR:HB | 1:B:387:PHE:CE2 | 2.53 | 0.43 |
| 1:C:341:ILE:O | 1:C:345:GLU:HG2 | 2.18 | 0.43 |
| 1:D:149:SER:N | 1:D:150:PRO:HD3 | 2.33 | 0.43 |
| 1:F:170:ARG:HB2 | 1:F:473:ASP:OD1 | 2.18 | 0.43 |
| 1:F:184:TYR:HD1 | 1:F:191:VAL:HG21 | 1.84 | 0.43 |
| 1:F:625:VAL:HG23 | 1:F:627:GLY:H | 1.82 | 0.43 |
| 2:I:566:THR:HA | 2:I:575:HIS:O | 2.18 | 0.43 |
| 2:K:429:LEU:O | 2:K:441:ILE:HD11 | 2.18 | 0.43 |
| 2:L:259:SER:O | 2:L:263:ARG:HG2 | 2.19 | 0.43 |
| 2:L:376:ARG:HB2 | 2:L:433:LYS:H | 1.83 | 0.43 |
| 2:M:196:LYS:HD3 | 2:M:196:LYS:HA | 1.78 | 0.43 |
| 2:M:259:SER:O | 2:M:263:ARG:HG2 | 2.19 | 0.43 |
| 1:A:184:TYR:O | 1:A:188:GLY:HA2 | 2.18 | 0.43 |
| 1:C:184:TYR:CD1 | 1:C:191:VAL:HG21 | 2.53 | 0.43 |
| 1:C:385:THR:HB | 1:C:387:PHE:CE2 | 2.53 | 0.43 |

Continued on next page...

Not For Manuscript Review

Continued from previous page...

| Atom-1 | Atom-2 | Interatomic distance (Å) | Clash overlap (Å) |
|---|---|---|---|
| 1:C:388:LEU:HD23 | 1:C:388:LEU:HA | 1.91 | 0.43 |
| 1:D:216:HIS:CG | 1:D:223:LEU:HD12 | 2.54 | 0.43 |
| 1:D:654:LYS:O | 1:D:658:LYS:HG2 | 2.19 | 0.43 |
| 1:F:160:SER:HA | 1:F:164:TYR:CG | 2.52 | 0.43 |
| 2:H:314:GLU:HB3 | 2:H:414:TRP:HD1 | 1.83 | 0.43 |
| 2:H:429:LEU:O | 2:H:441:ILE:HD11 | 2.18 | 0.43 |
| 2:H:477:THR:HG22 | 2:H:480:VAL:HG23 | 2.01 | 0.43 |
| 2:I:180:LYS:HD3 | 2:I:180:LYS:HA | 1.90 | 0.43 |
| 2:J:566:THR:HA | 2:J:575:HIS:O | 2.18 | 0.43 |
| 2:K:165:PRO:HD2 | 2:K:212:ARG:HG2 | 2.01 | 0.43 |
| 2:L:566:THR:HA | 2:L:575:HIS:O | 2.18 | 0.43 |
| 2:M:165:PRO:HD2 | 2:M:212:ARG:HG2 | 2.01 | 0.43 |
| 2:M:477:THR:HG22 | 2:M:480:VAL:HG23 | 2.01 | 0.43 |
| 2:M:652:MET:HG3 | 2:M:654:HIS:HD2 | 1.83 | 0.43 |
| 1:A:216:HIS:CG | 1:A:223:LEU:HD12 | 2.54 | 0.43 |
| 1:A:510:CYS:HB3 | 1:A:540:PHE:CD2 | 2.54 | 0.43 |
| 1:B:144:ARG:HG3 | 2:K:606:PHE:CE2 | 2.53 | 0.43 |
| 1:B:226:ASP:HA | 1:B:229:GLU:CD | 2.38 | 0.43 |
| 1:C:137:HIS:HD2 | 2:H:602:LEU:HD21 | 1.82 | 0.43 |
| 1:C:411:ASP:HB3 | 1:C:413:HIS:CE1 | 2.54 | 0.43 |
| 1:C:488:ILE:HG13 | 1:C:489:ALA:N | 2.33 | 0.43 |
| 1:E:144:ARG:HA | 1:E:144:ARG:NE | 2.32 | 0.43 |
| 1:E:149:SER:N | 1:E:150:PRO:HD3 | 2.33 | 0.43 |
| 1:E:170:ARG:HB2 | 1:E:473:ASP:OD1 | 2.18 | 0.43 |
| 1:E:226:ASP:HA | 1:E:229:GLU:CD | 2.39 | 0.43 |
| 1:F:226:ASP:HA | 1:F:229:GLU:CD | 2.38 | 0.43 |
| 1:F:510:CYS:SG | 1:F:523:ILE:HG12 | 2.59 | 0.43 |
| 2:H:165:PRO:HD2 | 2:H:212:ARG:HG2 | 2.01 | 0.43 |
| 2:I:315:VAL:HG22 | 2:I:315:VAL:O | 2.18 | 0.43 |
| 2:J:315:VAL:HG22 | 2:J:315:VAL:O | 2.18 | 0.43 |
| 2:K:124:MET:HE1 | 2:K:301:MET:HB3 | 2.00 | 0.43 |
| 2:K:253:ALA:HB1 | 2:K:312:THR:HA | 2.00 | 0.43 |
| 2:L:446:ARG:HD2 | 2:L:446:ARG:HA | 1.83 | 0.43 |
| 2:L:477:THR:HG22 | 2:L:480:VAL:HG23 | 2.01 | 0.43 |
| 2:L:488:TRP:NE1 | 2:L:582:THR:O | 2.52 | 0.43 |
| 2:M:488:TRP:NE1 | 2:M:582:THR:O | 2.52 | 0.43 |
| 2:M:566:THR:HA | 2:M:575:HIS:O | 2.18 | 0.43 |
| 1:C:226:ASP:HA | 1:C:229:GLU:CD | 2.38 | 0.43 |
| 1:C:597:SER:O | 1:C:623:ILE:HA | 2.19 | 0.43 |
| 1:E:170:ARG:CB | 1:E:473:ASP:OD1 | 2.67 | 0.43 |
| 1:E:510:CYS:SG | 1:E:523:ILE:HG12 | 2.59 | 0.43 |

Continued on next page...

Not For Manuscript Review

Continued from previous page...

| Atom-1 | Atom-2 | Interatomic distance (Å) | Clash overlap (Å) |
|---|---|---|---|
| 1:F:144:ARG:NE | 1:F:144:ARG:HA | 2.32 | 0.43 |
| 1:F:184:TYR:CD1 | 1:F:191:VAL:HG21 | 2.53 | 0.43 |
| 2:I:240:CYS:HA | 2:I:251:GLU:HG3 | 2.01 | 0.43 |
| 2:I:259:SER:O | 2:I:263:ARG:HG2 | 2.19 | 0.43 |
| 2:I:483:MET:SD | 2:I:569:PHE:HZ | 2.42 | 0.43 |
| 2:J:240:CYS:HA | 2:J:251:GLU:HG3 | 2.01 | 0.43 |
| 2:J:650:MET:CE | 2:J:651:LYS:HZ2 | 2.22 | 0.43 |
| 2:L:139:VAL:O | 2:L:141:VAL:HG23 | 2.19 | 0.43 |
| 2:L:165:PRO:HD2 | 2:L:212:ARG:HG2 | 2.01 | 0.43 |
| 2:L:328:LEU:O | 2:L:332:THR:N | 2.40 | 0.43 |
| 2:L:385:PRO:HG2 | 2:L:420:GLU:CD | 2.38 | 0.43 |
| 2:M:328:LEU:O | 2:M:332:THR:N | 2.40 | 0.43 |
| 2:M:429:LEU:CB | 2:M:445:LYS:HD3 | 2.49 | 0.43 |
| 2:M:446:ARG:HD2 | 2:M:446:ARG:HA | 1.83 | 0.43 |
| 1:B:184:TYR:HD1 | 1:B:191:VAL:HG21 | 1.84 | 0.43 |
| 1:B:488:ILE:HG13 | 1:B:489:ALA:N | 2.33 | 0.43 |
| 1:D:510:CYS:HB3 | 1:D:540:PHE:CD2 | 2.54 | 0.43 |
| 1:E:184:TYR:CD1 | 1:E:191:VAL:HG21 | 2.53 | 0.43 |
| 1:F:149:SER:N | 1:F:150:PRO:HD3 | 2.33 | 0.43 |
| 1:F:170:ARG:CB | 1:F:473:ASP:OD1 | 2.67 | 0.43 |
| 1:F:578:LEU:O | 1:F:582:VAL:HG23 | 2.18 | 0.43 |
| 2:H:253:ALA:HB1 | 2:H:312:THR:HA | 2.00 | 0.43 |
| 2:I:429:LEU:O | 2:I:441:ILE:HD11 | 2.18 | 0.43 |
| 2:J:259:SER:O | 2:J:263:ARG:HG2 | 2.19 | 0.43 |
| 2:J:483:MET:SD | 2:J:569:PHE:HZ | 2.42 | 0.43 |
| 2:K:62:ILE:HD12 | 2:K:71:TYR:H | 1.84 | 0.43 |
| 2:K:483:MET:SD | 2:K:569:PHE:HZ | 2.42 | 0.43 |
| 2:L:315:VAL:HG22 | 2:L:315:VAL:O | 2.18 | 0.43 |
| 2:L:429:LEU:CB | 2:L:445:LYS:HD3 | 2.49 | 0.43 |
| 2:M:139:VAL:O | 2:M:141:VAL:HG23 | 2.19 | 0.43 |
| 2:M:385:PRO:HG2 | 2:M:420:GLU:CD | 2.38 | 0.43 |
| 1:B:135:TYR:HE2 | 1:D:220:GLY:HA2 | 1.82 | 0.43 |
| 1:B:170:ARG:CB | 1:B:473:ASP:OD1 | 2.67 | 0.43 |
| 1:B:388:LEU:HD23 | 1:B:388:LEU:HA | 1.91 | 0.43 |
| 1:B:411:ASP:HB3 | 1:B:413:HIS:CE1 | 2.54 | 0.43 |
| 1:B:597:SER:O | 1:B:623:ILE:HA | 2.19 | 0.43 |
| 1:C:142:TYR:O | 1:C:143:GLU:CB | 2.65 | 0.43 |
| 1:C:152:ILE:HG13 | 1:C:497:PHE:O | 2.19 | 0.43 |
| 1:C:170:ARG:CB | 1:C:473:ASP:OD1 | 2.67 | 0.43 |
| 1:D:184:TYR:O | 1:D:188:GLY:HA2 | 2.18 | 0.43 |
| 1:D:450:ARG:HD2 | 1:D:450:ARG:HA | 1.68 | 0.43 |

Continued on next page...

Not For Manuscript Review

*Continued from previous page...*

| Atom-1 | Atom-2 | Interatomic distance (Å) | Clash overlap (Å) |
|---|---|---|---|
| 1:F:148:LYS:N | 1:F:148:LYS:HD2 | 2.34 | 0.43 |
| 2:H:653:GLU:H | 2:H:653:GLU:CD | 2.21 | 0.43 |
| 2:J:180:LYS:HD3 | 2:J:180:LYS:HA | 1.90 | 0.43 |
| 2:K:259:SER:O | 2:K:263:ARG:HG2 | 2.19 | 0.43 |
| 2:K:376:ARG:HB2 | 2:K:433:LYS:H | 1.83 | 0.43 |
| 2:K:429:LEU:CB | 2:K:445:LYS:HD3 | 2.49 | 0.43 |
| 2:K:653:GLU:H | 2:K:653:GLU:CD | 2.21 | 0.43 |
| 1:A:170:ARG:CB | 1:A:473:ASP:OD1 | 2.67 | 0.42 |
| 1:B:152:ILE:HG13 | 1:B:497:PHE:O | 2.19 | 0.42 |
| 1:C:184:TYR:HD1 | 1:C:191:VAL:HG21 | 1.84 | 0.42 |
| 1:D:170:ARG:CB | 1:D:473:ASP:OD1 | 2.67 | 0.42 |
| 1:D:237:ARG:HD3 | 1:D:691:GLU:OE2 | 2.19 | 0.42 |
| 2:H:44:CYS:HG | 2:H:45:GLU:H | 1.66 | 0.42 |
| 2:H:315:VAL:HG22 | 2:H:315:VAL:O | 2.18 | 0.42 |
| 2:H:429:LEU:CB | 2:H:445:LYS:HD3 | 2.49 | 0.42 |
| 2:H:483:MET:SD | 2:H:569:PHE:HZ | 2.42 | 0.42 |
| 2:I:328:LEU:O | 2:I:332:THR:N | 2.40 | 0.42 |
| 2:J:429:LEU:O | 2:J:441:ILE:HD11 | 2.18 | 0.42 |
| 2:L:102:PHE:CE2 | 2:L:106:ILE:HD11 | 2.54 | 0.42 |
| 2:L:314:GLU:HB3 | 2:L:414:TRP:HD1 | 1.83 | 0.42 |
| 2:L:483:MET:SD | 2:L:569:PHE:HZ | 2.42 | 0.42 |
| 2:M:314:GLU:HB3 | 2:M:414:TRP:HD1 | 1.83 | 0.42 |
| 2:M:315:VAL:HG22 | 2:M:315:VAL:O | 2.18 | 0.42 |
| 1:A:152:ILE:HG13 | 1:A:497:PHE:O | 2.19 | 0.42 |
| 1:A:428:LEU:HB3 | 2:H:606:PHE:CE1 | 2.53 | 0.42 |
| 1:B:148:LYS:HD2 | 1:B:148:LYS:N | 2.34 | 0.42 |
| 1:B:184:TYR:O | 1:B:188:GLY:HA2 | 2.18 | 0.42 |
| 1:B:510:CYS:SG | 1:B:523:ILE:HG12 | 2.59 | 0.42 |
| 1:C:148:LYS:N | 1:C:148:LYS:HD2 | 2.34 | 0.42 |
| 1:C:510:CYS:SG | 1:C:523:ILE:HG12 | 2.59 | 0.42 |
| 1:E:148:LYS:N | 1:E:148:LYS:HD2 | 2.34 | 0.42 |
| 1:E:578:LEU:O | 1:E:582:VAL:HG23 | 2.18 | 0.42 |
| 2:H:102:PHE:CE2 | 2:H:106:ILE:HD11 | 2.54 | 0.42 |
| 2:H:240:CYS:HA | 2:H:251:GLU:HG3 | 2.01 | 0.42 |
| 2:H:259:SER:O | 2:H:263:ARG:HG2 | 2.19 | 0.42 |
| 2:H:376:ARG:HB2 | 2:H:433:LYS:H | 1.84 | 0.42 |
| 2:K:315:VAL:HG22 | 2:K:315:VAL:O | 2.18 | 0.42 |
| 2:M:221:GLU:HB3 | 2:M:286:GLU:HG3 | 1.99 | 0.42 |
| 2:M:240:CYS:HA | 2:M:251:GLU:HG3 | 2.01 | 0.42 |
| 2:M:483:MET:SD | 2:M:569:PHE:HZ | 2.42 | 0.42 |
| 1:A:237:ARG:HD3 | 1:A:691:GLU:OE2 | 2.20 | 0.42 |

*Continued on next page...*

Not For Manuscript Review

Continued from previous page...

| Atom-1 | Atom-2 | Interatomic distance (Å) | Clash overlap (Å) |
|---|---|---|---|
| 1:A:567:GLU:HG2 | 1:B:407:LEU:O | 2.18 | 0.42 |
| 1:B:142:TYR:O | 1:B:143:GLU:CB | 2.66 | 0.42 |
| 1:B:220:GLY:HA2 | 1:F:135:TYR:HE2 | 1.85 | 0.42 |
| 1:B:237:ARG:HD3 | 1:B:691:GLU:OE2 | 2.20 | 0.42 |
| 1:B:654:LYS:O | 1:B:658:LYS:HG2 | 2.19 | 0.42 |
| 1:C:184:TYR:O | 1:C:188:GLY:HA2 | 2.18 | 0.42 |
| 1:C:480:LYS:HE2 | 1:C:480:LYS:HB2 | 1.91 | 0.42 |
| 1:C:528:ILE:HG12 | 2:H:652:MET:CE | 2.49 | 0.42 |
| 1:C:654:LYS:O | 1:C:658:LYS:HG2 | 2.19 | 0.42 |
| 1:D:152:ILE:HG13 | 1:D:497:PHE:O | 2.19 | 0.42 |
| 1:E:237:ARG:HD3 | 1:E:691:GLU:OE2 | 2.20 | 0.42 |
| 1:F:152:ILE:HG13 | 1:F:497:PHE:O | 2.19 | 0.42 |
| 1:F:259:GLU:OE2 | 1:F:260:ARG:N | 2.52 | 0.42 |
| 1:F:597:SER:O | 1:F:623:ILE:HA | 2.19 | 0.42 |
| 2:H:62:ILE:HD12 | 2:H:71:TYR:H | 1.84 | 0.42 |
| 2:I:188:PHE:CD2 | 2:I:189:THR:HG23 | 2.54 | 0.42 |
| 2:J:188:PHE:CD2 | 2:J:189:THR:HG23 | 2.54 | 0.42 |
| 2:J:328:LEU:O | 2:J:332:THR:N | 2.40 | 0.42 |
| 2:J:429:LEU:CB | 2:J:445:LYS:HD3 | 2.49 | 0.42 |
| 2:K:44:CYS:HG | 2:K:45:GLU:H | 1.67 | 0.42 |
| 2:K:102:PHE:CE2 | 2:K:106:ILE:HD11 | 2.54 | 0.42 |
| 2:K:565:PHE:HB2 | 2:K:577:VAL:CG2 | 2.37 | 0.42 |
| 2:L:240:CYS:HA | 2:L:251:GLU:HG3 | 2.01 | 0.42 |
| 1:B:510:CYS:HB3 | 1:B:540:PHE:CD2 | 2.54 | 0.42 |
| 1:C:177:GLN:HG2 | 1:C:198:LYS:HZ2 | 1.85 | 0.42 |
| 1:C:237:ARG:HD3 | 1:C:691:GLU:OE2 | 2.20 | 0.42 |
| 1:C:259:GLU:OE2 | 1:C:260:ARG:N | 2.52 | 0.42 |
| 1:D:510:CYS:SG | 1:D:523:ILE:HG12 | 2.59 | 0.42 |
| 1:E:216:HIS:CG | 1:E:223:LEU:HD12 | 2.54 | 0.42 |
| 1:E:597:SER:O | 1:E:623:ILE:HA | 2.19 | 0.42 |
| 2:H:139:VAL:O | 2:H:141:VAL:HG23 | 2.19 | 0.42 |
| 2:I:429:LEU:HB2 | 2:I:445:LYS:HD3 | 2.00 | 0.42 |
| 2:K:139:VAL:O | 2:K:141:VAL:HG23 | 2.19 | 0.42 |
| 2:K:188:PHE:CD2 | 2:K:189:THR:HG23 | 2.54 | 0.42 |
| 2:K:240:CYS:HA | 2:K:251:GLU:HG3 | 2.01 | 0.42 |
| 2:L:221:GLU:HB3 | 2:L:286:GLU:HG3 | 2.00 | 0.42 |
| 2:M:102:PHE:CE2 | 2:M:106:ILE:HD11 | 2.54 | 0.42 |
| 2:M:650:MET:CE | 2:M:651:LYS:HZ3 | 2.27 | 0.42 |
| 1:A:148:LYS:N | 1:A:148:LYS:HD2 | 2.34 | 0.42 |
| 1:A:510:CYS:SG | 1:A:523:ILE:HG12 | 2.59 | 0.42 |
| 1:B:216:HIS:CG | 1:B:223:LEU:HD12 | 2.54 | 0.42 |

Continued on next page...

Not For Manuscript Review

*Continued from previous page...*

| Atom-1 | Atom-2 | Interatomic distance (Å) | Clash overlap (Å) |
|---|---|---|---|
| 1:D:143:GLU:C | 1:D:145:SER:N | 2.70 | 0.42 |
| 1:D:148:LYS:HD2 | 1:D:148:LYS:N | 2.34 | 0.42 |
| 1:E:152:ILE:HG13 | 1:E:497:PHE:O | 2.19 | 0.42 |
| 1:E:259:GLU:OE2 | 1:E:260:ARG:N | 2.52 | 0.42 |
| 1:F:237:ARG:HD3 | 1:F:691:GLU:OE2 | 2.20 | 0.42 |
| 2:H:309:HIS:O | 2:H:313:GLU:HG3 | 2.20 | 0.42 |
| 2:I:429:LEU:CB | 2:I:445:LYS:HD3 | 2.49 | 0.42 |
| 2:J:62:ILE:HD12 | 2:J:71:TYR:H | 1.84 | 0.42 |
| 2:J:429:LEU:HB2 | 2:J:445:LYS:HD3 | 2.01 | 0.42 |
| 2:J:477:THR:HG22 | 2:J:480:VAL:HG23 | 2.01 | 0.42 |
| 2:J:488:TRP:NE1 | 2:J:582:THR:O | 2.52 | 0.42 |
| 2:K:309:HIS:O | 2:K:313:GLU:HG3 | 2.20 | 0.42 |
| 2:K:446:ARG:HA | 2:K:446:ARG:HD2 | 1.84 | 0.42 |
| 2:L:650:MET:CE | 2:L:651:LYS:HZ3 | 2.27 | 0.42 |
| 1:A:143:GLU:C | 1:A:145:SER:N | 2.70 | 0.42 |
| 1:A:259:GLU:OE2 | 1:A:260:ARG:N | 2.52 | 0.42 |
| 1:A:450:ARG:HD2 | 1:A:450:ARG:HA | 1.68 | 0.42 |
| 1:B:259:GLU:OE2 | 1:B:260:ARG:N | 2.52 | 0.42 |
| 1:C:216:HIS:CG | 1:C:223:LEU:HD12 | 2.54 | 0.42 |
| 1:C:510:CYS:HB3 | 1:C:540:PHE:CD2 | 2.54 | 0.42 |
| 1:E:210:ASP:O | 1:E:214:LYS:HG3 | 2.20 | 0.42 |
| 1:E:411:ASP:HB3 | 1:E:413:HIS:CE1 | 2.54 | 0.42 |
| 1:E:567:GLU:HG2 | 1:F:407:LEU:O | 2.20 | 0.42 |
| 1:E:654:LYS:O | 1:E:658:LYS:HG2 | 2.19 | 0.42 |
| 1:F:210:ASP:O | 1:F:214:LYS:HG3 | 2.20 | 0.42 |
| 1:F:411:ASP:HB3 | 1:F:413:HIS:CE1 | 2.54 | 0.42 |
| 1:F:654:LYS:O | 1:F:658:LYS:HG2 | 2.19 | 0.42 |
| 2:H:11:ARG:HD2 | 2:H:11:ARG:HA | 1.86 | 0.42 |
| 2:I:477:THR:HG22 | 2:I:480:VAL:HG23 | 2.01 | 0.42 |
| 2:I:488:TRP:NE1 | 2:I:582:THR:O | 2.52 | 0.42 |
| 2:L:188:PHE:CD2 | 2:L:189:THR:HG23 | 2.54 | 0.42 |
| 2:M:188:PHE:CD2 | 2:M:189:THR:HG23 | 2.54 | 0.42 |
| 2:M:244:ARG:NH1 | 2:M:245:ARG:HB2 | 2.35 | 0.42 |
| 1:B:480:LYS:HE2 | 1:B:480:LYS:HB2 | 1.91 | 0.42 |
| 1:B:547:ARG:NH1 | 1:D:354:SER:OG | 2.53 | 0.42 |
| 1:F:216:HIS:CG | 1:F:223:LEU:HD12 | 2.54 | 0.42 |
| 2:H:188:PHE:CD2 | 2:H:189:THR:HG23 | 2.54 | 0.42 |
| 2:I:653:GLU:H | 2:I:653:GLU:CD | 2.21 | 0.42 |
| 2:J:116:PRO:HB3 | 2:J:279:TYR:CE1 | 2.55 | 0.42 |
| 2:J:652:MET:HG3 | 2:J:654:HIS:HD2 | 1.83 | 0.42 |
| 2:L:116:PRO:HB3 | 2:L:279:TYR:CE1 | 2.55 | 0.42 |

*Continued on next page...*

Not For Manuscript Review

*Continued from previous page...*

| Atom-1 | Atom-2 | Interatomic distance (Å) | Clash overlap (Å) |
|---|---|---|---|
| 2:L:244:ARG:NH1 | 2:L:245:ARG:HB2 | 2.35 | 0.42 |
| 1:C:220:GLY:HA2 | 1:E:135:TYR:HE2 | 1.85 | 0.42 |
| 1:C:450:ARG:HA | 1:C:450:ARG:HD2 | 1.68 | 0.42 |
| 2:I:261:GLU:O | 2:I:265:ARG:HG3 | 2.20 | 0.42 |
| 2:J:653:GLU:H | 2:J:653:GLU:CD | 2.20 | 0.42 |
| 2:L:413:SER:CB | 2:L:425:LEU:HG | 2.50 | 0.42 |
| 2:M:116:PRO:HB3 | 2:M:279:TYR:CE1 | 2.55 | 0.42 |
| 1:A:210:ASP:O | 1:A:214:LYS:HG3 | 2.20 | 0.42 |
| 1:A:597:SER:O | 1:A:623:ILE:HA | 2.19 | 0.42 |
| 1:D:337:HIS:HB3 | 1:D:338:PHE:H | 1.70 | 0.42 |
| 2:H:244:ARG:NH1 | 2:H:245:ARG:HB2 | 2.35 | 0.42 |
| 2:I:62:ILE:HD12 | 2:I:71:TYR:H | 1.84 | 0.42 |
| 2:I:116:PRO:HB3 | 2:I:279:TYR:CE1 | 2.55 | 0.42 |
| 2:I:652:MET:HG3 | 2:I:654:HIS:HD2 | 1.83 | 0.42 |
| 2:J:102:PHE:CE2 | 2:J:106:ILE:HD11 | 2.54 | 0.42 |
| 2:J:165:PRO:HD2 | 2:J:212:ARG:HG2 | 2.01 | 0.42 |
| 2:K:224:ILE:HD12 | 2:K:226:PHE:HE1 | 1.85 | 0.42 |
| 2:K:244:ARG:NH1 | 2:K:245:ARG:HB2 | 2.35 | 0.42 |
| 2:M:62:ILE:HD12 | 2:M:71:TYR:H | 1.84 | 0.42 |
| 1:D:597:SER:O | 1:D:623:ILE:HA | 2.19 | 0.42 |
| 2:I:102:PHE:CE2 | 2:I:106:ILE:HD11 | 2.54 | 0.42 |
| 2:I:224:ILE:HD12 | 2:I:226:PHE:HE1 | 1.85 | 0.42 |
| 2:I:244:ARG:NH1 | 2:I:245:ARG:HB2 | 2.35 | 0.42 |
| 2:J:224:ILE:HD12 | 2:J:226:PHE:HE1 | 1.85 | 0.42 |
| 2:J:244:ARG:NH1 | 2:J:245:ARG:HB2 | 2.35 | 0.42 |
| 2:J:261:GLU:O | 2:J:265:ARG:HG3 | 2.20 | 0.42 |
| 2:K:11:ARG:HD2 | 2:K:11:ARG:HA | 1.86 | 0.42 |
| 2:L:279:TYR:OH | 2:L:282:ALA:O | 2.35 | 0.42 |
| 1:A:411:ASP:HB3 | 1:A:413:HIS:CE1 | 2.54 | 0.41 |
| 1:B:450:ARG:HA | 1:B:450:ARG:HD2 | 1.68 | 0.41 |
| 1:D:210:ASP:O | 1:D:214:LYS:HG3 | 2.20 | 0.41 |
| 2:H:224:ILE:HD12 | 2:H:226:PHE:HE1 | 1.85 | 0.41 |
| 2:H:561:ILE:O | 2:H:580:VAL:HA | 2.20 | 0.41 |
| 2:I:139:VAL:O | 2:I:141:VAL:HG23 | 2.19 | 0.41 |
| 2:I:165:PRO:HD2 | 2:I:212:ARG:HG2 | 2.01 | 0.41 |
| 2:J:254:PRO:HG3 | 2:J:315:VAL:CG2 | 2.47 | 0.41 |
| 2:K:561:ILE:O | 2:K:580:VAL:HA | 2.20 | 0.41 |
| 2:L:544:PHE:HB3 | 2:L:569:PHE:CD2 | 2.55 | 0.41 |
| 2:L:653:GLU:H | 2:L:653:GLU:CD | 2.21 | 0.41 |
| 2:M:413:SER:CB | 2:M:425:LEU:HG | 2.50 | 0.41 |
| 1:A:135:TYR:HE2 | 1:E:220:GLY:HA2 | 1.85 | 0.41 |

*Continued on next page...*

Not For Manuscript Review

*Continued from previous page...*

| Atom-1 | Atom-2 | Interatomic distance (Å) | Clash overlap (Å) |
|---|---|---|---|
| 1:B:210:ASP:O | 1:B:214:LYS:HG3 | 2.20 | 0.41 |
| 1:C:210:ASP:O | 1:C:214:LYS:HG3 | 2.20 | 0.41 |
| 1:D:142:TYR:CA | 1:D:145:SER:H | 2.33 | 0.41 |
| 1:F:558:THR:HA | 1:F:601:GLY:HA3 | 2.02 | 0.41 |
| 2:I:556:ARG:HB3 | 2:I:561:ILE:HD11 | 2.02 | 0.41 |
| 2:J:309:HIS:O | 2:J:313:GLU:HG3 | 2.20 | 0.41 |
| 2:M:279:TYR:OH | 2:M:282:ALA:O | 2.35 | 0.41 |
| 2:M:544:PHE:HB3 | 2:M:569:PHE:CD2 | 2.55 | 0.41 |
| 1:A:266:ILE:O | 1:A:299:LYS:NZ | 2.53 | 0.41 |
| 1:D:266:ILE:O | 1:D:299:LYS:NZ | 2.53 | 0.41 |
| 1:D:411:ASP:HB3 | 1:D:413:HIS:CE1 | 2.54 | 0.41 |
| 1:E:266:ILE:O | 1:E:299:LYS:NZ | 2.53 | 0.41 |
| 2:H:261:GLU:O | 2:H:265:ARG:HG3 | 2.20 | 0.41 |
| 2:I:309:HIS:O | 2:I:313:GLU:HG3 | 2.20 | 0.41 |
| 2:J:139:VAL:O | 2:J:141:VAL:HG23 | 2.19 | 0.41 |
| 2:J:556:ARG:HB3 | 2:J:561:ILE:HD11 | 2.03 | 0.41 |
| 2:K:574:HIS:CD2 | 2:M:48:ARG:HH21 | 2.38 | 0.41 |
| 2:L:62:ILE:HD12 | 2:L:71:TYR:H | 1.84 | 0.41 |
| 2:L:626:LYS:HA | 2:L:626:LYS:HD2 | 1.93 | 0.41 |
| 2:M:429:LEU:HB2 | 2:M:445:LYS:HD3 | 2.00 | 0.41 |
| 2:M:653:GLU:H | 2:M:653:GLU:CD | 2.21 | 0.41 |
| 1:A:142:TYR:CA | 1:A:145:SER:H | 2.34 | 0.41 |
| 1:A:337:HIS:HB3 | 1:A:338:PHE:H | 1.70 | 0.41 |
| 1:E:558:THR:HA | 1:E:601:GLY:HA3 | 2.02 | 0.41 |
| 1:F:266:ILE:O | 1:F:299:LYS:NZ | 2.53 | 0.41 |
| 2:I:254:PRO:HG3 | 2:I:315:VAL:CG2 | 2.47 | 0.41 |
| 2:J:561:ILE:O | 2:J:580:VAL:HA | 2.20 | 0.41 |
| 1:C:558:THR:HA | 1:C:601:GLY:HA3 | 2.02 | 0.41 |
| 2:H:565:PHE:HB2 | 2:H:577:VAL:CG2 | 2.37 | 0.41 |
| 2:I:561:ILE:O | 2:I:580:VAL:HA | 2.20 | 0.41 |
| 2:J:544:PHE:HB3 | 2:J:569:PHE:CD2 | 2.55 | 0.41 |
| 2:K:630:LEU:HD22 | 2:K:645:VAL:HG12 | 2.02 | 0.41 |
| 2:L:169:LYS:HB3 | 2:L:170:ALA:H | 1.52 | 0.41 |
| 1:A:347:GLN:HG3 | 1:A:351:LYS:HE2 | 2.02 | 0.41 |
| 1:B:558:THR:HA | 1:B:601:GLY:HA3 | 2.02 | 0.41 |
| 1:B:616:PHE:HD1 | 1:B:678:GLY:HA3 | 1.86 | 0.41 |
| 1:C:407:LEU:O | 1:D:567:GLU:HG2 | 2.20 | 0.41 |
| 1:C:616:PHE:HD1 | 1:C:678:GLY:HA3 | 1.86 | 0.41 |
| 1:E:235:GLY:HA3 | 2:L:494:ASP:OD1 | 2.21 | 0.41 |
| 2:H:116:PRO:HB3 | 2:H:279:TYR:CE1 | 2.55 | 0.41 |
| 2:I:565:PHE:HB3 | 2:I:567:PHE:CE1 | 2.55 | 0.41 |

*Continued on next page...*

Not For Manuscript Review

Continued from previous page...

| Atom-1 | Atom-2 | Interatomic distance (Å) | Clash overlap (Å) |
|---|---|---|---|
| 2:I:573:MET:HE3 | 2:K:58:GLU:OE1 | 2.14 | 0.41 |
| 2:K:116:PRO:HB3 | 2:K:279:TYR:CE1 | 2.55 | 0.41 |
| 2:K:488:TRP:NE1 | 2:K:582:THR:O | 2.52 | 0.41 |
| 2:K:544:PHE:HB3 | 2:K:569:PHE:CD2 | 2.55 | 0.41 |
| 2:K:556:ARG:HB3 | 2:K:561:ILE:HD11 | 2.02 | 0.41 |
| 2:L:127:LYS:O | 2:L:131:LYS:HG2 | 2.21 | 0.41 |
| 2:L:429:LEU:HB2 | 2:L:445:LYS:HD3 | 2.00 | 0.41 |
| 2:L:561:ILE:O | 2:L:580:VAL:HA | 2.20 | 0.41 |
| 2:L:562:VAL:CG1 | 2:L:580:VAL:HG22 | 2.44 | 0.41 |
| 2:M:224:ILE:HD12 | 2:M:226:PHE:HE1 | 1.85 | 0.41 |
| 2:M:561:ILE:O | 2:M:580:VAL:HA | 2.20 | 0.41 |
| 1:E:616:PHE:HD1 | 1:E:678:GLY:HA3 | 1.86 | 0.41 |
| 2:H:488:TRP:NE1 | 2:H:582:THR:O | 2.52 | 0.41 |
| 2:H:556:ARG:HB3 | 2:H:561:ILE:HD11 | 2.03 | 0.41 |
| 2:I:544:PHE:HB3 | 2:I:569:PHE:CD2 | 2.55 | 0.41 |
| 2:K:261:GLU:O | 2:K:265:ARG:HG3 | 2.20 | 0.41 |
| 2:K:376:ARG:CG | 2:K:433:LYS:HB2 | 2.51 | 0.41 |
| 2:L:191:MET:H | 2:L:191:MET:HG3 | 1.73 | 0.41 |
| 2:L:224:ILE:HD12 | 2:L:226:PHE:HE1 | 1.85 | 0.41 |
| 2:M:556:ARG:HB3 | 2:M:561:ILE:HD11 | 2.02 | 0.41 |
| 2:M:626:LYS:HA | 2:M:626:LYS:HD2 | 1.93 | 0.41 |
| 1:B:144:ARG:HG2 | 2:K:606:PHE:CZ | 2.56 | 0.41 |
| 1:D:137:HIS:HA | 2:M:602:LEU:HD11 | 2.03 | 0.41 |
| 1:D:347:GLN:HG3 | 1:D:351:LYS:HE2 | 2.02 | 0.41 |
| 1:F:616:PHE:HD1 | 1:F:678:GLY:HA3 | 1.86 | 0.41 |
| 2:H:376:ARG:CG | 2:H:433:LYS:HB2 | 2.51 | 0.41 |
| 2:H:630:LEU:HD22 | 2:H:645:VAL:HG12 | 2.03 | 0.41 |
| 2:I:144:GLY:O | 2:I:209:ILE:HD12 | 2.21 | 0.41 |
| 2:I:413:SER:CB | 2:I:425:LEU:HG | 2.50 | 0.41 |
| 2:J:144:GLY:O | 2:J:209:ILE:HD12 | 2.21 | 0.41 |
| 2:K:413:SER:CB | 2:K:425:LEU:HG | 2.50 | 0.41 |
| 2:L:144:GLY:O | 2:L:209:ILE:HD12 | 2.21 | 0.41 |
| 2:M:127:LYS:O | 2:M:131:LYS:HG2 | 2.21 | 0.41 |
| 1:B:177:GLN:CG | 1:B:198:LYS:HZ2 | 2.34 | 0.41 |
| 1:B:337:HIS:HB3 | 1:B:338:PHE:H | 1.70 | 0.41 |
| 1:C:266:ILE:O | 1:C:299:LYS:NZ | 2.53 | 0.41 |
| 1:C:347:GLN:HG3 | 1:C:351:LYS:HE2 | 2.02 | 0.41 |
| 1:C:479:VAL:HB | 2:H:653:GLU:OE1 | 2.21 | 0.41 |
| 1:C:567:GLU:HG2 | 1:D:407:LEU:O | 2.20 | 0.41 |
| 1:F:347:GLN:HG3 | 1:F:351:LYS:HE2 | 2.02 | 0.41 |
| 1:F:479:VAL:HB | 2:I:653:GLU:OE1 | 2.20 | 0.41 |

Continued on next page...

Not For Manuscript Review

*Continued from previous page...*

| Atom-1 | Atom-2 | Interatomic distance (Å) | Clash overlap (Å) |
|---|---|---|---|
| 2:H:413:SER:CB | 2:H:425:LEU:HG | 2.50 | 0.41 |
| 2:H:544:PHE:HB3 | 2:H:569:PHE:CD2 | 2.55 | 0.41 |
| 2:H:565:PHE:HB3 | 2:H:567:PHE:CE1 | 2.55 | 0.41 |
| 2:J:413:SER:CB | 2:J:425:LEU:HG | 2.50 | 0.41 |
| 2:J:565:PHE:HB3 | 2:J:567:PHE:CE1 | 2.55 | 0.41 |
| 2:K:565:PHE:HB3 | 2:K:567:PHE:CE1 | 2.55 | 0.41 |
| 2:K:607:GLY:O | 2:K:609:SER:N | 2.51 | 0.41 |
| 2:L:261:GLU:O | 2:L:265:ARG:HG3 | 2.20 | 0.41 |
| 2:L:556:ARG:HB3 | 2:L:561:ILE:HD11 | 2.03 | 0.41 |
| 2:L:565:PHE:HB3 | 2:L:567:PHE:CE1 | 2.55 | 0.41 |
| 2:M:144:GLY:O | 2:M:209:ILE:HD12 | 2.21 | 0.41 |
| 2:M:169:LYS:HB3 | 2:M:170:ALA:H | 1.52 | 0.41 |
| 2:M:261:GLU:O | 2:M:265:ARG:HG3 | 2.20 | 0.41 |
| 2:M:309:HIS:O | 2:M:313:GLU:HG3 | 2.20 | 0.41 |
| 2:M:565:PHE:HB3 | 2:M:567:PHE:CE1 | 2.55 | 0.41 |
| 1:A:378:ILE:HG12 | 1:A:422:HIS:HB2 | 2.03 | 0.41 |
| 1:A:391:PRO:O | 1:A:394:VAL:HG22 | 2.21 | 0.41 |
| 1:B:266:ILE:O | 1:B:299:LYS:NZ | 2.53 | 0.41 |
| 1:B:347:GLN:HG3 | 1:B:351:LYS:HE2 | 2.02 | 0.41 |
| 1:B:362:SER:HA | 1:B:385:THR:O | 2.21 | 0.41 |
| 1:B:461:LEU:HD12 | 1:B:461:LEU:O | 2.21 | 0.41 |
| 1:C:362:SER:HA | 1:C:385:THR:O | 2.21 | 0.41 |
| 1:D:378:ILE:HG12 | 1:D:422:HIS:HB2 | 2.03 | 0.41 |
| 1:D:391:PRO:O | 1:D:394:VAL:HG22 | 2.21 | 0.41 |
| 1:E:347:GLN:HG3 | 1:E:351:LYS:HE2 | 2.02 | 0.41 |
| 1:F:161:ASP:HB3 | 1:F:162:PRO:CD | 2.51 | 0.41 |
| 1:F:176:GLU:HB3 | 1:F:198:LYS:HZ2 | 1.84 | 0.41 |
| 1:F:428:LEU:HB3 | 2:M:606:PHE:CE1 | 2.51 | 0.41 |
| 2:H:124:MET:HE3 | 2:H:124:MET:HB3 | 1.85 | 0.41 |
| 2:L:309:HIS:O | 2:L:313:GLU:HG3 | 2.20 | 0.41 |
| 2:M:562:VAL:CG1 | 2:M:580:VAL:HG22 | 2.44 | 0.41 |
| 1:A:144:ARG:HG2 | 2:L:606:PHE:CE2 | 2.56 | 0.40 |
| 1:A:576:ALA:HB2 | 1:B:372:ALA:CB | 2.52 | 0.40 |
| 1:B:335:GLU:O | 1:B:337:HIS:ND1 | 2.55 | 0.40 |
| 1:B:378:ILE:HG12 | 1:B:422:HIS:HB2 | 2.03 | 0.40 |
| 1:C:335:GLU:O | 1:C:337:HIS:ND1 | 2.54 | 0.40 |
| 1:D:259:GLU:OE2 | 1:D:260:ARG:N | 2.52 | 0.40 |
| 1:E:142:TYR:CA | 1:E:145:SER:H | 2.33 | 0.40 |
| 1:E:161:ASP:HB3 | 1:E:162:PRO:CD | 2.52 | 0.40 |
| 1:F:142:TYR:CA | 1:F:145:SER:H | 2.33 | 0.40 |
| 1:F:429:HIS:NE2 | 2:M:607:GLY:HA2 | 2.36 | 0.40 |

*Continued on next page...*

Not For Manuscript Review

Continued from previous page...

| Atom-1 | Atom-2 | Interatomic distance (Å) | Clash overlap (Å) |
|---|---|---|---|
| 2:H:127:LYS:O | 2:H:131:LYS:HG2 | 2.21 | 0.40 |
| 2:H:144:GLY:O | 2:H:209:ILE:HD12 | 2.21 | 0.40 |
| 2:H:607:GLY:O | 2:H:609:SER:N | 2.51 | 0.40 |
| 2:J:127:LYS:O | 2:J:131:LYS:HG2 | 2.21 | 0.40 |
| 2:J:607:GLY:O | 2:J:609:SER:N | 2.51 | 0.40 |
| 2:K:64:PRO:HA | 2:K:65:PRO:HD3 | 1.96 | 0.40 |
| 2:K:75:GLU:H | 2:K:75:GLU:CD | 2.25 | 0.40 |
| 2:K:141:VAL:HG22 | 2:K:298:PHE:HB3 | 2.03 | 0.40 |
| 2:K:144:GLY:O | 2:K:209:ILE:HD12 | 2.21 | 0.40 |
| 2:K:373:THR:OG1 | 2:K:435:ALA:O | 2.32 | 0.40 |
| 1:A:142:TYR:HA | 1:A:145:SER:CB | 2.44 | 0.40 |
| 1:C:378:ILE:HG12 | 1:C:422:HIS:HB2 | 2.03 | 0.40 |
| 1:C:428:LEU:HB3 | 2:J:606:PHE:CE1 | 2.52 | 0.40 |
| 1:C:461:LEU:HD12 | 1:C:461:LEU:O | 2.22 | 0.40 |
| 1:C:670:SER:O | 1:C:672:ALA:N | 2.51 | 0.40 |
| 1:E:407:LEU:O | 1:F:567:GLU:HG2 | 2.22 | 0.40 |
| 2:M:180:LYS:HD3 | 2:M:180:LYS:HA | 1.89 | 0.40 |
| 2:M:191:MET:H | 2:M:191:MET:HG3 | 1.73 | 0.40 |
| 1:A:171:MET:HG2 | 1:A:468:GLY:HA3 | 2.03 | 0.40 |
| 1:A:397:ALA:HB2 | 2:K:650:MET:HG2 | 2.03 | 0.40 |
| 1:A:558:THR:HA | 1:A:601:GLY:HA3 | 2.02 | 0.40 |
| 1:B:171:MET:HG2 | 1:B:468:GLY:HA3 | 2.03 | 0.40 |
| 1:B:588:PRO:HB3 | 1:B:613:ARG:NH1 | 2.37 | 0.40 |
| 1:B:670:SER:O | 1:B:672:ALA:N | 2.51 | 0.40 |
| 1:C:171:MET:HG2 | 1:C:468:GLY:HA3 | 2.03 | 0.40 |
| 1:C:474:MET:SD | 1:C:620:ASN:HB2 | 2.62 | 0.40 |
| 1:D:142:TYR:HA | 1:D:145:SER:CB | 2.44 | 0.40 |
| 1:D:171:MET:HG2 | 1:D:468:GLY:HA3 | 2.03 | 0.40 |
| 1:D:558:THR:HA | 1:D:601:GLY:HA3 | 2.02 | 0.40 |
| 1:D:670:SER:O | 1:D:672:ALA:N | 2.51 | 0.40 |
| 1:F:391:PRO:O | 1:F:394:VAL:HG22 | 2.21 | 0.40 |
| 2:H:75:GLU:H | 2:H:75:GLU:CD | 2.25 | 0.40 |
| 2:H:141:VAL:HG22 | 2:H:298:PHE:HB3 | 2.04 | 0.40 |
| 2:I:127:LYS:O | 2:I:131:LYS:HG2 | 2.21 | 0.40 |
| 2:I:225:PHE:CZ | 2:I:332:THR:OG1 | 2.75 | 0.40 |
| 2:I:376:ARG:CG | 2:I:433:LYS:HB2 | 2.51 | 0.40 |
| 2:J:228:LYS:HE2 | 2:J:228:LYS:HB2 | 1.97 | 0.40 |
| 2:J:376:ARG:CG | 2:J:433:LYS:HB2 | 2.51 | 0.40 |
| 2:K:127:LYS:O | 2:K:131:LYS:HG2 | 2.21 | 0.40 |
| 1:A:177:GLN:HG2 | 1:A:198:LYS:HZ2 | 1.87 | 0.40 |
| 1:A:474:MET:SD | 1:A:620:ASN:HB2 | 2.62 | 0.40 |

Continued on next page...

Not For Manuscript Review

*Continued from previous page...*

| Atom-1 | Atom-2 | Interatomic distance (Å) | Clash overlap (Å) |
|---|---|---|---|
| 1:A:616:PHE:HD1 | 1:A:678:GLY:HA3 | 1.86 | 0.40 |
| 1:B:354:SER:OG | 1:F:547:ARG:NH1 | 2.55 | 0.40 |
| 1:B:474:MET:SD | 1:B:620:ASN:HB2 | 2.62 | 0.40 |
| 1:C:337:HIS:HB3 | 1:C:338:PHE:H | 1.70 | 0.40 |
| 1:C:588:PRO:HB3 | 1:C:613:ARG:NH1 | 2.37 | 0.40 |
| 1:E:391:PRO:O | 1:E:394:VAL:HG22 | 2.21 | 0.40 |
| 2:H:64:PRO:HA | 2:H:65:PRO:HD3 | 1.96 | 0.40 |
| 2:I:227:ASP:HA | 2:I:332:THR:HG23 | 2.03 | 0.40 |
| 2:I:607:GLY:O | 2:I:609:SER:N | 2.51 | 0.40 |
| 2:I:619:ILE:HD12 | 2:I:645:VAL:HG22 | 2.04 | 0.40 |
| 2:J:141:VAL:HG22 | 2:J:298:PHE:HB3 | 2.03 | 0.40 |
| 2:J:227:ASP:HA | 2:J:332:THR:HG23 | 2.03 | 0.40 |
| 2:J:279:TYR:OH | 2:J:282:ALA:O | 2.35 | 0.40 |
| 2:L:630:LEU:HD22 | 2:L:645:VAL:HG12 | 2.03 | 0.40 |
| 2:M:75:GLU:H | 2:M:75:GLU:CD | 2.25 | 0.40 |
| 1:A:295:ILE:HD11 | 1:A:674:LEU:HD13 | 2.03 | 0.40 |
| 1:A:670:SER:O | 1:A:672:ALA:N | 2.51 | 0.40 |
| 1:B:391:PRO:O | 1:B:394:VAL:HG22 | 2.21 | 0.40 |
| 1:B:593:LEU:HD23 | 1:B:593:LEU:HA | 1.96 | 0.40 |
| 1:C:295:ILE:HD11 | 1:C:674:LEU:HD13 | 2.03 | 0.40 |
| 1:D:616:PHE:HD1 | 1:D:678:GLY:HA3 | 1.86 | 0.40 |
| 1:F:335:GLU:O | 1:F:337:HIS:ND1 | 2.54 | 0.40 |
| 1:F:461:LEU:O | 1:F:461:LEU:HD12 | 2.22 | 0.40 |
| 2:H:262:THR:HG23 | 2:H:265:ARG:HH21 | 1.86 | 0.40 |
| 2:H:478:PRO:HB3 | 2:H:537:VAL:HG13 | 2.03 | 0.40 |
| 2:I:141:VAL:HG22 | 2:I:298:PHE:HB3 | 2.04 | 0.40 |
| 2:I:630:LEU:HD22 | 2:I:645:VAL:HG12 | 2.03 | 0.40 |
| 2:J:225:PHE:CZ | 2:J:332:THR:OG1 | 2.75 | 0.40 |
| 2:L:75:GLU:H | 2:L:75:GLU:CD | 2.25 | 0.40 |
| 2:L:376:ARG:CG | 2:L:433:LYS:HB2 | 2.51 | 0.40 |
| 2:M:506:THR:O | 2:M:524:HIS:HB2 | 2.22 | 0.40 |

There are no symmetry-related clashes.

#### 4.3 Torsion angles (i)

##### 4.3.1 Protein backbone (i)

In the following table, the Percentiles column shows the percent Ramachandran outliers of the chain as a percentile score with respect to all PDB entries followed by that with respect to all EM entries.

Not For Manuscript Review

The Analysed column shows the number of residues for which the backbone conformation was analysed, and the total number of residues.

| Mol | Chain | Analysed | Favoured | Allowed | Outliers | Percentiles | |
|---|---|---|---|---|---|---|---|
| 1 | A | 564/566 (100%) | 506 (90%) | 58 (10%) | 0 | 100 | 100 |
| 1 | B | 564/566 (100%) | 507 (90%) | 57 (10%) | 0 | 100 | 100 |
| 1 | C | 564/566 (100%) | 507 (90%) | 57 (10%) | 0 | 100 | 100 |
| 1 | D | 564/566 (100%) | 507 (90%) | 57 (10%) | 0 | 100 | 100 |
| 1 | E | 564/566 (100%) | 507 (90%) | 57 (10%) | 0 | 100 | 100 |
| 1 | F | 564/566 (100%) | 507 (90%) | 57 (10%) | 0 | 100 | 100 |
| 2 | H | 674/678 (99%) | 616 (91%) | 58 (9%) | 0 | 100 | 100 |
| 2 | I | 674/678 (99%) | 615 (91%) | 59 (9%) | 0 | 100 | 100 |
| 2 | J | 674/678 (99%) | 615 (91%) | 59 (9%) | 0 | 100 | 100 |
| 2 | K | 674/678 (99%) | 615 (91%) | 59 (9%) | 0 | 100 | 100 |
| 2 | L | 674/678 (99%) | 615 (91%) | 59 (9%) | 0 | 100 | 100 |
| 2 | M | 674/678 (99%) | 614 (91%) | 60 (9%) | 0 | 100 | 100 |
| All | All | 7428/7464 (100%) | 6731 (91%) | 697 (9%) | 0 | 100 | 100 |

There are no Ramachandran outliers to report.

##### 4.3.2 Protein sidechains (i)

In the following table, the Percentiles column shows the percent sidechain outliers of the chain as a percentile score with respect to all PDB entries followed by that with respect to all EM entries.

The Analysed column shows the number of residues for which the sidechain conformation was analysed, and the total number of residues.

| Mol | Chain | Analysed | Rotameric | Outliers | Percentiles | |
|---|---|---|---|---|---|---|
| 1 | A | 455/455 (100%) | 455 (100%) | 0 | 100 | 100 |
| 1 | B | 455/455 (100%) | 455 (100%) | 0 | 100 | 100 |
| 1 | C | 455/455 (100%) | 455 (100%) | 0 | 100 | 100 |
| 1 | D | 455/455 (100%) | 455 (100%) | 0 | 100 | 100 |
| 1 | E | 455/455 (100%) | 455 (100%) | 0 | 100 | 100 |
| 1 | F | 455/455 (100%) | 455 (100%) | 0 | 100 | 100 |
| 2 | H | 565/565 (100%) | 565 (100%) | 0 | 100 | 100 |

*Continued on next page...*

Not For Manuscript Review

*Continued from previous page...*

| Mol | Chain | Analysed | Rotameric | Outliers | Percentiles | |
|---|---|---|---|---|---|---|
| 2 | I | 565/565 (100%) | 565 (100%) | 0 | 100 | 100 |
| 2 | J | 565/565 (100%) | 565 (100%) | 0 | 100 | 100 |
| 2 | K | 565/565 (100%) | 565 (100%) | 0 | 100 | 100 |
| 2 | L | 565/565 (100%) | 565 (100%) | 0 | 100 | 100 |
| 2 | M | 565/565 (100%) | 565 (100%) | 0 | 100 | 100 |
| All | All | 6120/6120 (100%) | 6120 (100%) | 0 | 100 | 100 |

There are no protein residues with a non-rotameric sidechain to report.

Sometimes sidechains can be flipped to improve hydrogen bonding and reduce clashes. All (43) such sidechains are listed below:

| Mol | Chain | Res | Type |
|---|---|---|---|
| 1 | A | 177 | GLN |
| 1 | A | 186 | GLN |
| 1 | A | 413 | HIS |
| 1 | A | 548 | ASN |
| 1 | B | 177 | GLN |
| 1 | B | 186 | GLN |
| 1 | B | 413 | HIS |
| 1 | B | 548 | ASN |
| 1 | C | 177 | GLN |
| 1 | C | 186 | GLN |
| 1 | C | 413 | HIS |
| 1 | C | 548 | ASN |
| 1 | D | 177 | GLN |
| 1 | D | 186 | GLN |
| 1 | D | 548 | ASN |
| 1 | E | 177 | GLN |
| 1 | E | 186 | GLN |
| 1 | E | 548 | ASN |
| 1 | F | 177 | GLN |
| 1 | F | 186 | GLN |
| 1 | F | 548 | ASN |
| 2 | H | 36 | HIS |
| 2 | H | 466 | HIS |
| 2 | H | 495 | ASN |
| 2 | H | 654 | HIS |
| 2 | I | 36 | HIS |
| 2 | I | 466 | HIS |
| 2 | I | 495 | ASN |

*Continued on next page...*

Not For Manuscript Review

*Continued from previous page...*

| Mol | Chain | Res | Type |
|---|---|---|---|
| 2 | I | 654 | HIS |
| 2 | J | 36 | HIS |
| 2 | J | 466 | HIS |
| 2 | J | 495 | ASN |
| 2 | J | 654 | HIS |
| 2 | K | 36 | HIS |
| 2 | K | 466 | HIS |
| 2 | K | 495 | ASN |
| 2 | K | 654 | HIS |
| 2 | L | 36 | HIS |
| 2 | L | 466 | HIS |
| 2 | L | 495 | ASN |
| 2 | M | 36 | HIS |
| 2 | M | 466 | HIS |
| 2 | M | 495 | ASN |

##### 4.3.3 RNA (i)

There are no RNA molecules in this entry.

#### 4.4 Non-standard residues in protein, DNA, RNA chains (i)

There are no non-standard protein/DNA/RNA residues in this entry.

#### 4.5 Carbohydrates (i)

There are no monosaccharides in this entry.

#### 4.6 Ligand geometry (i)

6 ligands are modelled in this entry.

In the following table, the Counts columns list the number of bonds (or angles) for which Mogul statistics could be retrieved, the number of bonds (or angles) that are observed in the model and the number of bonds (or angles) that are defined in the Chemical Component Dictionary. The Link column lists molecule types, if any, to which the group is linked. The Z score for a bond length (or angle) is the number of standard deviations the observed value is removed from the expected value. A bond length (or angle) with $|Z| > 2$ is considered an outlier worth inspection. RMSZ is the root-mean-square of all Z scores of the bond lengths (or angles).

Not For Manuscript Review

| Mol | Type | Chain | Res | Link | Bond lengths | | | Bond angles | | |
|---|---|---|---|---|---|---|---|---|---|---|
| | | | | | Counts | RMSZ | #\|Z\| > 2 | Counts | RMSZ | #\|Z\| > 2 |
| 3 | BTI | G | 804 | - | 16,16,16 | 1.27 | 1 (6%) | 21,21,21 | 1.52 | 3 (14%) |
| 3 | BTI | G | 802 | - | 16,16,16 | 1.22 | 1 (6%) | 21,21,21 | 1.43 | 4 (19%) |
| 3 | BTI | G | 803 | - | 16,16,16 | 1.25 | 1 (6%) | 21,21,21 | 1.55 | 5 (23%) |
| 3 | BTI | G | 801 | - | 16,16,16 | 1.24 | 1 (6%) | 21,21,21 | 1.54 | 5 (23%) |
| 3 | BTI | G | 806 | - | 16,16,16 | 1.23 | 1 (6%) | 21,21,21 | 1.40 | 4 (19%) |
| 3 | BTI | G | 805 | - | 16,16,16 | 1.25 | 1 (6%) | 21,21,21 | 1.54 | 5 (23%) |

In the following table, the Chirals column lists the number of chiral outliers, the number of chiral centers analysed, the number of these observed in the model and the number defined in the Chemical Component Dictionary. Similar counts are reported in the Torsion and Rings columns. '-' means no outliers of that kind were identified.

| Mol | Type | Chain | Res | Link | Chirals | Torsions | Rings |
|---|---|---|---|---|---|---|---|
| 3 | BTI | G | 804 | - | - | 0/5/27/27 | 0/2/2/2 |
| 3 | BTI | G | 802 | - | - | 0/5/27/27 | 0/2/2/2 |
| 3 | BTI | G | 803 | - | - | 0/5/27/27 | 0/2/2/2 |
| 3 | BTI | G | 801 | - | - | 0/5/27/27 | 0/2/2/2 |
| 3 | BTI | G | 806 | - | - | 0/5/27/27 | 0/2/2/2 |
| 3 | BTI | G | 805 | - | - | 0/5/27/27 | 0/2/2/2 |

All (6) bond length outliers are listed below:

| Mol | Chain | Res | Type | Atoms | Z | Observed(Å) | Ideal(Å) |
|---|---|---|---|---|---|---|---|
| 3 | G | 805 | BTI | C2-S1 | -4.00 | 1.76 | 1.82 |
| 3 | G | 803 | BTI | C2-S1 | -3.95 | 1.76 | 1.82 |
| 3 | G | 801 | BTI | C2-S1 | -3.94 | 1.76 | 1.82 |
| 3 | G | 804 | BTI | C2-S1 | -3.86 | 1.76 | 1.82 |
| 3 | G | 806 | BTI | C2-S1 | -3.67 | 1.76 | 1.82 |
| 3 | G | 802 | BTI | C2-S1 | -3.66 | 1.76 | 1.82 |

All (26) bond angle outliers are listed below:

| Mol | Chain | Res | Type | Atoms | Z | Observed($^o$) | Ideal($^o$) |
|---|---|---|---|---|---|---|---|
| 3 | G | 803 | BTI | C4-C2-S1 | 3.05 | 108.11 | 105.20 |
| 3 | G | 805 | BTI | C4-C2-S1 | 3.01 | 108.07 | 105.20 |
| 3 | G | 801 | BTI | C4-C2-S1 | 3.00 | 108.06 | 105.20 |
| 3 | G | 801 | BTI | C6-C5-C4 | 2.88 | 111.16 | 108.66 |
| 3 | G | 803 | BTI | C6-C5-C4 | 2.86 | 111.14 | 108.66 |
| 3 | G | 805 | BTI | C6-C5-C4 | 2.80 | 111.09 | 108.66 |
| 3 | G | 802 | BTI | C6-C5-C4 | 2.78 | 111.07 | 108.66 |
| 3 | G | 806 | BTI | C6-C5-C4 | 2.72 | 111.02 | 108.66 |

*Continued on next page...*

Not For Manuscript Review

*Continued from previous page...*

| Mol | Chain | Res | Type | Atoms | Z | Observed($^o$) | Ideal($^o$) |
|---|---|---|---|---|---|---|---|
| 3 | G | 804 | BTI | C4-C2-S1 | 2.62 | 107.70 | 105.20 |
| 3 | G | 804 | BTI | C6-C5-C4 | 2.56 | 110.88 | 108.66 |
| 3 | G | 801 | BTI | C6-S1-C2 | 2.49 | 95.01 | 89.89 |
| 3 | G | 803 | BTI | C6-S1-C2 | 2.47 | 94.96 | 89.89 |
| 3 | G | 805 | BTI | C6-S1-C2 | 2.46 | 94.94 | 89.89 |
| 3 | G | 806 | BTI | C6-S1-C2 | 2.44 | 94.91 | 89.89 |
| 3 | G | 802 | BTI | C6-S1-C2 | 2.43 | 94.89 | 89.89 |
| 3 | G | 803 | BTI | C5-C6-S1 | 2.41 | 108.37 | 106.31 |
| 3 | G | 804 | BTI | C6-S1-C2 | 2.37 | 94.77 | 89.89 |
| 3 | G | 805 | BTI | C5-C6-S1 | 2.37 | 108.33 | 106.31 |
| 3 | G | 802 | BTI | C4-C2-S1 | 2.33 | 107.42 | 105.20 |
| 3 | G | 801 | BTI | C5-C6-S1 | 2.24 | 108.23 | 106.31 |
| 3 | G | 806 | BTI | C4-C2-S1 | 2.23 | 107.33 | 105.20 |
| 3 | G | 802 | BTI | N2-C3-N3 | 2.15 | 110.78 | 108.76 |
| 3 | G | 806 | BTI | N2-C3-N3 | 2.13 | 110.76 | 108.76 |
| 3 | G | 805 | BTI | N2-C3-N3 | 2.00 | 110.64 | 108.76 |
| 3 | G | 801 | BTI | N2-C3-N3 | 2.00 | 110.64 | 108.76 |
| 3 | G | 803 | BTI | N2-C3-N3 | 2.00 | 110.64 | 108.76 |

There are no chirality outliers.

There are no torsion outliers.

There are no ring outliers.

5 monomers are involved in 13 short contacts:

| Mol | Chain | Res | Type | Clashes | Symm-Clashes |
|---|---|---|---|---|---|
| 3 | G | 802 | BTI | 2 | 0 |
| 3 | G | 803 | BTI | 3 | 0 |
| 3 | G | 801 | BTI | 3 | 0 |
| 3 | G | 806 | BTI | 2 | 0 |
| 3 | G | 805 | BTI | 3 | 0 |

#### 4.7 Other polymers (i)

There are no such residues in this entry.

#### 4.8 Polymer linkage issues (i)

The following chains have linkage breaks:

Not For Manuscript Review

| Mol | Chain | Number of breaks |
|---|---|---|
| 2 | H | 1 |
| 2 | I | 1 |
| 2 | J | 1 |
| 2 | K | 1 |
| 2 | L | 1 |
| 2 | M | 1 |

All chain breaks are listed below:

| Model | Chain | Residue-1 | Atom-1 | Residue-2 | Atom-2 | Distance (Å) |
|---|---|---|---|---|---|---|
| 1 | H | 219:HIS | C | 220:ILE | N | 3.08 |
| 1 | I | 219:HIS | C | 220:ILE | N | 3.08 |
| 1 | J | 219:HIS | C | 220:ILE | N | 3.08 |
| 1 | K | 219:HIS | C | 220:ILE | N | 3.08 |
| 1 | L | 219:HIS | C | 220:ILE | N | 3.08 |
| 1 | M | 219:HIS | C | 220:ILE | N | 3.08 |

Not For Manuscript Review
